# Defining the vascular niche of human adipose tissue across metabolic states

**DOI:** 10.1101/2024.09.22.610444

**Authors:** Ibrahim AlZaim, Mohamed N. Hassan, Maja Schröter, Luca Mannino, Katarina Dragicevic, Marie Balle Sjogaard, Joseph Festa, Lolita Dokshokova, Sophie Weinbrenner, Blanca Tardajos Ayllon, Bettina Hansen, Rikke Kongsgaard Rasmussen, Julie N. Christensen, Olivia Wagman, Ruby Schipper, Min Cai, Wouter Dheedene, Anja Bille Bohn, Jean Farup, Lin Lin, Samuele Soraggi, Anna Dalsgaard Thorsen, Amanda Bæk, Henrik Holm Thomsen, Maximilian von Heesen, Lena-Christin Conradi, Paul Evans, Carolina E. Hagberg, Joerg Heeren, Margo Emont, Evan D. Rosen, Aernout Luttun, Anders Etzerodt, Lucas Massier, Mikael Rydén, Niklas Mejhert, Matthias Blüher, Konstantin Khodosevich, Robert A Fenton, Bilal N. Sheikh, Niels Jessen, Laura P.M.H. de Rooij, Joanna Kalucka

## Abstract

**Introduction:** Adipose tissue homeostasis depends on a healthy vascular network. Vascular malfunction is a hallmark of obesity^1^, and vascular endothelial dysfunction, in particular, accelerates metabolic diseases, including obesity and diabetes. Single-cell transcriptomics studies have mapped the cellular landscape of human white adipose tissue (WAT)^2–8^. However, the vascular niche remains relatively undefined^9^, especially regarding its heterogeneity, function, and role in metabolic disease. To address this gap, we created a single-cell transcriptome atlas of human subcutaneous adipose tissue (SAT), comprising nearly 70,000 vascular cells from 65 individuals. We characterized seven canonical adipose tissue endothelial cell (AdEC) subtypes and identified a distinct heterogenous population, here referred to as sub-AdECs. Sub-AdECs exhibit gene signatures characteristic of multiple cell types, including mesenchymal, adipocytic, and immune, suggesting they possess diverse properties and identities. Through computational analyses and whole-mount imaging, we validated the occurrence of sub-AdECs and show that these cells likely arise through endothelial-mesenchymal transition (EndMT), the modulation of which limits obesity-associated adipose tissue inflammation and fibrosis. Furthermore, we compared the transcriptomes of vascular cells from individuals living with or without obesity and type 2 diabetes and find metabolic disease-associated inflammatory and fibrotic transcriptomic patterns. The atlas and accompanying analyses establish a solid foundation for investigations into the biology of the adipose tissue vascular niche and its contribution to the pathogenesis of metabolic disease.

## Results

### An Integrated Single-Cell Atlas Reveals Vascular Niche Heterogeneity in Human SAT

A comprehensive mapping of the vascular niche within human adipose tissue is still lacking, despite its biological importance^1,9^. Using fluorescence-activated nuclear sorting (**Supplementary Fig. 1a**) followed by single-nucleus RNA sequencing (snRNA-seq), we profiled over 120,000 nuclei from subcutaneous adipose tissue (SAT) biopsies collected from 14 donors representing three distinct metabolic states—including individuals with and without obesity and type 2 diabetes (T2D) (**Supplementary Table 1**). This dataset was then merged with seven publicly available single-cell (sc)RNA-seq and snRNA-seq datasets of human SAT^2–8^ (see *Methods* section), yielding an integrated atlas comprising 329,774 cells from 65 donors (**Extended Data Fig. 1a–f**; **Supplementary Tables 1–4**). By annotating cellular populations based on putative gene signatures^7,8^ we found that over 80% of vascular cells belonged to snRNA-seq datasets, with a major contribution from our *in-house* dataset (over 60%) (**Extended Data Fig. 1g,h; Supplementary Table 5**). We next focused on mapping the heterogeneity of vascular cells, which after subsetting and re-clustering, comprised ten well-defined cell populations: eight AdEC and two mural cell populations (**Fig. 1a,b**; **Extended Data Figure 2a**; **Supplementary Tables 6 and 8)**. Among the endothelial cells, we found three venous AdEC populations expressing the pan-venous marker *ACKR1*^10^: (V1) canonical venous AdECs expressing *TLL1* and *ELOVL7*; (V2) AdECs enriched in leukocyte adhesion molecules (*ICAM1*, *VCAM1*, *SELE*, *SELP*) ^11,12^; and (V3) capillary-venous AdECs expressing *PLVAP*^13^ and *CD74* (**Fig. 1b,c**; **Supplementary Fig. 1b**). We also identified capillary AdECs defined by *RBP7* and *GPIHBP1* expression. These included canonical capillary AdECs (C1; *CADM2*, *BTNL9*) and interferon-activated capillary AdECs (C2; *IFITM2*, *IFI27*), which shared features with V3 (*e.g. CD74*) (**Fig. 1b,d**; **Supplementary Fig. 1c**). Both C1 and C2 expressed lipid-handling genes (*FABP4*, *CD36*), indicating specialized metabolic functions^3^ (**Fig. 1d Supplementary Fig. 1c**; **Supplementary Table 7**). Notably, C2 and V3 displayed interferon-response profiles, with elevated *HLA genes*, *IRF3*, *ISG15*, and JAK-STAT signaling signatures, suggesting roles as non-professional antigen-presenting cells^14^ (**Extended Data Fig. 2b-d**; **Supplementary Table 9**). Supporting this notion, AdECs have previously been shown to express an interferon-related gene signature *in vitro* in response to combined inflammatory and fibrotic stimuli^15^ (**Extended Data Fig. 2e**)

**Fig. 1:**
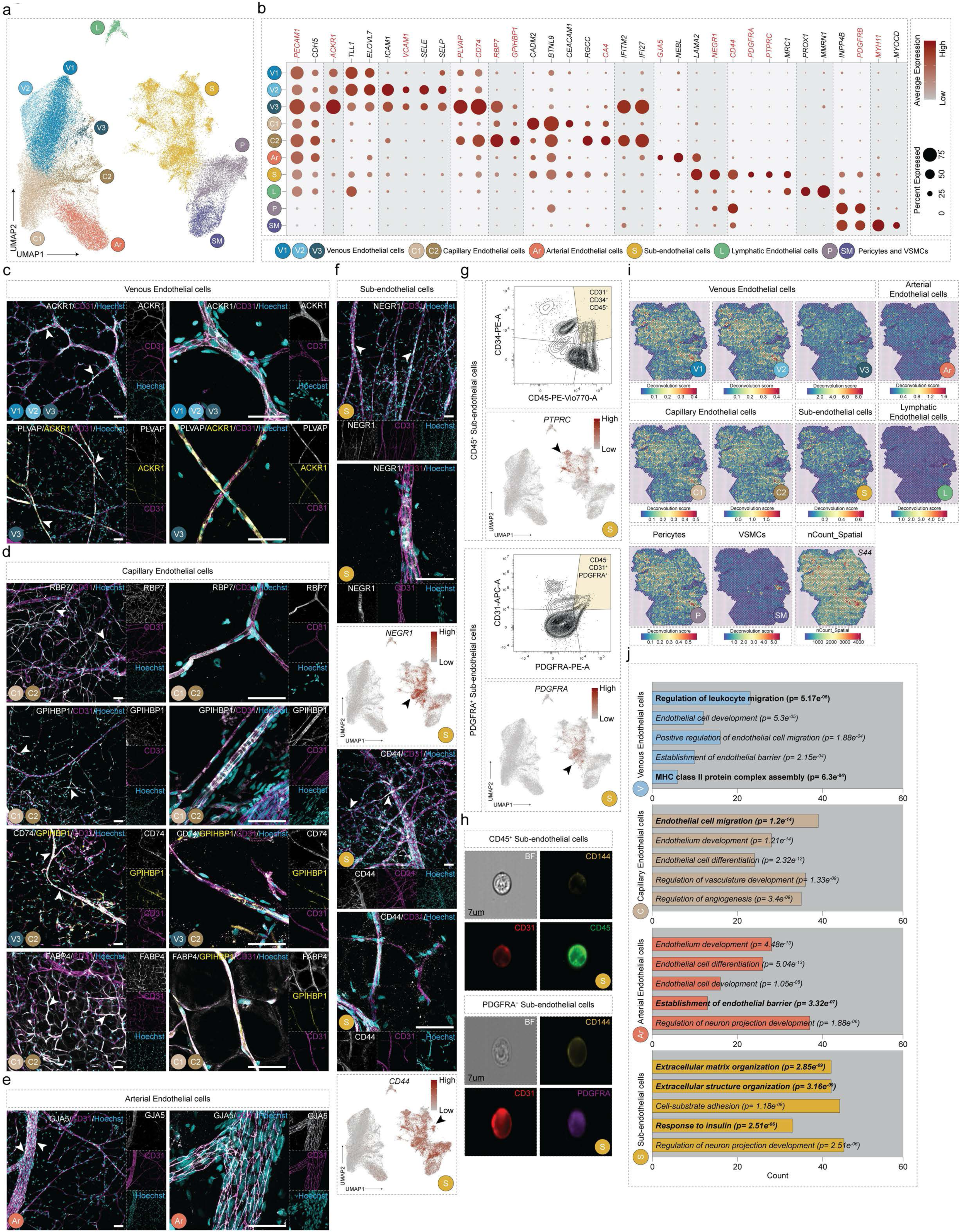
An integrated single-cell atlas of human SAT vascular cells reveals extensive heterogeneity. **a**, UMAP projection of 68,503 vascular cells from human subcutaneous adipose tissue (SAT), identifying ten distinct cell populations. **b**, Dot plot displaying representative marker gene expression across the populations shown in (a). Dot size represents the proportion of cells expressing each gene; color denotes expression level (red: high, grey: low). Genes highlighted in red were validated at the protein level. **c–e,** Immunohistochemical staining of wholemount human adipose tissue for proteins encoded by marker genes in venous (V1–V3; c), capillary (C1–C2; d), and arterial (Ar; e) populations. White arrowheads mark ECs positive for the target protein. **f,** *Top*: Immunohistochemical staining for marker proteins of the sub-AdEC population. *Bottom*: Corresponding feature plots showing gene expression. White arrowheads mark ECs positive for the target protein. Black arrowheads indicate populations with pronounced expression. **g**, *Top*: Representative contour plots from conventional flow cytometry showing sub-AdECs co-expressing the hematopoietic marker CD45 (CD31⁺CD34⁺CD45⁺) or the fibroblastic marker PDGFRA (CD45⁻CD31⁺PDGFRA⁺) in human SAT stromovascular cells. *Bottom*: Corresponding feature plots illustrating gene expression. Black arrowheads indicate populations with pronounced gene expression; color scale denotes expression level (red: high, grey: low). **h,** Representative in-focus images captured by ImageStream showing sub-AdECs in bright field (BF), co-expressing the endothelial proteins CD144/VE-Cadherin (yellow) and CD31 (red), together with either the hematopoietic marker CD45 (CD144⁺CD31⁺CD45⁺; *top*) or the fibroblastic marker PDGFRA (CD144⁺CD31⁺PDGFRA⁺; *bottom*). **i**, Representative sections showing the spatial distribution of deconvolution scores for major vascular cell populations within the human adipose tissue vascular niche, as inferred by *Cell2location*. Color scale represents deconvolution score (red: high, blue: low). **j**, Bar plots showing five representative Gene Ontology (GO) terms among the most significantly enriched pathways in venous, capillary, arterial, and sub-AdEC cell populations. Bolded GO terms have been highlighted in the text. The x-axis indicates the number of associated genes. Scale bars (c-f): 50μm; h: 7μm.

In addition, we identified one arterial AdEC population (*GJA5*, *NEBL*^14^)(**Fig. 1b**; **Supplementary Fig. 1d**), one lymphatic AdLEC population (L; *PROX1*, *MMRN1*), and two mural cell types: pericytes (P; *INPP4B*, *PDGFRB*) and vascular smooth muscle cells (SM; *MYH11*, *MYOCD*) (**Fig. 1b**; **Supplementary Fig. 1e**). Lastly, we defined a heterogeneous population termed sub-AdECs (S), named to reflect their lower expression of pan-endothelial markers, alongside co-expression of fibroblastic and leukocytic genes (*NEGR1*, *CD44*, *PDGFRA*, *PTPRC*, *MRC1*) **(Fig. 1b)**. The presence of sub-AdECs in human SAT was validated using immunohistochemistry (**Fig. 1f**), flow cytometry (**Fig. 1g,h**; **Supplementary Fig. 2a,b**; **Supplementary Table 7**), and was also confirmed in the reanalyzed CD31⁺ SAT fraction from Dhumale P. et al.^16^ (**Supplementary Fig. 2c**; **Supplementary Table 10**). Finally, all vascular subtypes were identified using spatial transcriptomics and showed distinct spatial patterns when mapped onto publicly available data from human SAT^17^ (**Fig. 1i**).

To identify functions specific to each vascular bed, independent of metabolic state, we grouped the AdEC populations by vascular type (e.g., venous: V1–V3; capillary: C1, C2). Using Jaccard similarity analysis, we observed that the sub-AdECs clustered closely with capillary AdECs, suggesting transcriptomic similarity and a possible capillary origin (**Extended Data Fig. 2f**). Moreover, module scoring and gene ontology (GO) analysis revealed distinct signatures: venous AdECs were enriched for genes involved in antigen processing and leukocyte migration, capillary AdECs for endothelial cell migration, and arterial AdECs for endothelial barrier maintenance—mirroring their roles in other tissues (**Fig. 1j**; **Extended Data Fig. 2g-h**; **Supplementary Table 11**). In contrast, sub-AdECs exhibited high expression of genes involved in extracellular matrix (ECM) remodeling and, notably, insulin signaling—features not previously highlighted in adipose endothelial cells (**Fig. 1j**). This suggests a potential role of sub-AdECs in ECM deposition and insulin responsiveness, both of which are critical for adipose tissue homeostasis in obesity and T2D. Enrichment analysis confirmed their matrisome-related gene expression^18^, with higher scores for collagen, glycoprotein, and ECM-affiliated transcripts compared to canonical AdEC populations (**Extended Data Fig. 2g**; **Supplementary Table 8**). Pericytes and vascular smooth muscle cells (SM), which support vascular tone and blood flow, were enriched for genes involved in ECM organization and muscle contraction, consistent with their contractile functions^19^ (**Extended Data Fig. 2h**).

To further explore the regulatory programs driving vascular cell identity, we inferred transcription factor (TF) activity across populations (**Extended Data Fig. 2i,j**; **Supplementary Table 12 and 13**). Canonical AdECs showed high activity of FOXO1, a key endothelial TF promoting vascular stability and maturity/quiescence^20^. Capillary AdECs displayed strong PPARG activity, reflecting their role in lipid metabolism (**Extended Data Fig. 2i,j**). Venous AdECs showed enriched NR2F2 activity, in line with its role in venous specification^21^, while AdLECs demonstrated pronounced PROX1 activity, consistent with their lymphatic identity^18^ (**Extended Data Fig. 2i**). Moreover, sub-AdECs exhibited inferred activities of MAFB and CEBPA— transcription factors associated with endothelial sprouting^22^ and barrier formation^23^ (**Extended Data Fig. 2i**), — as well as stress-responsive TFs such as BATF^24^, NRF1, and IKZF1^25–27^ (**Extended Data Fig. 2i**), suggesting a unique regulatory profile. In mural cells, TF activity analysis revealed enrichment of STAT4 and NFATC2, regulators of muscle cell migration and angiogenic signaling^28,29^, alongside lesser-characterized TFs such as SOX13, TFAP2A, and NR3C1 (glucocorticoid receptor), implicating potential novel regulators of mural cell function.

To assess whether sub-AdECs and other vascular populations are conserved across species, we mapped publicly available single-cell datasets from mouse^7^ and pig^30^ SAT onto our human atlas using scmap or integrated datasets across species (**Extended Data Fig. 3a-e**; **Supplementary Fig. 3a-d**; **Supplementary Tables 2–4, 6, 14- 17**). These comparisons revealed conserved vascular populations—including sub-AdECs—which we further validated in mouse inguinal adipose tissue (**Extended Data Fig. 3f-i**; **Supplementary Fig. 3e,f**, *Methods*). Together, these findings highlight the unexpectedly rich transcriptomic and functional heterogeneity within the human adipose vascular niche—features that are mirrored in mammalian systems.

### Disease-Associated Transcriptomic Alterations in the Adipose Tissue Vascular Niche

Vascular deterioration is a hallmark of obesity and T2D, and AdECs appear particularly susceptible to this dysfunction^1,31^. Although our SAT atlas showed no significant difference in the overall abundance of AdECs across BMI groups, we observed a significant negative correlation between BMI and mural cell abundance (**Extended Data Fig. 4a-d**). Furthermore, differential gene expression and GO analyses uncovered both shared and metabolic state-specific alterations in AdECs and mural cells (**Fig. 2a-c**; **Extended Data Fig. 4e,f**; **Extended Data Fig. 5a**; **Supplementary Table 18,19**). For example, venous AdECs in obesity and obesity-associated diabetic states upregulated the expression of *VCAN*, which encodes versican, a major ECM component that modulates matrix structure and promotes immune cell adhesion and migration^32–34^, and also exhibited changes in the expression of the leukocyte adhesion molecule *VCAM1*^35^ (**Fig. 2d**; **Extended Data Fig. 5b**). In V1 AdECs from donors with obesity and T2D, we observed enrichment of pathways related to lipid handling, alongside downregulation of genes involved in cellular responses to hypoxia and reactive oxygen species (**Fig. 2b; Extended Data Fig. 4f)**. The V2 population exhibited decreased expression of genes regulating immune cell extravasation (**Fig. 2b and Supplementary Table 19**). Meanwhile, arterial AdECs showed reduced expression of genes associated with the negative regulation of vascular permeability (**Fig. 2b**; **Extended Data Fig. 4f**; **Supplementary Table 19)**. Together, these findings suggest that obesity and T2D may impair vascular control of immune cell trafficking, potentially contributing to adipose tissue inflammation.

**Fig. 2.**
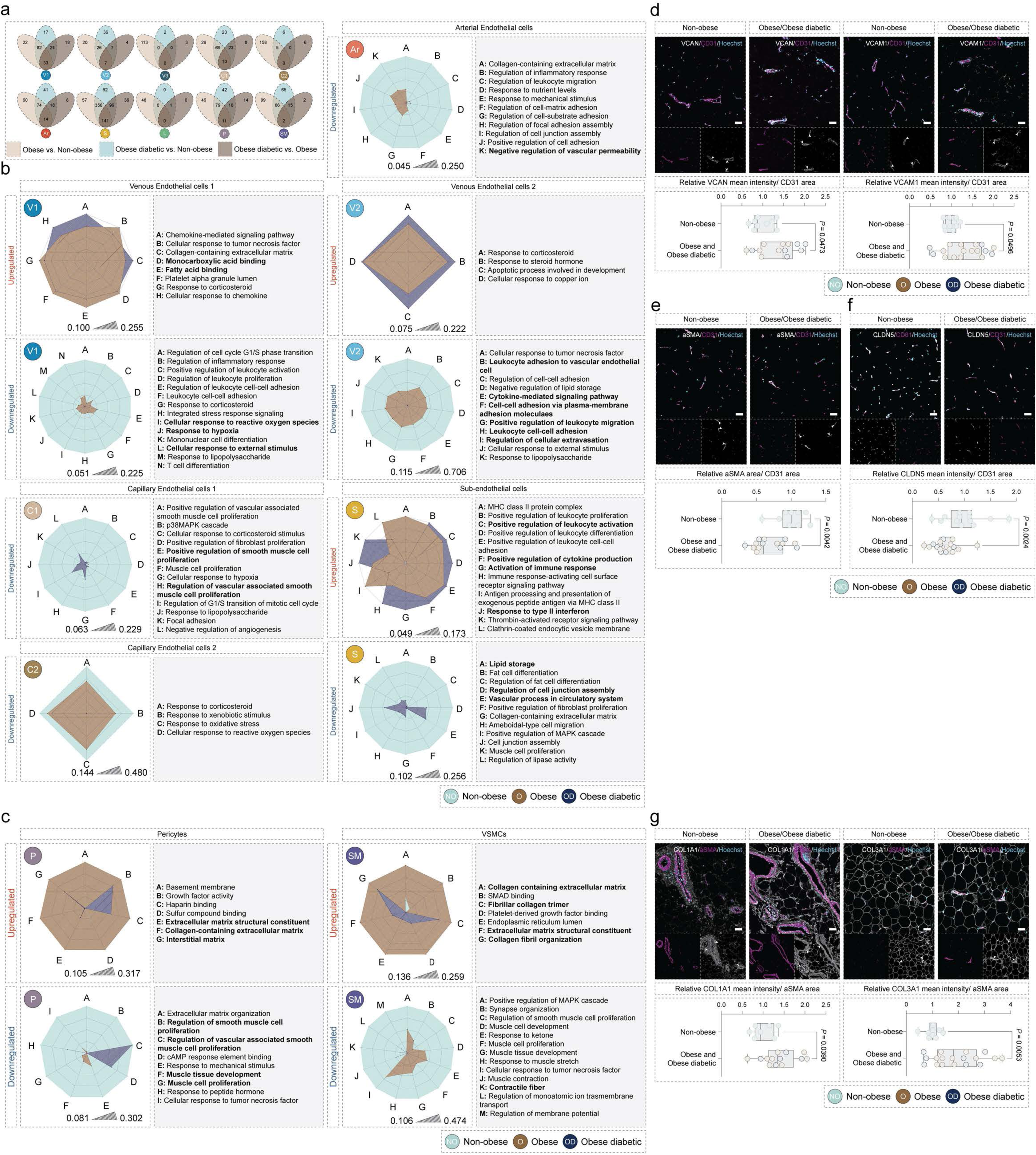
Adipose tissue vascular cells exhibit transcriptomic alterations associated with metabolic disease. **a,** Venn diagrams showing the overlap of significantly differentially expressed genes (DEGs) in each vascular cell population across three comparisons: Non-obese vs. Obese, Non-obese vs. Obese diabetic, and Obese diabetic vs. Obese. **b,** Radar plots illustrating shared upregulated and downregulated GO terms in canonical AdEC and sub-AdEC populations across the Non-obese vs. Obese and Non-obese vs. Obese diabetic comparisons. Scale bars represent the average area under the curve (AUC) for each GO term in a given cell population. **c,** Radar plots showing commonly upregulated and downregulated GO terms in mural cell populations across the same comparisons as in (b). AUC scale bars as in (b). **d,** *Top*: Representative images of immunohistochemical staining of human adipose tissue sections showing deregulated proteins in the endothelial compartment, including versican (VCAN) and vascular cell adhesion molecule 1 (VCAM1). *Bottom*: Quantification of VCAN (Non-obese, n=9; Obese, n=8; Obese diabetic, n=5) and VCAM1 (Non-obese, n=10; Obese, n=12; Obese diabetic, n= 5) staining in the endothelial compartment (CD31 positive). **e,** *Top*: Representative images of immunohistochemical staining for mural cell coverage in human adipose tissue sections, assessed by aSMA⁺/CD31⁺ area, across Non-obese (n=9), Obese (n=8), and Obese diabetic (n=6) individuals. *Bottom*: Quantification of mural coverage based on aSMA⁺/CD31⁺ area. **f**, *Top*: Representative images of immunohistochemical staining for the tight junction protein claudin-5 (CLDN5) in human adipose tissue sections. *Bottom*: Quantification of CLDN5 staining in the endothelial compartment (Non-obese, n=9; Obese, n=8; Obese diabetic, n=6). **g**, *Top*: Representative images of immunohistochemical staining for extracellular matrix proteins collagen I (COL1A1) and collagen III (COL3A1) in the mural compartment. *Bottom*: Quantification of COL1A1 (Non-obese, n=7; Obese, n=8; Obese diabetic, n=5) and COL3A1 (Non-obese, n=10; Obese, n=11; Obese diabetic, n=7) staining. In quantifications, Obese and Obese diabetic donors were pooled where indicated, as the deregulated genes exhibited changes in the same direction and of comparable magnitude. All scale bars: 50μm. Statistics: In box plots (d-g), error bars represent mean ± s.e.m. *P* values were calculated using unpaired *t*-tests. *P* < 0.05 was considered statistically significant.

C1 AdECs from donors with obesity and T2D downregulated genes associated with promoting VSMC proliferation, suggesting a potential reduction in mural cell coverage of the microvasculature^36,37^ **(Fig. 2b**; **Extended Data Fig. 4f**; **Supplementary Table 11**). This was supported by quantitative analysis of tissue sections, showing a reduced ratio of α-SMA-positive area to CD31-positive area, consistent with fewer morphologically mature vessels (**Fig. 2e**). Interestingly, we observed an upregulation of *CLDN5* expression at the transcript level across the endothelial compartment in obesity, which could reflect a compensatory mechanism aimed at preserving vascular barrier integrity and limiting immune cell extravasation (**Extended Data Fig. 5c**). However, this contrasts with our protein-level data, which show a downregulation of claudin (**Fig. 2f**), consistent with previous reports of obesity-associated claudin switching and endothelial barrier disruption in different organs^38,39^. This discrepancy suggests post-transcriptional regulation or increased protein turnover in the context of metabolic stress. GO analysis further revealed that sub-AdECs are enriched for pathways related to cytokine responses, antigen presentation, and leukocyte-mediated inflammation in obesity and T2D, suggesting that these cells may actively participate in immune regulation (**Fig. 2b**; **Extended Data Fig. 4f**; **Supplementary Table 19**). At the same time, sub-AdECs displayed reduced expression of genes involved in cellular junctions and vascular processes, further supporting their association with endothelial dysfunction in metabolic disease.

Pericytes and VSMCs showed reduced expression of genes related to cell proliferation and contractile function, alongside increased expression of genes involved in ECM deposition, including *COL1A1* and *COL3A1* **Extended Data Fig. 5a,d)**. They also upregulated pathways responsive to pro-fibrotic stimuli, consistent with a shift toward a pro-fibrotic VSMC phenotype, similar to that observed in atherosclerosis^40^ **(Fig. 2c**; **Extended Data Fig. 5a**; **Supplementary Table 11**). Transcript-level changes were supported by increased COL1A1 and COL3A1 protein levels in mural cells, indicating enhanced fibrosis under obesogenic conditions (**Figure 2g**; **Extended Data Fig. 5e**).

We next probed for altered TF activities in metabolic disease and observed in V1, C1, and arterial AdECs in donors living with obesity and T2D, an increased inferred activity of SREBP-1 (encoded by *SREBF1*), which is activated in the vascular endothelium in response to disturbed flow^41^ and which upregulates the expression of lipogenic enzymes in response to metabolic changes^42^ (**Extended Data Fig. 5f**; **Supplementary Table 20**). V1, V2, and C2 AdECs exhibited increased inferred activity of TEAD4, which links endothelial nutrient acquisition to angiogenesis^43^. Notably, TEAD4 also plays a role in mechanosensing via YAP/TAZ, suggesting a possible response to altered extracellular stiffness in disease^44^. Furthermore, sub-AdECs of donors living with obesity and T2D showed increased activity of RBPJ, a transcriptional regulator of Notch signaling^45^, BATF (an immune-responsive TF component of the pro-inflammatory AP1 complex^24^), and GABPA (previously predicted to play a pro-angiogenic role in ECs^22^). Notably, in V1 and sub-AdECs, the inferred activity of SRF, which is important for sprouting angiogenesis and the maintenance of vessel integrity^46,47^, was downregulated (**Extended Data Fig. 5f**). Collectively, these data reveal an overall dysfunction of the adipose tissue vascular niche, possibly underpinning a key role in the pathogenesis of metabolic disease.

### Sub-endothelial cells exhibit signatures to modulate inflammation and fibrosis

In our human SAT atlas, sub-AdECs accounted for nearly one-third of all AdECs and showed heterogeneous expression of marker genes typically associated with other cell types, such as *PTPRC*, *PDGFRA*, and *NEGR1* **(Fig. 1a,b**; **Supplementary Table 7)**. To explore their diversity in more detail, we subset and re-clustered the sub-AdEC population, identifying seven subpopulations (**Fig. 3a,b**; **Extended Data Fig. 6a,b**; **Supplementary Tables 21 and 22**). These subpopulations expressed pan-endothelial markers, though at lower levels than canonical AdECs (**Fig. 1b**), as well as mesenchymal markers such as *CD44* and *ZEB2*, consistent with a hybrid <u>end</u>othelial to <u>m</u>esenchymal transition (EndMT) phenotype^48^ (**Extended Data Fig. 6c**). By deconvoluting marker genes of various cell types onto publicly available spatial transcriptomics data, we observed a strong spot-wise correlation between canonical and sub-AdEC scores (**Fig. 3c**; highlighted with a black box). We next calculated an EC score for each spot using a curated set of pan-EC genes (see: *Methods,* **Supplementary Table 23**) and found that sub-AdEC subpopulations predominantly deconvoluted onto spots with relatively high EC scores (**Fig. 3d**). This spatial association supports the endothelial identity of sub-AdECs and their localization within the vascular niche.

**Fig. 3:**
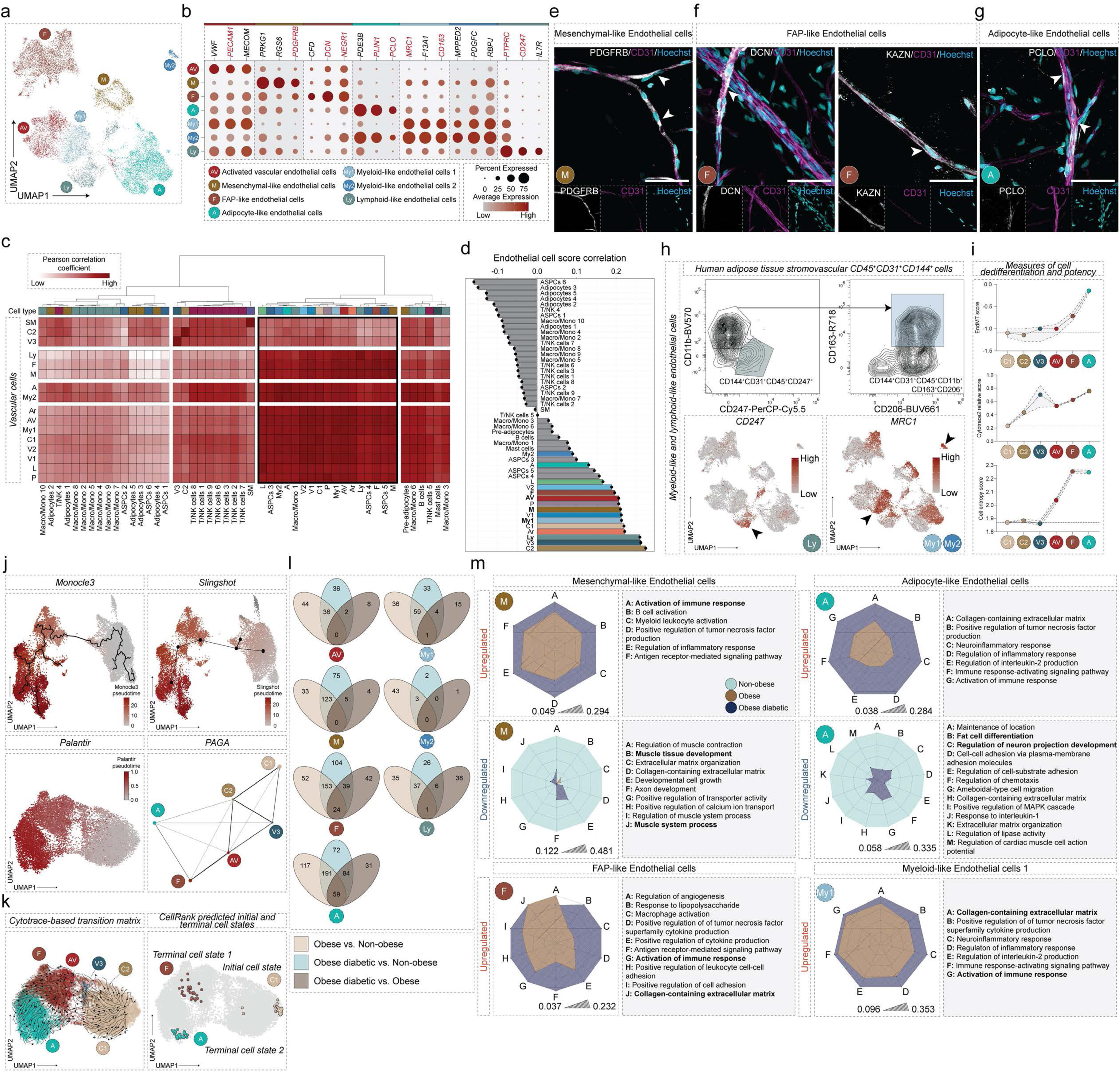
Sub-endothelial cells exhibit transcriptional signatures associated with inflammation and fibrosis in metabolic disease. **a**, UMAP projection of 16,658 sub-AdECs from the human SAT atlas, revealing seven distinct subpopulations. **b**, Dot plot showing expression of representative marker genes across sub-AdEC subpopulations. Dot size reflects the percentage of cells within each cluster expressing a given gene; color scale indicates expression level (red: high, grey: low). Genes highlighted in red were validated at the protein level. **c**, Heatmap showing average spot-wise Pearson correlation coefficients (n=10) between cell type-specific deconvolution scores of vascular (*left*) and all identified cell types (*bottom*). **d**, Bar plot displaying average spot-wise correlation between a pre-computed EC score and the cell type-specific deconvolution scores (n=10). **e–g**, Representative immunohistochemical staining of wholemount human adipose tissue for proteins encoded by key marker genes of sub-AdEC subpopulations: PDGFRB (Mensynchymal-like ECs (M), **e**), DCN and KAZN (FAP-like ECs (F), **f**), and PCLO (Adipocyte-like ECs (A), **g**). **h**, Myeloid-like (My1, My2) and lymphoid-like EC (Ly): *Top:* Representative contour plots from full-spectrum flow cytometry showing CD144/VE-Cadherin⁺CD31⁺CD45⁺CD247⁺ (*left*) and CD144/VE-Cadherin⁺CD31⁺CD45⁺CD11b⁺CD163⁺CD206⁺ (*right*) sub-AdEC populations within the SVF of human SAT. *Bottom*: Corresponding feature plots showing gene expression of *CD247* and *MRC1* (encoding CD206) in sub-AdEC subpopulations. Arrows highlight populations with pronounced expression. Color scale indicates gene expression level (red: high, grey: low). **i**, Median measures of cell dedifferentiation and potency (EndMT score, *Cytotrace2* relative score, and Cell entropy score calculated using *SLICE*) across select endothelial cell populations. **j**, Pseudotime trajectory inference showing the directionality of cell differentiation inferred using *Monocle3*, *Slingshot*, and *Palantir* along with RNA velocity-based cell type connectivity inferred by *PAGA.* **k**, *Cytotrace*-based transition matrix, and *CellRank* unsupervised prediction of initial and terminal cell states using the cell embedding from *Palantir*. **l**, Venn diagrams showing the overlap of significantly differentially expressed genes in each sub-AdECs subpopulation in the three comparisons (Non-obese vs. Obese, Non-obese vs. Obese diabetic, and Obese diabetic vs. Obese). **m**, Radar plots presenting common upregulated and downregulated GO terms in sub-AdECs subpopulations in the Obese vs. Non-obese and Obese diabetic vs. Non-obese comparisons. Bolded GO terms have been highlighted in the text. Scale bars (e-g): 50μm; Scale bars (m) Average area under curve.

Sub-AdEC subpopulations included **<u>a</u>**ctivated **<u>v</u>**ascular AdECs (AV; expressing *VWF* and *MECOM*) and **<u>m</u>**esenchymal-like AdECs (M; expressing *RGS6* and *PDGFRB,* **Fig. 3b,e**; **Supplementary Fig. 4a**). We also identified **f**ibro-**<u>a</u>**dipogenic **<u>p</u>**rogenitor (also known as adipose stromal and progenitor cells; ASPCs^49^) (FAP)-like AdECs, (expressing *DCN* and *KAZN,* **Fig. 3b,f**; **Supplementary Fig. 4b**). Finally, adipocyte-like AdECs (A) were identified by expression of *PLIN1* and *PCLO* (**Fig. 3b,g**; **Supplementary Fig. 4c**). While the adipogenic potential of ECs *in vivo* remains debated^8,50–52^, we employed a vascularized adipose spheroid model^53^, which recapitulates the unilocular morphology and functional characteristics of human adipocytes and provides a vascular niche optimized to maintain EC identity and viability. This allowed us to test whether ECs could acquire adipogenic traits under defined differentiation conditions. After 10 days of adipogenic induction, ECs accumulated PLIN1-coated lipid droplets *in vitro*, supporting their potential to adopt adipocyte-like features (**Extended Data Fig. 6d**). Similarly, the heterogeneity within the sub-AdECs was also found in murine adipose tissue (**Supplementary Fig. 4d**).

Additional subpopulations included two **<u>my</u>**eloid-like AdEC populations (My1 and My2; expressing *MRC1* and *CD163*) and one **<u>ly</u>**mphoid-like AdEC population (Ly; expressing *IL7R* and *CD247*, **Fig. 3b,h**; **Supplementary Fig. 4e)**, pointing to a potential immunomodulatory role^54^. A similar immune cell–like endothelial population expressing *TYROBP* and *FCER1G* was previously identified in an integrated single-cell atlas of human WAT^8^. Furthermore, endothelial cells undergoing EndMT followed by acquisition of a macrophage-like transcriptome have been described in murine carotid arteries exposed to disturbed flow^48^.

To explore the broader functional identity of sub-AdECs prior to dissecting the heterogeneity of individual subclusters, we first examined their overall differentiation state and potential lineage relationships. Given that sub-AdECs exhibited a gene expression signature characteristic of capillary AdECs (**Extended Data Fig. 6e**), we investigated whether FAP-like, adipocyte-like, and activated vascular AdECs might originate from canonical capillary AdECs using pseudotime trajectory inference (see: *Methods*). As a first step, we computed EndMT, cell potency, and cell entropy scores, which revealed that sub-AdECs display an endothelial– mesenchymal hybrid state (**Fig. 3i**; **Extended Data Fig. 6f**). We then applied complementary tools (*Monocle3, Slingshot, PAGA, and Palantir*) to model pseudotime trajectories (**Fig. 3j**; **Extended Data Fig. 6f)**. These analyses identified two terminal trajectories emerging from canonical AdECs via an intermediate activated vascular AdEC state: FAP-like and adipocyte-like AdECs. *CellRank* analysis further supported these differentiation paths (**Fig. 3k**). This pattern is consistent with previous studies in which adipogenically primed precursor cells diverged into fibrogenic and adipogenic fates^55,56^, suggesting that sub-AdECs may arise from canonical endothelial populations through partial EndMT. In line with this transition, we observed a gradual increase in the expression of TFs implicated in EndMT, including *ZEB2* and *TWIST1*, alongside a corresponding decrease in the endothelial transcription factor *ERG* (**Extended Data Fig. 6g**). *In silico* knockout of *TWIST1,* of which gene regulatory network is available, completely abolished the pseudotime progression, underscoring its potential role in driving this transition (**Extended Data Fig. 6h**).

To experimentally validate these findings, we exposed primary human AdECs to pro-inflammatory and pro-fibrotic stimuli known to induce EndMT^57^. This led to a pronounced downregulation of endothelial marker genes and a robust induction of mesenchymal gene expression (**Extended Data Fig. 6i**), supporting snRNA-seq results. At the protein level, we observed increased expression of SNAIL, CD44, TAGLN, and VCAM1 (**Extended Data Fig. 6j**), along with loss of VE-Cadherin expression at day 7 (**Extended Data Fig. 6j,k**) and increased nuclear translocation of ZEB2 (**Extended Data Fig. 6k**). Functionally, these changes were accompanied by enhanced immune cell adhesion and increased migratory capacity *in vitro* (**Extended Data Fig. 6l**). Together, these findings suggest that AdECs are primed to activate gene programs typically inactive in ECs, reinforcing their potential for phenotypic plasticity and adaptation within a diseased microenvironment.

### Functional Characterisation of EndMT Regulators in Adipose Tissue Vascular Remodelling

To investigate regulators of sub-AdEC emergence *in vivo*, we focused on ZEB2 and TWIST1, both of which are strongly implicated in endothelial–mesenchymal plasticity^58^. Our rationale was strengthened by three main findings: (i) *in silico Twist1* knockout disrupting pseudotime progression from capillary AdECs to subtypes; (ii) high expression of *ZEB2* in sub-AdECs and ZEB2 nuclear translocation under pro-EndMT conditions *in vitro*; and (iii) identification of sub-AdEC-like populations in lineage-traced murine adipose tissue^59^ (**Supplementary Fig. 4d**; **Supplementary Fig. 5**; **Supplementary Table 24**). To assess their roles under metabolic stress, we analyzed archived tissue from endothelial-specific Zeb2 knockout (*Zeb2*^ΔEC^) mice^60^ and prospectively collected tissue from endothelial-specific Twist1 knockout (*Twist1*^ΔEC^) mice^61^, both fed obesogenic diets (**Extended Data Fig. 7a, f**). Although adipocyte size remained unchanged in *Zeb2*^ΔEC^ and *Twist1*^ΔEC^ mice compared to their littermate controls (**Extended Data Fig. 7b, g**), both models exhibited a marked reduction in fibrosis, as assessed by Picrosirius red staining (**Extended Data Fig. 7c, h**). This reduction in fibrotic remodeling was accompanied by decreased immune cell infiltration, measured by F4/80 staining (**Extended Data Fig. 7d, i**). Notably, the vascular rarefaction typically induced by obesity was attenuated in both knockout models. This was accompanied by preserved, or even more pronounced VE-Cadherin staining, particularly evident in *Twist1*^ΔEC^ animals, indicative maintained vascular integrity (**Extended Data Fig. 7e, j**). Together, these findings indicate that ZEB2 and TWIST1 are involved in endothelial remodeling in adipose tissue during obesity, and that their loss preserves vascular integrity under metabolic stress.

To investigate the potential roles of sub-AdEC subpopulations in adipose tissue biology during obesity and T2D, we performed differential gene expression analysis, GO analysis, gene set variation analysis (GSVA), and matrisome signature scoring^18^. These analyses revealed both shared and distinct transcriptional programs across metabolic states (**Fig. 3l)**. Mesenchymal-like, adipocyte-like and myeloid-like subpopulations were enriched for immune-related signatures, while activated vascular and FAP-like sub-AdECs were associated with ECM remodeling (**Fig. 3m**; **Extended Data Fig. 8a**–d; **Supplementary Tables 25–26**).

To gain insight into upstream regulators, we inferred TF activity both in a metabolic state-independent manner and across disease states (**Extended Data Fig. 9a**–c; **Supplementary Tables 27-29)**. Inferred activity of canonical endothelial TFs—ERG, ETV6, FOXO1, ELF1, and ELF2—was consistently detected across all sub-AdEC subpopulations, supporting their endothelial origin. Under obese and diabetic conditions, we observed increased activity of SPI1, a known driver of EndMT^62^, and RUNX2, which promotes endothelial proliferation and migration in metabolically stressed environments^63,64^. Notably, inferred ZEB2 activity was reduced across all sub- AdEC subsets in metabolic disease, possibly reflecting a transition toward a later or stabilized EndMT state in which ZEB2 expression is no longer maintained (**Extended Data Fig. 9c**).

### Intercellular communication and valuable drug targets within the vascular niche

Accumulating evidence suggests that adipose vascular cells engage in both, homeostatic and pathological intercellular communication^7,45,65,66^. To investigate the extent of intercellular communication implicating the vascular niche, we sub-clustered the major cell types comprising the atlas at high granularity (**Supplementary Fig. 6a-e**; **Supplementary tables 30)** and computed differential connectomics using *CellChat*^67^. This analysis revealed an increased number and strength of cell–cell interactions in obesity, but a reduction in both metrics in obesity compounded by T2D, relative to the non-obese state (**Fig. 4a**). These interactions were primarily mediated by adipocytes, adipose stem and progenitor cells (ASPCs), and adipose endothelial cells—particularly sub-AdEC subpopulations (**Fig. 4b,c**; **Extended Data Fig. 10a,b**). Intercellular communication inferred using *LIANA^+^*^68^ corroborated the findings from *CellChat* (**Fig. 4c**; **Extended Data Fig. 10b**; **Supplementary Table 31**) and aligned with previously published studies^7^. These altered intercellular interactions were either conserved across metabolic states or showed context-specific modulation (**Supplementary Fig. 6f**). Notably, pro-angiogenic signaling via vascular endothelial growth factor (VEGF)^69,70^ was diminished in obesity compounded by T2D, while transforming growth factor beta (TGF-β)^70^ signaling appeared to be specific to obesity. In contrast, bone morphogenetic protein (BMP) signaling was elevated in both obesity and obesity with T2D (**Supplementary Fig. 6f**). These findings suggest that the adipose endothelium is highly responsive to angiogenic cues in a metabolic state–dependent manner.

**Fig. 4:**
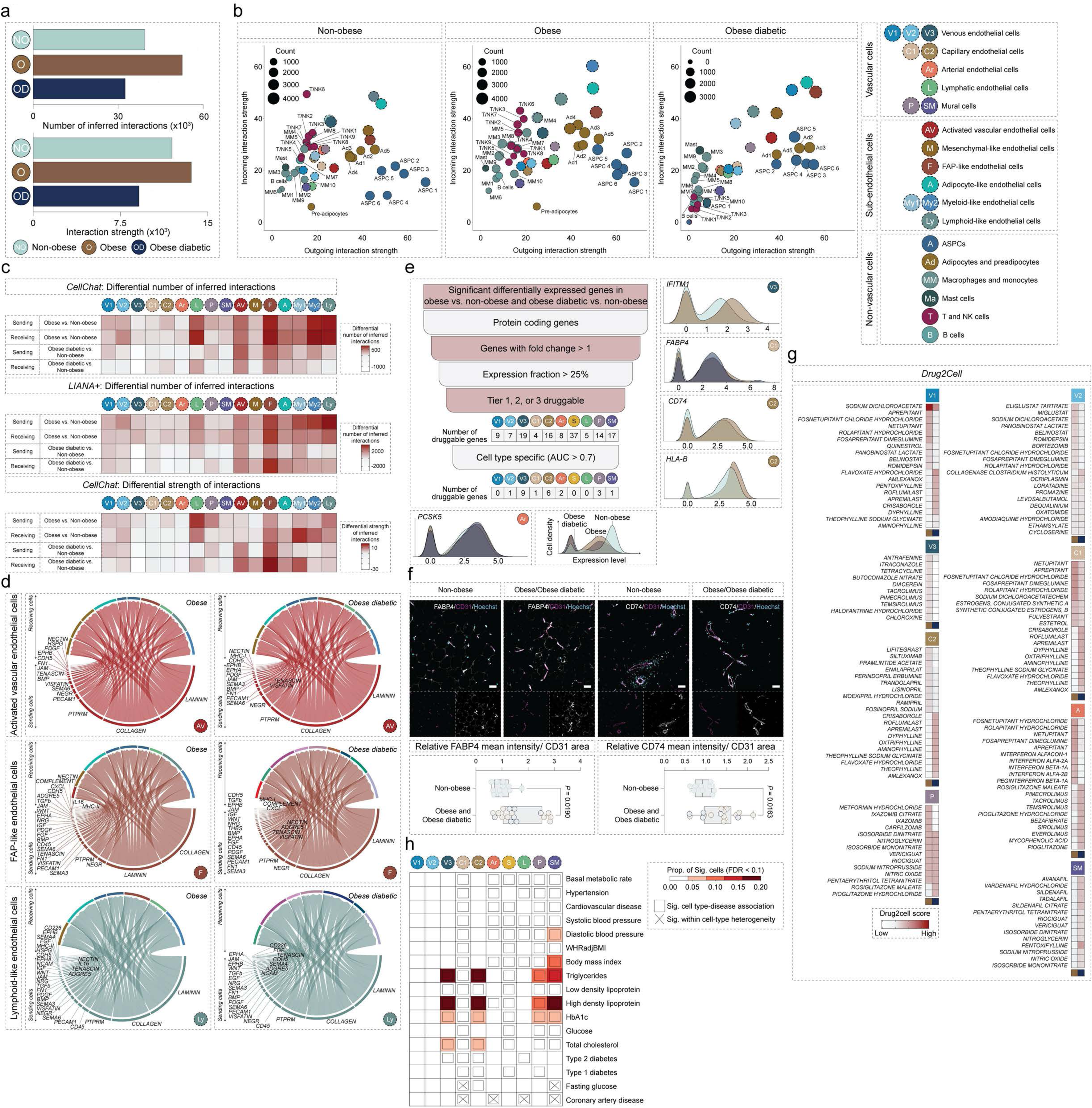
Vascular cell populations are implicated in intercellular communication in metabolic health and disease. **a**, Bar plots depicting the number of intercellular interactions and their strength in each metabolic state. **b**, Scatter plots showing the strength of the outgoing (x-axis) and the incoming (y-axis) interaction strength for each cell type in each metabolic state. **c**, Differential number of interactions implicating the vascular cell compartment inferred by *CellChat* and *LIANA^+^* and the differential strength of interactions inferred by *CellChat*. **d**, Chord diagrams depicting the results of *CellChat* inferred ligand–receptor analysis for significant signaling pathways between the three sub-AdECs populations with the highest differential number and strength of interactions (AV, F, and Ly) and their main interacting partners (receiving cells) in obesity and obesity compounded by T2D. **e**, Filtered druggable differentially expressed protein-coding targets in the vascular cells of human SAT in each metabolic state. Ridge plots represent the expression level of specific genes in vascular populations as a factor of cell density (frequency of cells). **f**, Representative immunohistochemical images validating the upregulation of the two druggable targets FABP4 and CD74, at the protein level, in the adipose tissue of donors living with obesity or obesity compounded by T2D. Quantification of immunohistochemical stainings of FABP4 (Non-obese; n=11, Obese; n=10, and Obese diabetic; n=7) and CD74 (Non-obese; n=9, Obese; n=12, and Obese diabetic; n=5) in the endothelial cell compartment. Scale bar: 50μm. **g**, Heatmaps depicting the differential enrichment of *Drug2cell* scores of drugs identified to target cell populations within the vascular niche in donors living with obesity or obesity compounded by T2D. **h**, Heatmap depicting *scDRS* calculated significant associations between the vascular cell types of human SAT vascular niche and seventeen relevant metabolic and cardiovascular traits. In quantifications, Obese and Obese diabetic donors were pooled where indicated, as the deregulated genes exhibited changes in the same direction and of comparable magnitude. In box plots, error bars represent ± s.e.m and *P* values were calculated using an unpaired t-test. *P* < 0.05 was considered statistically significant.

To study intercellular communication specifically involving sub-AdEC subpopulations, we performed ligand–receptor analysis, focusing on the three subsets with the most pronounced differential interaction profiles: activated vascular (AV), FAP-like (F), and lymphoid-like (Ly) sub-AdECs, along with their primary interacting cell types (**Fig. 4d**). This analysis revealed pronounced signaling through the collagen and laminin pathways, among others, possibly implicating a potential role for sub-AdECs in driving tissue fibrosis (**Fig. 4d**; **Extended Data Fig. 10c**; **Supplementary Table 32).** These findings were supported by *NicheNet*^71^, which identified activation of key inflammatory and fibrotic signaling programs in the corresponding receiver cells (**Supplementary Fig. 7**,8), further supporting previous reports implicating EndMT in fibrotic and inflammatory responses^58^.

Adding to the translational impact of our SAT vascular cell atlas, we investigated whether differentially expressed protein-coding genes represented potential drug targets in obesity and T2D. This analysis identified *IFITM1* in the V3 population, *FABP4*^72^ in the C1 population, *CD74*^73–76^ and *HLA-B*^77^ in the C2 population, and *PCSK5*^78^ in arterial AdECs, among others, as potentially promising druggable targets (**Fig. 4e**; **Supplementary Table 33)**. Indeed, endothelial expression of FABP4 and CD74 was markedly elevated in human adipose tissue sections from individuals with obesity and T2D (**Fig. 4f**; **Extended Data Fig. 10d**). Complementarily, we used *Drug2Cell*^79^ and GREP (Genome for REPositioning drugs)^80^ to infer possible drug repurposing to target the vascular niche in metabolic disease (**Fig. 4g**; **Supplementary Table. 34,35**). Drug targets identified included modulators of EC permeability, angiogenesis, and immune cell interactions, such as phosphodiesterase 3 (PDE3) and PDE4^81,82^ (e.g. the xanthine drug family, including Dyphylline) and the neurokinin 1 receptor^83–85^ (e.g. Netupitant). In contrast, drug targets for the mural cell compartment were largely dominated by activators of soluble guanylate cyclase and inhibitors of PDE5A (e.g. Sodium Nitroprusside) (**Fig. 4g**). Moreover, we investigated the enrichment of polygenic disease-associated genes implicated by gene-wide association studies in the vascular cell populations using *scDRS*^86^. This approach associated vascular populations with traits pertaining to metabolic and cardiovascular disease such as measures of cholesterol and glucose homeostasis (**Fig. 4h**). Collectively, these analyses support the notion that certain AdEC populations are highly relevant to metabolic disease, displaying altered intercellular interactions and expressing druggable gene products that may be targeted therapeutically.

## Discussion

Vascular health is central to adipose tissue homeostasis, yet the complexity of the adipose vasculature and its maladaptation in obesity and T2D remains poorly understood. By constructing an integrated atlas of the human SAT vascular niche, we comprehensively characterized the transcriptomic heterogeneity of vascular cells and their functional roles in metabolic health and disease, revealing diverse endothelial and mural cell populations, their remodeling in disease, and potential targets for therapeutic intervention.

We identify and characterize a transcriptionally diverse endothelial landscape, including classical venous, capillary, arterial, and lymphatic subsets, as well as a previously unappreciated population of sub-AdECs. These cells are abundant in SAT and display a hybrid transcriptional identity that bridges endothelial, fibroblastic, and immune ontologies. Interestingly, these cells have been noted in recently published datasets; however, a clear description and comprehensive profiling have been lacking^13,87^. Here, we provide both computational and experimental evidence that sub-AdECs likely emerge through partial EndMT and retain a degree of plasticity that may underlie their context-specific functions. We further demonstrate the presence of sub-AdECs across mammalian systems and show that their possible emergence via EndMT contributes to tissue fibrosis and inflammation. Inflammatory signaling is increasingly recognized as an intrinsic feature of endothelial plasticity, including EndMT^88^, however, whether the inflammatory milieu of obese adipose tissue actively drives pathological EndMT remains to be fully elucidated. Notably, we identified sub-AdEC subpopulations expressing hematopoietic genes, including CD45, whose expression in endothelial cells has been shown to be sufficient to induce EndMT^89^ and promote vascular wall remodeling in a mouse model of carotid artery ligation^90^. We also identified sub-AdEC subpopulations expressing adipocytic and fibroblastic gene signatures, further supporting previous reports of adipocyte-like and immune-like endothelial states^8,51,55,91^. Consistently, both our study and previous reports have found these cells to be equally abundant in non-obese individuals and those with obesity and T2D^87^. This suggests that sub-AdEC subpopulations may exert context-dependent functions—either homeostatic or pathological, which require further investigation. Leveraging computational tools to infer intercellular communication, we identify profound alterations in signaling networks in obesity and T2D, with sub-AdECs emerging as key signaling hubs. Using multiple ligand–receptor inference algorithms, we uncover enriched collagen and laminin signaling from sub-AdECs to adipose stromal cells, consistent with their potential role in driving fibrotic remodeling. At the same time, classical pro-angiogenic signals, such as VEGF, are diminished in T2D, suggesting a failure to initiate effective vascular regeneration^92^. Recognizing the therapeutic relevance of these alterations, we next assessed the druggability of endothelial subtypes. Several populations express targets such as *FABP4* and *CD74* which are upregulated in metabolic disease and associated with vascular dysfunction. Through computational drug repurposing and polygenic trait enrichment analysis, we identify candidate interventions aimed at preserving vascular integrity, modulating immune activation, and limiting ECM remodeling.

Together, our findings outline a transcriptional and functional roadmap of adipose tissue vascular remodeling in metabolic disease. The discovery of plastic, immune-responsive endothelial populations and conserved regulatory circuits offers insight into the persistence of vascular dysfunction in obesity and diabetes. More broadly, this atlas provides a foundation for exploring the adipose vascular niche and its therapeutic targeting in metabolic disorders.

## Methods

### Study cohort and clinical sample acquisition

Human deep subcutaneous adipose tissue samples were obtained from non-obese donors undergoing hemicolectomy, laparoscopic transabdominal preperitoneal approach, fundoplication with mesh hiatoplasty, or band-reinforced gastric bypass and donors with obesity undergoing elective cosmetic abdominoplasty, gastric sleeve resection, or gastric bypass surgeries at the Clinic for General, Visceral, and Pediatric Surgery of the University Medical Center Göttingen in Göttingen, Germany. Given the wide range of obesity and T2D phenotypes, we used all the available clinical parameters to classify the donors. Obesity was defined based on body mass index (BMI) (kg/m^2^): Non-obese donors (BMI < 30) and donors with obesity (BMI > 30). Donors were classified as having type two diabetes (T2D) based on the presence of a previous history of T2D and antidiabetic medication. This yielded three groups in total matched for age and BMI; non-obese (2 females/ 2 males, age: 45.75±13.27, BMI: 23.27±0.72), obese (3 females/ 4 males, age: 38±5.78, BMI: 53.22±8.94), and obese diabetic (2 females/ 1 male, age: 42±6.16, BMI: 51.3±6.16). All human samples used in the present study were obtained from participants who provided written informed consent according to the University Medical Center Göttingen institutional review board ethical approval (38/4/21).

### Animals

All animal procedures were performed in accordance with institutional and national regulations and were approved by the Animal Experiments Inspectorate, Denmark (2022-15-0201-01284). Animals were housed under pathogen-free barrier conditions at the Department of Biomedicine, Aarhus University, and were fed standard mouse chow *ad libitum*. Animals were housed at 21±2°C with 55±10 relative humidity, *ad libitum* access to water, and a 12-h light/12-h dark rhythm. Wild type C57BL/6N were acquired from Taconic Biosciences. In order to lineage trace endothelial cells, Salsa^iEC^ mice were generated by crossing inducible Cdh5(PAC)-CreERT2 mice (151520, Ximbio) with *Gt(ROSA)26Sor^tm1.1(CAG-tdtomato/GCaMP6f)Mdcah^*/J) (kindly provided by Felicity M. Davis). Mice were weaned at 6 weeks of age and received intraperitoneal tamoxifen (T5648, Sigma-Aldrich) in corn oil (C8267, Sigma-Aldrich) injections (10mg/mL) for five consecutive days followed by a 2-day washout period and one additional injection of tamoxifen. Following successful recombination, endothelial cells in Salsa^iEC^ mice constitutively expressed tdTomato allowing for their tracing following long-term intervention. Salsa^iEC^ mice were maintained on a control diet (D12450H) for 12 weeks. At sacrifice, mice were anesthetized through an intraperitoneal injection of ketamine (0.1%) (100 mg/kg) (511485, MSD Animal Health) and xylazine (0.05%) (10mg/kg) (148999, Bayer) in 0.9% sodium chloride solution. Subsequently, mice were perfused with 5 mL PBS, following which, adipose tissue was dissected, weighed, and further processed for digestion as described below. Mice were then perfused with 5mL of 1% PFA (AMPQ40506.5000, Ampliqon) in PBS (SH30256.01, Cytiva), and organs were dissected, weighed, and further processed. *Zeb2*^ΔEC^ mice were generated by crossing *Cdh5*-CreERT2 mice with *Zeb2*^flox/flox^ mice (on a 129 Sv/CD1 background) carrying a *Zeb2* exon 7 flanked by *LoxP* sites and were kindly provided by Aernout Luttun^60^. At 5 weeks of age, Cre-mediated recombination was induced by tamoxifen (intraperitoneal injections of 1mg tamoxifen in 100μL sunflower oil for 5 consecutive days). At eight weeks of age, *Zeb2*^ΔEC^ and *Zeb2*^flox/flox^ mice were maintained on either a western diet (high-fat/high sucrose/high cholesterol; TD.88137, Ssniff) or a control diet (Rat/Mouse – Maintenance, Ssniff) for a duration of eight weeks. *Twist1*^fl/fl^ *Cdh5*^CreERT2/+^ *ApoE*^-/-^ mice (*Twist1*^ΔEC^ mice) were generated by crossing *Twist1*^fl/fl^ *Cdh5*^CreERT2/+^ with *ApoE*^-/-^ mice (all mice were on a C57BL/6J background)^61^. *Twist1*^fl/fl^ *Cdh5*^CreERT2/+^ *ApoE*^-/-^ and *Twist1*^+/+^ *Cdh5*^CreERT2/+^ *ApoE*^-/-^ aged 8 weeks were given a western diet (TD.88137, Envigo) for 8 weeks, followed by intraperitoneal tamoxifen injections (2mg per day for 5 consecutive days) and a high fat diet for 6 weeks.

### Genotyping

Mouse tails were incubated in 25 mM NaOH/0.2 mM EDTA, pH 12 for 30 minutes at 95°C, followed by the addition of 40 mM Tris-HCl, pH 4.7 to extract DNA. Lysed tail DNA was then mixed with HOT FIREPol Blend Master mix (04-25-001125, Solis BioDyne) and the appropriate primers to a final concentration of 0.3 μM, followed by thermocycling with either of the following conditions: 95°C for 15 minutes, 30 cycles of 95°C, 15s; 60°C, 30s; 72°C, 30s; followed by 72°C for 5 minutes (for Cdh5-Cre) and 95°C for 15 minutes, 35 cycles of 95°C, 15s; 57°C, 30s; 72°C, 30s; followed by 72°C for 5 minutes (for Salsa). The products were then analysed by gel electrophoresis. Genotyping primers used for Cdh5-Cre were as follows: mVE-cadherin forward: 5’- ACACCTGCTACCATATCATCCTAC-3’, mCre 73 reverse: 5’-CATCGACCGGTAATGCAG-3’, mGAPDH forward: 5’- CCACTCACGGCAAATTCAACGGCA-3’, mGAPDH reverse: 5’-TCCAGGCGGCACGTCAGATCCACG-3’. Genotyping primers used for Salsa were as follows: Salsa WT forward (IMR 7318): 5’-CTCTGCTGCCTCCTGGCTTCT-3’, Salsa WT reverse (IMR 7319): 5’-CGAGGCGGATCACAAGCAATA-3’, Salsa mutant reverse (IMR 7320): 5’- TCAATGGGCGGGGGTCGTT-3’. Genotyping primers used to genotype *Zeb2*^ΔEC^ and *Twist1*^ΔEC^ mice were previously described^60,61^.

### Isolation of human adipose tissue nuclei

Nuclei were isolated from human subcutaneous adipose tissue samples according to a previously published protocol with slight modification^93^. Approximately 1g of frozen human subcutaneous adipose tissue was finely minced in a petri dish on dry ice and transferred to a 15mL douncer containing 6.5mL Tween salt-Tris (TST) buffer [ST buffer (146mM NaCl (AM9759, Invitrogen), 10mM Tris-HCl (PH 7.5) (15567027, Invitrogen), 1mM CaCl_2_ (97062-820, VWR International Ltd), 21mM MgCl_2_ (M1028, Sigma-Aldrich), 1mM DTT (R0861, Thermo Scientific), and 0.2U/μL RNAse inhibitor (10777019, Invitrogen) containing 0.01% BSA (A7906, Sigma-Aldrich) and 0.03% Tween 20 (P1379, Sigma-Aldrich) in nuclease-free water (AM9930, Invitrogen)] on ice. Samples were homogenized using the douncer, applying 7 strokes with the loose pestle and 5 strokes with the tight pestle. Another 3mL of TST buffer was added and the homogenate was left to rest on ice for 10 minutes, during which occasional strokes were applied to ensure complete homogenization. The homogenate was then filtered through a 40μm cell strainer (732-2757, VWR Collection) and thoroughly washed with 9mL of ST buffer, then centrifuged at 500 x g for 10 minutes at 4 °C. The nuclear pellet was washed once with 3mL ST buffer and then twice with 3mL FANS buffer (1% BSA, 1mM DTT, and 0.2U/μL RNAse inhibitor (03 335 402 001, Protector RNAse inhibitor, Roche) in RNAse free DPBS (AM9624, Invitrogen)), and was centrifuged at 500g for 10 minutes at 4 °C after each wash. Finally, the nuclear pellet was resuspended in 750μL FANS buffer and filtered through a 40μm cell strainer prior to nuclear sorting. All solutions were sterile filtered prior to use.

### Fluorescence-activated nuclear sorting

Isolated nuclei were stained with Hoechst (62249, Thermo Fisher Scientific) and subjected to fluorescence-activated nuclear sorting (FANS) on a 4 laser FACSAria III high speed cell sorter into FANS buffer containing 2% BSA. A detailed gating strategy of nuclei is presented in Supplementary Fig. 1a. A small proportion of the sorted Hoechst-positive events were stained with propidium iodide (PI) and counted on a Luna-FL automated fluorescence cell counter (Logos Biosystems, South Korea) and nuclei were submitted for snRNA-seq using the 10x Genomics system.

### Single-nucleus RNA-sequencing

Immediately following FANS, nuclei were loaded onto the 10x Genomics Chromium controller. Libraries were prepared according to the manufacturer’s instructions using the 10x Genomics Single-Cell 3’ v3.1 (Chromium Next GEM Single Cell 3’ Kit v3.1, 16 rxns (PN-1000268, 10x Genomics), Chromium Next GEM Chip G Single Cell Kit, 48 rxns (PN-1000120, 10x Genomics), and Dual Index Kit TT Set A, 96 rxns (PN-1000215, 10x Genomics). Libraries were sequenced on a DNBSEQ-G400 platform (MGI Tech).

### Human SAT publicly available and in-house datasets processing, integration benchmarking, graph embedding, and visualization

An overview of all datasets included in the SAT atlas is available in (Supplementary table 1 and 2). The dataset repositories GEO, dbGaP, and EMBL-EBI were searched for scRNA-seq and snRNA-seq datasets of human SAT generated using the 10x Single Cell RNAseq and the 10x Single Nucleus RNAseq technologies. Donors were categorized into donors with or without obesity and type two diabetes as described above. FASTQ files containing raw sequencing reads or pre-processed Seurat (RDS) files were either downloaded from the appropriate repositories^2–7^ or were provided by the authors^8^. Raw fastq files were processed to produce count matrices using cellranger v6.1.2 (10x Genomics) by mapping to the human GRCh38 reference genome^94^; *CellBender* (v0.2.2)^95^ was used to remove empty droplets and ambient RNA. For each individual sample, the output from CellBender was read in R using the function Read10x_h5 provided by Seurat. Then a Seurat object was created with *min.cells* of 3 and *min.feautures* of 200. Ribosomal, mitochondrial, and hemoglobin genes, as well as other confounding transcripts including *MALAT1* and *NEAT1* were removed prior to downstream data processing. Outliers were removed based on the number of Unique molecular identifiers (UMIs) and genes detected per cell. A linear model was constructed based on the log of UMIs against the log of genes and outliers were removed. *DoubletFinder* (v2.0.3)^96^ was applied to exclude potential doublets. All samples from all studies were merged and outliers of UMIs and genes were removed. Data was log normalized using *NormalizeData* function then multiplied by a scale factor of 10000. Top 2000 highly variable genes were selected using vst method and scaled regressing out the number of UMIs. Dimensionality reduction was performed using principal component analysis (PCA) and projected using Uniform Manifold Approximation and Projection (UMAP).

### Data integration and benchmarking

Different integration methods, namely *Harmony* (v0.1.1)^97^, single-cell variational inference (*scVI*) (v1.0.4)^98^, and batch balanced K-nearest neighbor (*BBKNN)* (v1.6.0)^99^ have been evaluated for batch effect correction.

*Integration with Harmony*. We used *RunHarmony* to integrate the data, with samples, cohort, and chemistry (Drop-seq and Chromium versions 2, 3, and 3.1) as covariates. The first 43 harmony components were selected for generating the UMAP and clustering was performed at different resolutions assessed by the clustree package (v0.5.0)^100^.

*Integration with scVI.* Data was processed as previously described, then the top 2000 highly variables genes were used for integration. The Seurat object was converted into AnnData file format and processed with reticulate package (v1.34). Function setup_anndata was applied specifying the samples, cohort, and chemistry as a batch key argument for batch correction. Then scVI model was created, trained and the latent representation was obtained. The resulted matrix was used as input to CreateDimReducObject and add the embeddings were added to the main object. The first 10 PCs were selected to construct the UMAP.

*Integration with BBKNN.* We used the function bbknn, with samples, cohort, and chemistry as a batch key argument. Clusters were identified using Leiden algorithm at resolution 0.1. Then ridge regression was performed over the integrated data. Then bbknn integration was repeated using leiden clusters as a confounder key, followed by PCA analysis and UMAP construction.

The assessment of the three integration approaches was carried out based on the calculation of LISI scores, kBET acceptance rates (1- rejection rate), and ARI coefficients for integration across technologies, cohorts, and samples. We estimated average LISI scores using *compute*_*lisi* command (v1.0)^97^ by adding embeddings of the UMAPs along the metadata. kBET scores were calculated based on Pearson’s chi-squared test using kBET package (v0.99.6)^101^ and the different integrated reductions in the Seurat object. ARI was calculated using the *adj.rand.index* function in *pdfCluster* (v1.0.4)^102^ by supplying factors of cohort, technology chemistry(Drop-seq and Chromium versions 2, 3, and 3.1 – single cell or single nucleus), or sample and the identified clusters. Additionally, we also benchmarked the different integration strategies using scIB-metrics^103^, which provides a consolidated evaluation of integration taking into account metrics of batch effect removal (kBET, k-nearest-neighbor (kNN) graph connectivity, the average silhouette width (ASW) across batches, iLISI, and PCA regression) and biological variability conservation (cell identity labels (label conservation), label-free conservation, graph cLISI, ARI, normalized mutual information (NMI), relative distances (cell-type ASW), isolated label scores and label-free conservation). To this end, unintegrated, *Harmony*, *scVI*, and *BBKNN* objects were merged into one Seurat object using *CreateDimReducObject* then converted to an *AnnData* object to be used for benchmarking. Benchmarking was then carried out using scIB Benchmarker library with the batch key set to the samples, study, and chemistry, while the label key was set as the cell annotation. Results were visualized using the *plot_results_table* method. Guided by our benchmarking, *Harmony* integration collectively ranked the highest, in terms of both, batch effect correction and the conservation of biological variation, and therefore it was applied for the downstream analyses.

### Clustering, cellular population annotation, and subpopulation identification

Clusters were identified after running *FindNeighbors* with the first 43 harmony corrected PCs and *FindClusters* in Seurat at different resolutions ranging from 0.1 to 3. To assign clusters to different cell types, *FindAllMarkers* was used with *min.pct* 0.15 *logFC threshold* 0.3 using Wilcoxon rank sum test. At resolution 0.1, *FindSubcluster* was utilized to subcluster a widely dispersed cellular population where subclusters without noteworthy gene expression differences were collapsed and only high-quality cells were retained for downstream analysis. Clusters were manually assigned to discrete cell types based on the top marker genes of each cluster and 9 distinct cellular populations were identified. Cellular populations were annotated as; adipocytes and preadipocytes (*ADIPOQ*, *AQP7*), ASPCs (*APOD*, *CXCL14*), adipose endothelial cells (AdECs) (*PECAM1*, *CDH5*), adipose lymphatic endothelial cells (ALECs) (*MMRN1*, *PROX1*), mural cells (*RGS6*, *ACTA2*), macrophages and monocytes (*MRC1*, *FMN1*), mast cells (*CPA3*, *HPGD*), B cells (*IGKC*, *BANK1*), and T and NK cells (*CXCR4*, *CD69*). Each major cell population was then sub-clustered to a comparable granularity as described below. Marker gene lists of all the subpopulations are available in Supplementary Table 30.

*ASPCs*. ASPCs were subset and dimensionality reduction was performed following data normalization and scaling, regressing out the number of UMIs per cell. Batch effect was corrected with *Harmony* using 30 harmony components. Cells were then clustered at a resolution of 0.2, yielding six clusters. Marker genes for each cluster were then computed using *FindAllMarkers* function from Seurat with *min.pct*=0.15 and *logfc.threshold*=0.3. The six ASPC populations exhibited divergent transcriptomic signatures and based on their expression of canonical marker genes^7,8^, were annotated as ASPCs 1 to ASPCs 6.

*Adipocytes*. Adipocytes and pre-adipocytes (which clustered together) were subset. The annotation of single cell-derived pre-adipocytes was retained and the single nucleus-derived adipocytes were further sub-clustered. Data was normalized and scaled regressing out the number of UMIs per cell before dimensionality reduction. Following batch effect correction with harmony (using 30 harmony components), clusters not presenting bona fide adipocyte populations and that resembled other cell types were removed. Moreover, clusters comprising cells from only one study or clusters with low UMI counts were also removed, following which, data was reprocessed, reintegrated (using 30 harmony components) and re-clustered at a resolution of 0.1, yielding six clusters. Clusters without noteworthy gene expression differences were collapsed to yield coherent cell populations. Adipocyte populations were annotated based on their expression of previously published adipocyte marker genes^7,8^ and were annotated as Adipocyte 1 to Adipocyte 6.

*Vascular cells*. High-quality vascular cells were selected and subjected to another iteration of highly variable gene selection, dimensionality reduction, data integration, and clustering analysis using 30 harmony components at resolution 0.7. The heterogenous population of sub-endothelial cells were then subset for further analysis applying the same pipeline as performed over the vascular cells, selecting the first 15 harmony components and clustering at resolution 0.2.

*Macrophages and monocytes*. Macrophages and monocytes were similarly subset and the data was normalized and scaled (regressing out number of UMIs per cell) before dimensionality reduction was performed. Batch effect correction was done using *Harmony* (30 harmony components) and clusters not presenting bona fide macrophage and monocyte populations and that resembled other cell types were removed. Moreover, clusters comprising cells from only one study were removed, following which data were reprocessed, reintegrated (30 harmony components), and re-clustered at a resolution of 0.3 yielding ten clusters, of which one was subclustered to yield coherent populations. Monocyte, dendritic cell, and macrophage subpopulations were annotated based on their expression of canonical marker genes^4,7,8^ and were annotated as Macro/Mono 1 to Macro/Mono 10.

*T and NK cells*. T and NK cells were subset and dimensionality reduction was performed following data normalization and scaling, regressing out the number of UMIs per cell. Batch effect correction was done using *Harmony* (30 harmony components) and cells were clustered at resolution 0.2. Cell clusters not representing *bona fide* T and NK cell populations and that resembled other cell types were removed and the data was re- processed. Batch effect was then corrected using 30 harmony components and cells were clustered at a resolution of 0.5, yielding cell clusters derived from all the included datasets. T and NK cell populations were annotated based on their expression of canonical marker genes^4,7,8^ and were annotated as T/NK 1 to T/NK 9.

### Compositional analysis

To assess changes in cell composition we used *scCODA*^104^, a Bayesian model for compositional single cell data analysis. Only single nucleus RNA sequencing data was used as it comprised all the major cell populations. Non-obese samples were set as the reference condition and arterial endothelial cells, being the cluster with the lowest variability across samples and with a low amount of dispersion, were selected as reference. *scCODA* was ran ten times using the Hamiltonian Monte Carlo sampling method with default parameters. Results were averaged per cell type and average scores were used as a measure of cell type abundance.

### Module score calculation, GSVA, and PROGENy

Vascular cells were scored for the expression of widely characterized vascular bed-specific and mural cell-specific gene signatures using the *AddModuleScore* function from *Seurat* with default parameters (Supplementary Table 8). ECM components and regulator scores in vascular cells were computed based on gene lists from the matrisome database^18^. The single cell implementation of gene set variation analysis (GSVA)^105^, scGSVA (https://github.com/guokai8/scGSVA) was also used to compare scores of gene set signatures among vascular cell subpopulations. scGSVA was implemented with default parameters using the *UCell* package^106^. Pathway activity scores were also calculated using PROGENy^107^, which leverages publicly available perturbation experiments to yield a common core of pathway responsive genes.

### Differential Expression and Gene Ontology (GO) Analysis across metabolic states

*FindMarkers* function was used to compare every cellular population between each two metabolic states at a time employing a logistic regression test with sex, cohort, and chemistry as covariates. The upregulated and downregulated genes (average log2 fold change above and below 0.5) were then used as input for the *enrichGO* function from clusterProfiler package (v4.8.3)^108^. All identified genes were used as background for GO enrichment. GO terms of deregulated genes with p adjusted values following Bonferroni correction inferior to 0.05 were considered significant. GO terms were arranged by q value and top GO terms were visualized. Top 5 GO terms unique to a specific comparison were visualized by bar plots. GO terms shared between both pairwise comparisons were represented as radar charts by calculating the average area under the curve (AUC) of each gene list per metabolic state using *AUCell_run* from AUCell package (v1.22.0)^109^. GO terms in bold have been highlighted in the text.

### Transcription factor activity and gene regulatory network inference

To infer transcription factor activities, two alternative approaches were performed. First, we estimated relative transcription factors activities from the collected datasets, we used the top 3 confidence levels of the DoRothEA database (v1.12.0). We used the function run_mlm from decoupleR (v2.6.0)^110^ to fit a multivariate linear model for each sample with setting the *min.size* to 5 genes per transcription factor. Transcription factor classes were adopted from *TFClass*^111^. Transcription factors in bolded red have been highlighted in the text. Second, gene regulatory networks were inferred using pySCENIC (v0.12.1)^112^. pySCENIC infers regulons (co-expression modules) composed of a given transcription factor and its putative targets, measuring their activity in individual cells. Co-expressed modules were inferred from a predefined curated list of 1,892 human transcription factors provided by the pySCENIC repository using the *GRNBoost2* algorithm with the transposed count matrix as input^113^. The output expression matrix adjacencies file was used as input in addition to cistarget databases to generate regulons. The aucell command was then ran with the regulons along with a loom file created from the anndata file as input and the generated area under the curve matrix was summarized by the mean of each cell type.

### Murine and porcine SAT publicly available data processing, reference mapping, and integration across species

Murine snRNA-seq data generated by Emont M. et al. was downloaded from the online repository^7^. Only inguinal samples were selected for further analysis. Murine data was processed as previously described. Briefly, ribosomal and mitochondrial genes, *Malat1* and *Neat1*, in addition to hemoglobin genes were removed. Datasets were filtered out for outliers based on the number of UMIs and genes for each cell. Linear model was constructed and only residuals of more than -0.5 were kept. Then data integration with harmony was performed over the samples, choosing 30 harmony components, and clustered using the Louvain algorithm at resolution 0.5. High-quality vascular cells were selected and processed as mentioned before for the human data with the top 30 PCs at resolution 0.5. Murine scRNA-seq data generated using sorted eYFP^+^ cells from Cdh5-CreER^T2+^ ROSA26-eYFP mice by Dunaway L. S. was downloaded from the online repository^59^. This dataset included cells isolated from mesenteric and epidydimal fat depots and was processed as described above. Data integration was similarly done using harmony, choosing 30 harmony components, and clustering was performed using the Louvain algorithm at resolution 0.9. Porcine scRNA-seq data generated by Wang F. et al. was downloaded from the online repository^30^. Subcutaneous adipose tissue cells were retained followed by highly variable gene selection and dimensionality reduction. The first 25 PCs were selected to construct the UMAP, and clustering was performed at different resolutions, later evaluated with clustree and resolution 0.3 was chosen. Vascular cells were selected, highly variable genes recalculated and the first 19 PCs were used for UMAP and clustering at resolution 0.3. Reference mapping was performed using scmap (v1.22.3)^114^ package setting *nFeatures* to 5000 genes. We used the *scmapCluster* algorithm setting the human dataset as query, and either the murine or the porcine datasets as reference with threshold 0 opting to minimize unassigned cells. Murine orthologous genes were mapped using the *covert_mouse_to_human_symbols* from the *NicheNet* package. Porcine genes were downloaded from the Pig RNA Atlas and EnsemblID were converted into gene symbols using *biomart* (v.2.60.1)^115^. Data were visualized using the *getSankey* function and the resulting table was used to evaluate the *scmap* similarity index. Alternatively, murine and porcine vascular cells were integrated with their human counterparts using a conditional variational autoencoders (cVAE)-based model, *sysVI*^116^, which combines variational mixture of posteriors prior (VampPrior) and latent cycle-consistency loss to integrate datasets with substantial batch effects. Briefly, the human and murine/porcine data were merged into one anndata file. Then data were processed with scanpy using *pp.normalize_total*, *pp.log1p*, and *pp.highly_variable_genes* with setting *batch_key*=’system’ in the latter function. Approximately 2000 shared highly variable genes were selected and subset to train the model with *batch_key*=’system’ and samples, cohort, and chemistry used set as categorical covariate keys for 200 epochs and default parameters before using *get_latent_representation* function. We then calculated *pp.neighbors* and *tl.umap* and visualized the embedding for both species using *pl.embedding*.

### Jaccard Similarity Analysis

Clusters of endothelial cells belonging to a specific vascular bed were combined. Marker genes were calculated for each cell type per metabolic state using the Seurat pipeline. Then the top 100 markers for each cell population were selected for pairwise Jaccard similarity coefficients calculation. The Jaccard coefficient is the size of intersection divided by the union sets size.

### Computation of Endothelial, Mesenchymal, and EndMT scores

Endothelial and Mesenchymal scores were calculated as previously described for epithelial-to-mesenchymal transition (EMT)^117^. Briefly, a list of endothelial and mesenchymal markers was compiled through manual curation of the literature, as follows:

- Endothelial genes: *PECAM1*, *CDH5*, and *VWF*.
- Mesenchymal genes: *ACTA2*, *FAP*, *S100A4*, *AIFM2*, *COL1A1*, *COL3A1*, *TWIST1*, *ZEB2*, *SNAI1*, *SNAI2*, *SMAD2*, *SMAD3*, *SMAD4*, *YAP1*, *KDM4B*, *MAPK8*, *MAPK14*, *TGFB1*, *TGFB2*, *TGFB3*, and *TGFBR1*.

EndMT scores for each endothelial cell was computed by subtracting the average RNA-seq z-scores of the endothelial marker genes from the average RNA-seq z-scores of the mesenchymal marker genes. Endothelial and mesenchymal scores were visualized using scatter plots and EndMT scores were scaled and visualized using boxplots.

### Computational trajectory inference

Trajectory inference was performed using the *Monocle3* package (v1.3.1)^118^. Certain populations of ECs (Capillary endothelial cells 1, Capillary endothelial cells 2, Venous endothelial cells 3, Activated vascular endothelial cells, FAP-like endothelial cells, and adipocyte-like endothelial cells) were extracted, data was normalized, top variable genes were selected, scaled and dimensions were reduced by PCA using default parameters. Batch effect was removed using harmony and plotted in 2- dimensional space using UMAP with the significant corrected harmony components. The Seurat object was transformed into a cell data set object and the rest of monocle3 pipeline was performed including *cluster_cells*, *learn_graph*, *order_cells* and *graph_test* for differential gene expression across pseudotime using morans test statistic. Differentially expressed genes changing across pseudotime with q-value exceeding 0.05 were filtered out. Significant differentially expressed genes were aggregated for each cell type using *aggregate_gene_expression* and the top 50 genes per cluster were visualized using the package pheatmap v1.0.12. Selected transcription factors, which expression changed across pseudotime were visualized using ggplot2 (v3.4.4) and their expression was smoothed using loess regression. To complement the trajectory analysis, *CytoTRACE2*^119^ was implemented with default parameters using the raw counts without normalization as input to infer cellular differentiation states. Cell state entropy, as a measure of cell differentiation states, was estimated using single cell lineage inference using cell expression similarity and entropy (*SLICE*)^120^. We performed deterministic calculation of entropy scores of single cells with default parameters. To visualize and characterize putative transitions between cellular states, partition-based graph abstraction (*PAGA*)^121^ was used. *Scanpy*^122^ 1.10.2 package was used to run *scanpy.pp.neighbors* with 30 harmony components. The PAGA graph was constructed using *draw_graph* and *tl.paga* methods. Data was then visualized in a force-directed graph calculated using *scanpy.pl.paga* on the pre-computed coordinates from *PAGA*. The robustness of the Monocle3- inferred trajectory was validated by independent pseudotime prediction tools including *Palantir*^123^, *Slingshot*^124^*, and CellRank*^125^. Slingshot (v.2.7.0) pseudotime inference was performed using *getLineage* and slingshot functions setting the starting and the ending clusters to C1 and Adipocyte-like Endothelial cells, respectively. *Palantir* trajectory prediction was performed using the Palantir python package (v.1.3.6). A diffusion map was generated using the *run_diffusion_maps* function. MAGIC imputation was implemented (*run_magic_imputation*) and *Palantir* trajectory prediction was generated using the *run_palantir* function with *num_waypounts* = 500 and the early cell set as predicted by Palantir (*find_terminal_states*). CellRank was used to predict the initial and terminal cell states in an un-biased manner. To run *CellRank*, *velocyto*^126^ was ran over all in-house generated samples and randomly downsampled 15,000 cells were used for downstream analyses. *scVelo*^127^ (v.0.3.2) was used for RNA velocity analysis. Briefly, data was filtered (*min_shared_counts* = 20, *n_top_genes* = 2000, *subset_highly_variable* = True) and normalized before moments were calculated. The functions *recover_dynamics*, *tl.velocity* (mode = dynamical), and *velocity_graph* were used. *CellRank* was performed using the *CytoTRACEKernel* function with default parameters from the *CellRank* python package (v.2.0.5) and the functions *compute_cytotrace* and *compute_transition_matrix* were used before initial and terminal states prediction were ran with 12 states. Data were visualized using the *CellRank* plotting functions provided on the package Github.

### In silico gene perturbation

*CellOracle* (v.0.20.0) ^128^ was used to simulate transcription factor knockout *in silico*. The *in-house* dataset used to construct the pseudotime trajectory was used as input and the standard *CellOracle* workflow was followed. We used *CellOracle*’s built-in human promoter base gene regulatory network (GRN) as scATAC-seq data of human adipose tissue is not available. PCA was performed before imputing KNN with default parameters. The *get_links* function was used to calculate GRNs for each cluster, then only significant links (p-value < 0.05, threshold_number= 2000) were retained. Network scores were calculated using *get_network_score*. Predictive models were built for simulation using *fit_GRN_for_simulation* function. We focused on a known endothelial-to-mesenchymal transcription factor, *TWIST1*, which was knocked out *in silico* using the *simulate_shift* function. Transition probability between cells was then estimated and the embedding shift was calculated with default parameters. Quiver plots, vector fields, and inner product between vectors were used to visualize the simulated effect of knocking out *TWIST1 in silico*.

### Single-cell Disease Relevance Score (scDRS)

scDRS was used to evaluate polygenic disease enrichment of human SAT vascular cell subpopulations as previously described^86^. Briefly, the publicly available Genome-wide association studies (GWAS) summary statistics of 74 diseases and traits (average N=346.000) were downloaded, and relevant traits were selected for downstream analysis. Statistical significance was determined by generating thousand sets of cell-specific raw control scores of Monte Carlo samples of matched control gene sets where the gene set size, mean expression, and expression variance of disease-relevant genes were matched. Raw disease and control scores of each cell were then normalized and cell-level p-values were computed based on the empirical distribution of the pooled normalized scores.

### Identification of druggable targets in vascular cells

To identify potential druggable targets in the vascular cells in obesity and T2D, we performed three complementary target prioritization methods. First, we performed a series of filtering steps on genes tested for differential expression in the Obese vs. Non-obese and Obese diabetic vs. Non-obese comparisons. *FindMarkers* function employing a logistic regression test with sex, cohort, and chemistry as covariates was used for pairwise comparisons and upregulated and downregulated genes (average log2 fold change above and below 0.5) were then used as input. Only differentially expressed protein-coding genes that exhibited a strong effect (absolute value of fold change > 1) in either comparison with expression detected in at least 25% of cells comprising each population were retained. We then annotated these genes by whether they were found in the druggable genome database based on Finan et al.^129^ (which incorporated 4,479 genes) and only retained genes that exhibited cell-type specificity (based on the calculation of AUC) using FindAllMarkers (test.use= "roc") with AUC > 0.7. We included druggable targets of all tiers where Tier 1 genes (1,427 genes) represent efficacy targets of approved small molecules and biotherapeutics as well as clinical-phase drug candidates. Tier 2 (682 genes) included genes encoding targets with known bioactive drug-like small-molecule binding partners as well as genes encoding targets with ≥250% identity with approved drug targets. Tier 3 genes (2,370 genes) included those encoding for secreted or extracellular targets with more distant similarity to approved drug targets as well as members of key druggable gene families not included in Tiers 1 and 2. Genes belonging to Tier 3 were stratified into Tier 3A (genes proximal (±50 kbp) to a GWAS SNP and had an extracellular location) or Tier 3B. Druggable targets in each vascular population were then visualized using ridge plots. Full list of druggable targets with their tiers, including targets that did not meet the predetermined AUC threshold are depicted in Supplementary table 22. Second, we used *Drug2cell* (v.0.1.1)^79^, which employs pairs of drugs and their targets from the ChEMBL database, to infer drug targets in vascular cell subpopulations as clusters of interest. To that end, the vascular *Seurat* object was converted into an *AnnData* file and pairwise scoring of each cell population was achieved by running the *score* function using the default parameters. Differential expression was performed using *rank_genes_groups* on the calculated scores comparing obese or obese diabetic states to the non-obese one using Wilcoxon rank sum test. Third, we utilized the GREP (Genome for REPositioning drugs) software^80^ and performed an enrichment analysis of metabolic state-based differentially expressed genes in the different vascular cell subpopulations in the drug targets of clinical indications categorized by disease chapter and captured potentially repositionable drugs. Two publicly accessible databases were used for information on approved drug targets or clinical trial targets; Drug bank: https://www.drugbank.ca/ and Therapeutic Target Database: https://db.idrblab.net/ttd/. The number of validated, repurposing, and unestablished gene-disease pairs are presented in Supplementary Table 35.

### Connectome analysis

Differential ligand-receptor analysis was performed using the packages LIANA^+^ (v0.1.9)^68^ and *CellChat* (v1.6.1)^67^ using the annotated data with high granularity. Briefly, using *CellChat*, cell annotation labels were transferred back into a Seurat object comprising all cells. Low quality cells identified within each cell type Seurat object were excluded from the analysis. Single dataset connectome analysis was performed individually on non-obese, obese, and obese diabetic cells using “Secreted signaling”, “ECM-Receptor”, “Cell-Cell contact”, and “Non-protein Signaling” data from the CellChatDB database and the functions *identifyOverExpressedGenes*, *identifyOverExpressedInteractions* and *computeCommunProb*. Communications in less than 10 cells were filtered out. Next, *computeCommunProPathway* and *aggregateNet* were used and connectome analyses comparing different datasets (non-obese, obese, and obese diabetic) were carried out. For each pairwise comparison, individual CellChat objects were merged using *mergeCellChat* to calculate differential interactions through *compareInteractions* and *netVisual_diffInteraction* functions. To visualize the incoming and outgoing cellular interactions in 2D space, *netAnalysis_signalingRole_scatter* and *netAnalysis_signalingChnages_scatter* functions were used. The results were visualized using heatmaps, circular diagrams and bar plots. All functions were used with default parameters unless otherwise specified. For the analysis using *LIANA^+^* (v.0.1.9), an anndata file with highly granular cell type annotation was converted from Seurat and low-quality cells were removed as described above. For each metabolic state subset, data was normalized and log-transformed. The method *rank_aggregate* was used with consensus set as *resource_name* which runs *CellPhoneDB*, *Connectome*, *log2FC*, *NATMI*, *SingleCellSignalR*, and *CellChat*. Number of interactions was aggregated per interacting cell pair. Focusing on specific interacting cell pairs, we used *NicheNet (v.2.2.0)*^71^ to predict the influence of ligands from sending cells on gene expression of receiving cells. Briefly, g*et_expressed_genes* function (pct= 0.05) was used for sending and receiving cell clusters separately before running Seurat’s FindMarkers between metabolic states of interest using logistic regression test to account for sex, study, and chemistry covariates with the non-obese state set as reference. The list of differentially expressed genes were then filtered with p-adjusted value below 0.05 and average logFC more than 25%. We then used *predict_ligand_activities* and the results were ranked by *aupr* values where the top 30 ranked ligands were visualized. To infer the target gene and receptor of top-ranked ligands, *get_weighted_ligand_receptor_links* function was used with default parameters. All plots were generated using the *ggplot_R* package as described in the *NicheNet* Github.

### Spatial deconvolution

*Cell2location*^130^ (v.0.1.4) was used for deconvolution of publicly available human SAT spatial transcriptomics data (n=10)^17^ using our fine-grained annotated SAT atlas data as reference. The analysis was conducted with default parameter settings. To assess cell type co-localization, spot-wise Pearson correlations were computed for each subject across all pairs of cell types. The correlation values were then averaged across subjects to obtain representative results. A low pearson correlation coefficient suggested distinct spatial distribution between two given cell types, while a high correlation indicated that two cell types exhibited similar spatial distributions. Spots were also scored based on a curated list of core endothelial cell marker genes to mark the spatial distribution of endothelial cells (Supplementary Table 23).

### Isolation of human and murine adipose tissue stromovascular fraction and endothelial cells

Human adipose tissue endothelial cells were isolated from fresh human adipose tissue according to a previously published protocol with slight modifications^93^. Human deep subcutaneous adipose tissue samples were obtained through the same abdominal surgeries described earlier at the Department of Internal Medicine, Viborg Regional Hospital, Viborg, Denmark according to the approval of the Scientific Ethics Committee for the Central Denmark Region (record number 1-10-72-312-20) and the presence of informed written consent from all participants. Adipose tissue was cut into small pieces and digested for 30 minutes at 37 °C in supplemented KnockOut^TM^ DMEM-based digestion buffer (10829018, Gibco) containing 1x Penicillin/Streptomycin (15140122, Gibco), 2x Antibiotic-Antimycotic (15240062, Gibco), 1mM Sodium Pyruvate (11360070, Gibco), 1x MEM Non-essential amino acids (11140035, Gibco), 2mM L-glutamine (25030024, Gibco), and 1x Endothelial Cell Growth Factor supplements (ECGS/heparin) (C-30120, PromoCell) in addition to 0.1% Collagenase I (17100017, Thermo Fisher Scientific), 0.25U/mL Dispase (17105041, Gibco), and 7.5mg/mL DNAse I (D4527, Sigma-Aldrich). The solution was then filtered through a 100μm cell strainer and allowed to rest. The floating fat layer was removed, and the reaction was stopped with PBS-based wash buffer containing 0.5% BSA (A7906, Sigma-Aldrich) and 2mM EDTA (E177, VWR Chemicals). The solution was then spun down at 300 x g for 7 minutes and the pellet was either resuspended in wash buffer for downstream endothelial cell isolation or was resuspended in 1ml StemMACS^TM^ Cryo-Brew solution (130-109-558, Miltenyi Biotec) and frozen until later use. For downstream endothelial cell isolation, cells were washed twice with wash buffer and then incubated with CD45 human Microbeads (130- 045-801, Miltenyi Biotec) for 20 minutes. Cells were then washed with wash buffer and passed through an activated MS column (130-042-201, Miltenyi Biotec). The column was washed three times. The eluted solution was centrifuged, and cells were incubated with CD31 human Microbeads and FcR Blocking Reagent (130-091- 935, Miltenyi Biotec) according to the manufacturer’s protocol for 20 minutes. Cells were washed and passed through an activated MS column as previously described. Magnetically labeled cells were immediately flushed out by firmly pushing the plunger in the column. Isolated cells were plated on 0.1% gelatin-pre-coated culture flasks in supplemented Endothelial Cell Growth Medium 2 (ECGM2) (C-22011, PromoCell) containing 2x Antibiotic/Antimycotic at 37 °C, 5% CO2, and were passaged at confluence. Cells were used in single-donor cultures and between passages 1 and 5.

### Isolation of human umbilical vein endothelial cells (HUVECs) and the generation of mCherry-expressing HUVECs

HUVECs were freshly isolated from the umbilical cords of multiple donors and tested for mycoplasma (6DV1-MPCBC, Eurofins). All umbilical cords were donated from the Department of Obstetrics and Gynecology, Aarhus University Hospital, Denmark. Briefly, umbilical cords were collected and rinsed with preheated, sterile PBS + 2X antibiotic-antimycotic (AB/AM) (15240062, Gibco) and sterile gauze. After cutting both ends (approx. 2 cm) with sterile scissors, the vein was rinsed by injecting 20 mL preheated, sterile PBS + 2x AB/AM with a sterile feeding needle (Agnthos, 7902) and the bottom was clamped off with a sterile clip. Next, 10 mL preheated 0.2% collagenase type I in 0.9% NaCl + 2mM CaCl2 + 2x AB/AM was injected with a sterile feeding needle, after which the other side was clamped off and the umbilical cord was incubated for not more than 13 min. at 37°C. The collagenase mix was then collected in a 50 mL Falcon tube, and the vein was rinsed with 20 mL Medium 199 (M199) (22340-020, Gibco) supplemented with 2 mM L-Glutamine, Endothelial Cell Growth Supplement/Heparin (C-30120, PromoCell) and 20% FBS, which was collected in the same tube. The total volume was then filtered through a 40 µm nylon cell strainer and collected in a fresh tube. After centrifugation for 5 minutes at 1000 rpm and aspirating the supernatant, the pellet was resuspended in 10 mL of supplemented M199 and transferred to a T-75 flask precoated with 0.1% gelatin from bovine skin (G9391, Sigma-Aldrich) in PBS, which was removed after 15 minutes of incubation at 37°C. The day after isolation, cells were washed twice with PBS and added fresh supplemented M199 medium with 2X AB/AM added during the first week after isolation but not after. HUVECs were maintained in ECGM-2 media (C-22011, PromoCell) or M199 full media (M199 media (22340020, Gibco), 1x L-Glutamine (2mM) (25030-024, Thermo Scientific), 20% FBS, (F7524, Sigma-Aldrich), ECG/H supplement(C-30120, PromoCell). Cells were used in single-donor cultures and were between passages 1 and 5. mCherry expressing HUVECs were generated by homologous recombination using CRISPR/Cas9 and rAAV6 homologue donor delivery as previously described^131^. Briefly, 150,000 HUVECs were nucleofected in OptiMEM (31985062, Gibco) containing Alt-R® S.p. Cas9 Nuclease (1081059, IDT), AAVS1 gRNA, and AAV6-AAVS1-SFFV-mCherry homology directed repair template^131^ (kindly provided by Rasmus O Bak) using a Lonza electroporation cassette (P3 Primary Cell 4D-Nucleofector X kit) with the primary cell P3 program (pulse code: CM138). Cells were then grown to confluency in ECGM-2 media and once confluent, mCherry positive cells were sorted, grown to confluency in supplemented ECGM-2 media, trypsinized, and frozen until further used.

### Human adipose tissue organoid culture and immunostaining

Primary human SVF cells were used to form human adipose tissue organoids as previously described^53^. Briefly, primary human SVF cells (Lonza Bioscience, Switzerland) combined with mCherry expressing HUVECs (5% of seeded cells) were seeded in a 96-well ultra-low attachment plate with a round bottom (CLS7007, Corning) at 10,000 cells per well in EGM-2 (CC-3162, Lonza) in a 5% CO2 humidified incubator at 37 °C. On day 6 (d6), spheroids were then embedded in growth factor-reduced Matrigel (11553620, Corning) and maintained in EGM-2 until d10. On d10, half of the cell culture media was replaced with 2x concentrated preadipocyte growth medium (PGM-2, PT-8002, Lonza) and media was changed on d15 and d20. Organoids were harvested on d10, d15, and d20, fixed with 4% PFA for 30 minutes, and then washed and maintained in PBS until stained. Organoids were incubated in blocking buffer (3% BSA 1% Tween-20 in PBS) for 2 hours and then incubated with primary antibodies (1:100 mouse anti-Perilipin 1 (690156S, Progen) and 1:100 rabbit anti-RFP (600-401-379-RTU, Rockland)) overnight at 4 °C. Organoids were then washed and incubated with BODIPY 493/503 (D3922, Invitrogen), 1:100 donkey anti-mouse Alexa Fluor 647 (A-31571, Invitrogen), and 1:100 donkey anti-rabbit Alexa Fluor 568 (A10042, Invitrogen) for two hours at room temperature. Organoids were then washed and counterstained with Hoechst (1:500) prior to mounting.

### In vitro induction of EndMT

Primary human AdECs were cultured in ECGM2 supplemented with endothelial cell growth supplement on 0.1% gelatin coated culturing flasks in a 5% CO2 humidified incubator at 37 °C. Media with supplements was replaced three times per week and cells were passaged at confluence by washing once with PBS, trypsinization, and centrifugation at 500g for 5 minutes. For the induction of EndMT, AdECs were seeded at a density of 150,000 cells/well in 12-well plates. The next day, supplemented ECGM2 was replaced with a 1:1 media comprising supplemented ECGM2 and M-199 (22350078, Gibco). Cells were then either treated with a combination of TGF-β2 (100-35B, PeproTech) (10ng/mL), TGF-β1 (100-21, PeproTech) (10ng/mL), and IL-1β (200-01B, PeproTech) (10ng/mL), or their respective vehicles according to the manufacturer’s instructions. Media and treatment were replenished every other day and cells were harvested either at three or seven days following EndMT induction.

### Cell viability assay

Cell viability was assessed using the Cell Proliferation Kit I (MTT) (11465007001, Roche) following the manufacturers’ instructions and absorbance was measured at 570nm using a microplate reader (Biotek).

### Migration assay

AdECs were seeded as described above in 24-well plates coated with 0.1% gelatin and were treated as indicated for 7 days. On day 7, media was discarded and the cells were washed twice with PBS. A single vertical scratch was made using a pipet tip (200μL). Cells then received either the EndMT cocktail or its respective vehicles supplemented with 1.5μg/mL Mitomycin C (M4287, Sigma-Aldrich). Images were taken at zero and eighteen hours using an Olympus light microscope. Images were analyzed using Fiji (v.1.16) and data was expressed as scratch area relative to vehicle-treated controls.

### Monocyte adhesion assay

Primary AdECs seeded on coverslips as described above were treated as indicated for 7 days. THP-1 leukemic monocytes were maintained in RPMI (R8758, Sigma-Aldrich) supplemented with L-glutamine (2mM), 10% fetal bovine serum, and 1% penicillin-streptomycin. One million THP-1 cells were pre- stained with 1μM CellTracker Green CMFDA (C7025, Invitrogen) for 30 minutes according to the manufacturer’s instructions. On day 7, AdECs media was aspirated and cells were washed twice with PBS then cocultured with pre-labeled THP-1 cells in RPMI for one hour in the incubator. Non-adherent THP-1 cells were gently washed away and coverslips were mounted, counterstained with Hoechst (1:500), and immediately imaged using a LSM 900 inverted confocal microscope.

### Immunocytochemistry

Cells were seeded at a density of 75,000 cells/mL onto glass cover slips in 24 well plates and treated as described above for a duration of 7 days. Cells were washed with PBS containing MgCl_2_ and CaCl_2_ (SH30264.01, Cytiva) before fixation with 4% PFA for 15 minutes at room temperature. Cells were then permeabilized in 0.5% Triton X-100 (0694, VWR Chemicals) in PBS for 30 minutes at room temperature followed by blocking with 0.5% Triton X-100, 2% BSA in PBS (blocking buffer) for one hour at room temperature. Cells were then incubated with the following primary antibodies overnight at 4 degrees Celcius in blocking buffer; 1:150 goat anti-VE-Cadherin (AF1002, R&D Systems) and 1:500 rabbit anti-ZEB2 (14026-1-AP, Proteintech). Cells were then washed four time (five minutes each) with PBS and incubated with secondary antibodies (1:500 donkey anti-goat Alexa Fluor 647 (A21447, Invitrogen) and 1:500 donkey anti-rabbit Alexa Fluor 568 (A10042, Invitrogen) as well as 1:100 phalloidin 488 (A12379, Invitrogen) and 1:500 Hoechst for 2 hours at room temperature. Coverslips were then washed with PBS and mounted using Fluoromount-G (00-4958-02, Invitrogen). Images were acquired using a LSM 900 inverted confocal microscope.

### RNA isolation, cDNA synthesis, and Quantitative RT-PCR

Total RNA was extracted using TRI Reagent (T9424, Sigma-Aldrich) and converted to cDNA using the iScript cDNA synthesis kit (1708891, Bio-Rad). RNA expression analysis was performed by quantitative RT-PCR (LightCycler 480 system, Roche) using primer sets (Merck Millipore) designed with the Primer Designing Tool from NCBI (https://www.ncbi.nlm.nih.gov/tools/primer-blast/). For comparison of gene expression between metabolic states, expression levels (normalized to the geometric mean of the housekeeping genes GAPDH and ACTB) were expressed relative to control conditions via ΔCT method. Details on primers used are available in Supplementary Table 36.

### Protein isolation and immunoblotting

Cell lysates were prepared in RIPA lysis buffer (89900, Thermo Scientific) with protease and phosphatase inhibitor cocktail (78440, Thermo Fisher Scientific) and Benzonase Nuclease (E1014, Millipore). Lysates were then centrifuged at 4°C for 10 minutes at 12,000 x g and supernatant was used for analysis. Protein content was measured using Pierce BCA Protein Assay Kit (23227, Pierce). Lysates were separated by SDS-PAGE under reducing conditions, transferred to a PVDF membrane (1704157,Bio-Rad), and analyzed by immunoblotting. Primary antibodies used were goat anti-VE-Cadherin (AF1002, R&D Systems), rabbit anti-TAGLN (ab14106, abcam), rabbit anti-CD44 (GTX102111, GeneTex), rabbit anti-SNAIL (3879, Cell Signaling Technology), rabbit anti-ZEB2 (14026-1-AP Proteintech), rabbit anti-VCAM-1 (13662, Cell Signaling Technology), and mouse anti-Vinculin (V9131, Sigma-Aldrich). Appropriate secondary antibodies were polyclonal goat anti-mouse immunoglobulins/HRP (P044701-2, Agilent) and polyclonal goat anti-rabbit immunoglobulins/HRP (P044801-2, Agilent). Signal was detected using the ECL system (34096, Thermo Scientific) and densitometric quantifications were done using the Fiji software (https://fiji.sc).

### Whole mount immunohistochemistry

Whole human subcutaneous adipose tissue and murine inguinal adipose tissue samples were fixed in 4% paraformaldehyde for 24 hours before being washed and maintained in PBS at 4 °C for a maximum of 2 weeks. Whole adipose tissue was incubated in protein blocking buffer (3% BSA and 1% Triton X-100 in PBS) for 2 hours at room temperature on a rocking platform. Subsequently, tissues were incubated in 1:100 primary antibodies diluted in protein blocking buffer overnight, washed with PBS, and incubated with 1:100 secondary antibodies or pre-labeled primary antibodies diluted in protein blocking buffer overnight. Tissues were then washed with PBS and counterstained with Hoechst (1:500) prior to mounting. The following stains and antibodies were used: BODIPY 493/503 (D3922, Invitrogen), rabbit anti-RBP7 (HPA034749, Sigma-Aldrich), rabbit anti-FABP4 (710189, Invitrogen), rabbit anti-CA4 (158-10472-R039-100, SinoBiological), rat anti-GPIHBP1 (a generous gift from Stephen G. Young, University of California, Los Angeles, USA), rabbit anti-VCAM-1 (13662, Cell Signaling Technology), rabbit anti-ACKR1 (HPA017672, Sigma-Aldrich), mouse anti-CD36 (MA5-14112, Invitrogen), mouse anti-GJA5 (sc-365107, Santa Cruz Biotechnology), rabbit anti-MYH11 (ab53219, abcam), rabbit anti-alpha-SMA (19245, Cell Signaling Technology), rabbit anti-PDGFRB (MA5-15143, Invitrogen), rabbit anti-NEGR1 (HPA011894, Sigma-Aldrich), mouse anti-DCN (H00001634-M01, Abnova), rabbit anti-KAZN (PA5-56925, Invitrogen), rabbit anti-PCLO (HPA015858, Sigma-Aldrich), mouse anti-CLDN5 (35-2500, Invitrogen), mouse anti-HLA-DR, DP, DQ (555556, BD Pharmingen), rabbit anti-IRF3 (11904, Cell Signaling Technology), rabbit anti-TAGLN (ab14106, abcam), mouse anti-NG2 (14-6504-80, Invitrogen), rat anti-PLVAP (MECA-32, DHSB), affinity purified anti-AQP1 (2353AP), rabbit anti-ISG15 (2743, Cell Signaling Technology), rat anti-CD31 (557355, BD Pharmingen), rabbit anti-CD31 (NB100-2284, Novus Biologicals), recombinant human IgG1 anti-CD31-APC (130-117-226, Miltenyi Biotec), donkey anti-mouse Alexa Fluor 488 (A-21202, Invitrogen), goat anti-rat Alexa Fluor 488 (ab150157, abcam), and goat anti-rabbit Alexa Fluor 488 (A-11008, Invitrogen). Z-stack images were acquired using a LSM 900 inverted confocal microscope equipped with four diode lasers (405nm, 488nm, 561nm, and 640nm) and an Airyscan2 detector. Images were either acquired using a 10x objective (PIn-Neofluar, NA: 0,3, working distance: 5,2mm) or a 40x objective (PIn-Neofluar, NA: 1,30, working distance: 0,21mm).

### Histology, immunohistochemistry, and image analysis

Paraffin embedded human as well as murine adipose tissue samples from *Zeb2*^ΔEC^ and *Twist1*^ΔEC^ mice, and their control counterparts were sectioned using an automated microtome HM 355S (Epredia) into 5 μm sections.

### H&E

Sections were gradually deparaffinized using Xylene (28973.363, VWR Chemicals), graded ethanol, and water. Sections were then stained with Mayer’s Hemalum solution (1.09249.0500, Sigma-Aldrich), washed, and stained with 1% Eosin Y (alcoholic) (RBC-0201-00A, CellPath). Sections were then dehydrated in graded ethanol and Neoclear before mounting using Eukitt Quick-hardening mounting medium (03989, Sigma-Aldrich).

### Picrosirius red

Sections were deparaffinized overnight in xylene and rehydrated in graded ethanol solutions. Staining was performed using a commercial picrosirius red staining kit (24901-500, Polysciences) according to the manufacturer’s protocol. Subsequently, sections were dehydrated in graded ethanol, cleared in xylene, and mounted. Tissue sections were an Olympus light microscope with identical microscope and camera settings for different groups and quantitative assessment of collagen staining was performed using Fiji (v.2.14.0).

### Immunohistochemistry

Paraffin embedded sections were deparaffinized in Xylene and rehydrated in graded ethanol then subjected to antigen retrieval in TEG buffer (0,01M Tris (T1503, Sigma-Aldrich), 0,5mM EGTA (E4378, Sigma-Aldrich) in double distilled water, pH 9) in a steamer for 60 minutes. Sections were allowed to cool to room temperature after which sections were washed and incubated with protein blocking buffer overnight (3% BSA, 0,1% Triton X-100 in PBS) for one hour at room temperature. Sections were then incubated with primary antibodies diluted in protein blocking buffer at 4 degrees overnight, washed with PBS, and incubated with 1:1000 secondary antibodies diluted in protein blocking buffer for 2 hours at room temperature. Tissues were then washed with PBS and counterstained with Hoechst (1:5000) prior to mounting. The following primary antibodies were used to stain human sections: 1:800 mouse anti-CD31 (3528, Cell Signaling Technology), 1:100 rabbit anti-CD31 (NB100-2284, Novus Biologicals), 1:200 mouse anti-alpha smooth muscle actin (M085129, Agilent), 1:400 rabbit anti-FABP4 (710189, Invitrogen), 1:100 rabbit anti-COL3A1 (NB120-6580, Novus Biologicals), 1:500 rabbit anti-VCAN (MA5-42721, Invitrogen), 1:400 rabbit anti-COL1A1 (72026, Cell Signaling Technology), 1:200 rabbit anti-VCAM1 (13662, Cell Signaling Technology), 1:1000 rabbit anti-CD74 (77274, Cell Signaling Technology) and 1:200 mouse anti-CLDN5 (35-2500, Invitrogen). The following primary antibodies were used to stain mouse sections: 1:100 rabbit anti-CD31 (NB100-2284, Novus Biologicals), 1:150 goat anti-VE-Cadherin (AF1002, R&D Systems), and 1:150 rabbit anti-F4/80 (70076, Cell Signaling Technology). Secondary antibodies used were donkey anti-goat AF647 (A21447, Invitrogen), donkey anti-goat AF568 (A11057, Invitrogen), donkey anti-mouse AF568 (A10037, Invitrogen), donkey anti-rabbit Alexa Fluor 555 (A31572, Invitrogen), donkey anti-mouse Alexa Fluor 555 (A-31570, Invitrogen), donkey anti-goat AF488 (ab150129, abcam), donkey anti-mouse AF488 (A21202, Invitrogen), and donkey anti-rabbit Alexa Fluor 488 (A-21206, Invitrogen). Z-stack images were acquired using a LSM 900 inverted confocal microscope and exported using the Zeiss software. Image analysis was performed using Fiji (v.2.14.0).

### Image acquisition and analysis

For the quantification of adipocyte size, five random fields were acquired per section using an Olympus light microscope and adipocyte size was measured using the *Adiposoft*^132^ plugin in Fiji. Erroneous traces of adipocytes were manually corrected and adipocyte size from the five random fields was averaged. Tissue fibrosis was quantified based on picrosirius red stained fraction, which was quantified from five random fields per section following thresholding in Fiji. Macrophage tissue infiltration was assessed based on the percentage of F4/80^+^ cells of all nucleated cells. Nuclei were counted with a threshold-based mask for Hoechst and F4/80^+^ cells were manually counted and normalized to the number of Hoechst^+^ nuclei. The expression of endothelial cells (CD31^+^) or mural cells (aSMA^+^) of different proteins was quantified in a vascular cell-based ROI manner from five random fields. First, a threshold-based mask was applied to the relevant vascular marker (CD31/aSMA), generating ROIs of which the area was measured. Next, channels were split, the vessel-based ROIs were applied to the channel containing the relevant marker of interest and the mean gray value (MGV) of the signal was calculated within the ROIs. Ratios were calculated by normalizing the average MGV of the marker of interest to the average area of the relevant vessel marker. Monocyte adhesion onto cultured endothelial cells was measured by manually counting CellTracker Green CMFDA^+^ cells then normalizing their count to that of Hoechst^+^ cells.

### Conventional flow cytometry

Human SAT stromovascular cells were thawed and resuspended in a total of 10ml ECGM2 media, spun down, and the cell pellet was resuspended in wash buffer then incubated with CD31 human Microbeads and FcR Blocking Reagent according to the manufacturer’s instructions for 20 minutes. Cells were then washed with wash buffer and passed through an activated MS column. Magnetically labeled cells were immediately flushed out by firmly pushing the plunger in the column. Isolated cells were spun down, resuspended in 100ul wash buffer, and stained with Hoechst (1:500), CD45-PE-Vio770 (1:50) (130-110-634, Miltenyi Biotec), CD31-APC (1:50) (130-117-226, Miltenyi Biotec), CD34-PE (1:50) (12-0349-41, Invitrogen), and PDGFRA-PE (1:50) (A15785, Invitrogen) for 30 minutes at 4 °C. For murine inguinal adipose tissue stromovascular cells, cells were similarly thawed, resuspended in 100ul wash buffer and stained with CD31-PE (1:50) (130-102- 971, Miltenyi Biotec), CD102-APC (1:50) (130-112-030, Miltenyi Biotec), CD45-PE-Cy7 (1:100) (25-045-81, Invitrogen), and PDGFRA-BV421 (135923, BioLegend). Cells were then washed twice with wash buffer and murine cells were subsequently stained with 1:10,000 SYTOX Green (S7020, Invitrogen) directly before acquiring data. Stained and unstained cells as well as stained compensation beads were acquired on a NovoCyte 3000 flow cytometer equipped with three lasers (405nm, 488nm, and 640 nm) and 13 fluorescence detectors using the NovoExpress (v. 1.6.2) software (Agilent, Santa Clara, CA). Data analysis was carried out using FlowJo (v. 10.10.0). Detailed gating strategies of human CD45^-^CD31^+^PDGFRA^+^ and CD31^+^CD34^+^CD45^+^ cells are presented in Supplementary Fig. 2a. Detailed gating strategy of murine CD31^+^CD102^+^CD45^+^ and CD31^+^ CD102^+^PDGFRA^+^cells are presented in Supplementary Fig. 3e,f.

### Full-spectrum flow cytometry

For spectral flow cytometry, human SAT and murine inguinal adipose tissue stromovascular cells were thawed in RPMI 1640 containing 100μg/mL DNAse I (100-0762, StemCell) for 15 minutes at room temperature. For human cells, one million cells were then stained using Live/Dead fixable blue (L23105, Invitrogen) in 1X PBS for 20 minutes at room temperature. Subsequently, cells were incubated with an antibody mix (1:200 HAL-DR-BUV615 (G46-6, 751142, BD Optibuild), 1:600 CD206-BUV661 (19.2, 741641, BD Optibuild), 1:300 CD19-BUV805 (HIB19, 742007, BD Optibuild), 1:100 CD127-BV421 (HIL-7R-M21, 562437, BD Biosciences), 1:200 CD11b-BV570 (ICRF44, 301325, Biolegend), 1:200 CD3-BV605 (SK7, 563219, BD Horizon), 1:100 CCR2-BV786 (1D9, 747855, BD Optibuild), 1:50 CD144-PE (REA199, 130-118-358, Miltenyi Biotec), 1:50 CD31-APC (REA1028, 130-117-226, Miltenyi Biotec), 1:100 CD163-R718 (Mac2-158, 568185, BD Horizon), and 1:1000 CD45-APC-Vio770 (REA747, 130-110-635, Miltenyi Biotec)) containing human IgG (Privigen) and OligoBlock^133^ in FACS buffer (1mM EDTA pH 8.0, 2.5% fetal calf serum, 0.09% NaN3 (52300.0500, Ampliqon) in 1X PBS pH 7.4) at 4 degrees for 30 minutes. Finally, cells were intracellularly stained with 1:50 CD247-PerCP- eFluor710 (6B10.2, 46-247-42, Invitrogen) overnight at 4 degrees using the eBioscienceTM Foxp3/Transcription factor staining buffer set (00-5523-00, Invitrogen) according to the manufacturer’s instructions. Stained, unstained, and fluorescence minus one (FMO) cells were run on an ID7000 full spectrum flow cytometer equipped with five lasers (355nm, 405nm, 488nm, 561nm, and 637nm) and 147 fluorescence detectors (SONY Biotechnologies, San Jose, CA). Unmixing was performed using UltraComp eBeadsTM Plus compensation beads (01-3333-41, Invitrogen) and using the ID7000 Software version 2.0.2 (SONY Biotechnologies, San Jose, CA) utilizing the Weighted Least Squares Method (WLSM) algorithm for spectral unmixing. Subsequent analyses were performed in FlowJo (v. 10.10.0) using the PeacocQC^134^ (v. 1.5.0). Detailed gating strategies of human CD144^+^CD31^+^CD45^+^CD247^+^ and CD144^+^CD31^+^CD45^+^CD11b^+^CD163^+^CD206^+^ cells are presented in Supplementary Fig. 4e.

### ImageStream

Human subcutaneous adipose tissue stromovascular cells were thawed in 10ml supplemented ECGM2, washed in wash buffer, fixed in 4% PFA for 15 minutes at room temperature, then pre-permeabilized with 0,2% Triton X-100 for 15 minutes at room temperature. Cells were subsequently washed twice and stained with prelabeled antibodies at 4 degrees Celcius for one hour. Antibodies used were the following: 1:20 CD45- BB515 (564585, BD Horizon), 1:50 PDGFRA-biotin (130-115-236, Miltenyi Biotec), 1:50 CD31-APC, and 1:50 CD144-PE (130-118-358, Miltenyi Biotec). Cells were then washed twice and incubated with 1:25 BV421 streptavidin (563259, BD Horizon) at 4 degrees Celcius for 20 minutes. Cells were finally washed and data were acquired through the Amnis ImageStream imaging flow cytometry (Luminex) equipped with 405nm (100 mW), 488nm (170mW), 561nm (200mW), and 642nm (150mW) lasers and the brightfield set intensity was set to 619. Laser intensities were set to maximal values that do not saturate the camera. Camera gain was set to 1, sensitivity to 32, and images were acquired at 60X magnification. Data were analyzed using IDEAS 6.2 software (Amnis). Cells in focus were gated using the Gradient RMS feature, which measures the sharpness quality of an image by detecting large variations in pixel values in an image computed using the average gradient of a pixel normalized for variations in intensity levels. Cells were gated on area (sum of pixels within a mask in um^2^) and side scatter intensity, following which, single cells were then gated using the area and aspect ratio (the minor axis divided by the major axis). Specific gating strategies to identify CD144^+^CD31^+^CD45^+^ cells and CD144^+^CD31^+^PDGFRA^+^ cells are presented in Supplementary Fig. 2b.

## Data Availability

The raw sequencing data generated in this study is available at Gene expression omnibus (GEO) under the accession code GSE268904. Publicly available datasets were downloaded from their respective repositories; GEO database under the accession codes GSE176171, GSE129363, GSE155960, GSE128889, GSE241015, GSE235192, GSE241015, Single Cell Portal (SCP) with accession code SCP1903, and EMBL-EBI with accession code E-MTAB-12865. Tabula Sapiens data were retrieved from the Tabula Sapiens portal. Data from Massier et al. were obtained after contacting the authors.

## Code availability

The scripts utilized for the analysis done in this study can be accessed through GitHub on the following link; https://github.com/Kalucka-lab/SAT_Atlas).

### Statistics

Data are represented as mean ± s.e.m and statistical analyses were performed using GraphPad Prism v.10.2.2. Statistical significance between two groups was calculated using two-tailed one sample t test (for comparison to point-normalized data), unpaired two-tailed t test with Welch correction, or OneWay ANOVA. *P* values inferior to 0.05 were considered significant. Statistical tests used for all computational analyses are mentioned under each method section. Gene ontology analysis was performed using over-representation analysis based on hypergeometric distribution. Differential expression analysis across pseudotime was computed using Moran’s I statistics. For connectome analysis, significant interactions were calculated using permutation test. For all computational analyses, *P* values were adjusted after correcting for false discovery rate using Benjamini-Hochberg and was considered significant if inferior to 0.05. For the identification of druggable genome targets, gene expression specificity was assessed based on AUC for each gene calculated using receiver operating characteristic (roc) test.

## Author contribution

I.A., M.N.H., L.d.R., and J.K. conceptualized the project and experimental design. I.A., M.S., M.B.S., Jo.F., L.D., B.H., R.K.R., J.N.C., O.W., R.S., M.C., A.B.B., J.F., L.L. and J.K. performed the experiments. M.N.H. led the bioinformatics work, with analyses conducted by I.A., M.N.H., L.M., K.D., and S.W.; S.W. was responsible for spatial transcriptomics. B.T.A., W.D., S.S., A.D.T., A.B., H.H.T., M.v.H., L.C., P.E., C.E.H., J.H., A.L., M.E., E.D.R., A.E., L.M., M.R., N.M., M.B., K.K., R.A.F., B.N.S., and N.J. provided biological samples, computational data, reagents, and contributed to result interpretation. J.K. supervised the project, provided guidance, conceptualized the study, and secured funding. I.A., M.N.H., and J.K. wrote the manuscript, with figures prepared by I.A. All authors read, edited, and approved the final version of the manuscript.

## Acknowledgments

This work was supported by Lundbeckfonden (R307-2018-3667), Carlsbergfonden (CF19-0687), and Riisfort Fonden to J.K.; Novo Nordisk Foundation (NNF20OC0063268 to S.S., NNF21OC0071718 to L.L., NNF21OC0067146 to K.K., NNF21OC0067647 to R.A.F.); Swedish Research Council (#2019-02046) and Karolinska Institutet (2-189/2022 and 2020-00893) to C.E.H.; DFG-funded consortium SFB-Transregio 333, project ID 450149205 to J.H.; internal KU Leuven funding (C14/19/095) to A.L.; a fellowship from the Fonds voor Wetenschappelijk Onderzoek (FWO; 1157318N) to W.D.; and RC2 DK116691 to E.D.R. M.B. has received personal honoraria from Amgen, AstraZeneca, Bayer, Boehringer Ingelheim, Daiichi-Sankyo, Lilly, Novo Nordisk, Novartis, and Sanofi.

We thank Inger Merete S. Paulsen, Prof. Danny Huylebroeck, and Alice Maestri for technical assistance, provision of materials, and valuable discussions. We are also grateful to the study participants for their involvement.

---

**Extended Data Fig 1.**
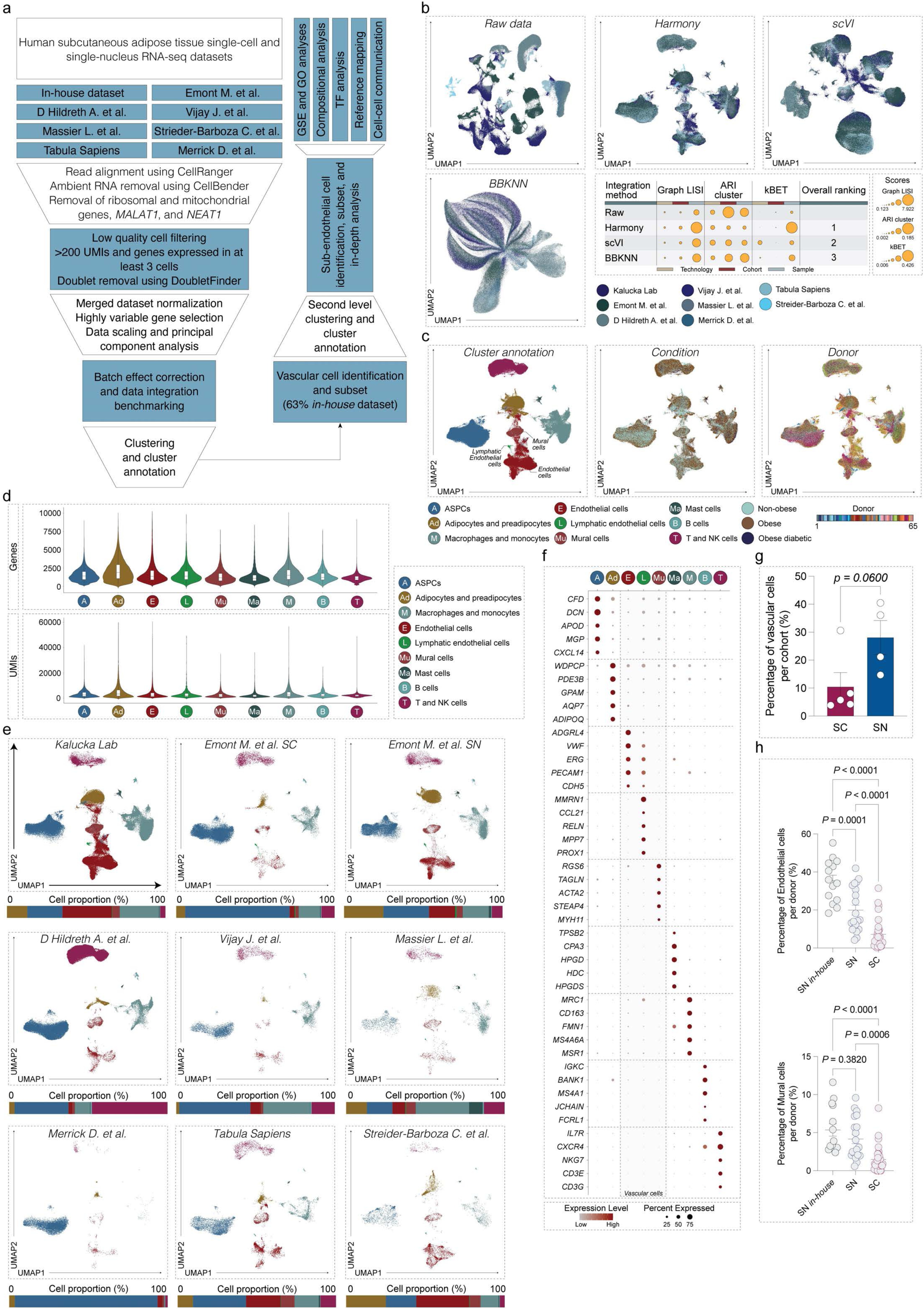
Construction of an integrated single-cell transcriptome atlas of human SAT reveals heterogeneous vascular cell populations. **a,** Schematic overview of the workflow used to construct the human subcutaneous adipose tissue (SAT) atlas and the downstream analyses performed. **b,** UMAP projections of 329,774 cells grouped by cohort, shown before (raw data) and after integration using *Harmony*, *scVI*, and *BBKNN*. Table (bottom right) summarizes Graph LISI, ARI, and kBET scores for integration performance, with overall method ranking. **c,** UMAP of Harmony-integrated SAT data, colored by cluster annotation, metabolic state, and donor (each donor represented by a distinct bar). **d,** Violin plots showing the number of UMIs and detected genes per cell type across the SAT atlas. **e**, UMAP projections split by cohort and technology (single-cell [SC] and single-nucleus [SN]). Bar graphs show the relative proportion of cell types as a percentage of all cells/nuclei per cohort. **f,** Dot plot of top marker gene expression for major cell populations. Dot size reflects the percentage of cells within a cluster expressing each gene; color indicates expression level (red: high, grey: low). **g**, Bar graph showing the proportion of vascular cells per cohort, split by technology. Each dot represents one cohort (n=4–5). **h**, Bar graphs showing the relative abundance of endothelial and mural cells as a percentage of all cells per donor, stratified by technology (SN and SC). SN samples are further split into *in-house* (n=14) and publicly available (n=20) datasets; SC, n= 31. Error bars represent mean ± s.e.m. *P* values were calculated using unpaired t-test (g) or one-way ANOVA (h). *P* < 0.05 was considered significant.

**Extended Data Fig 2:**
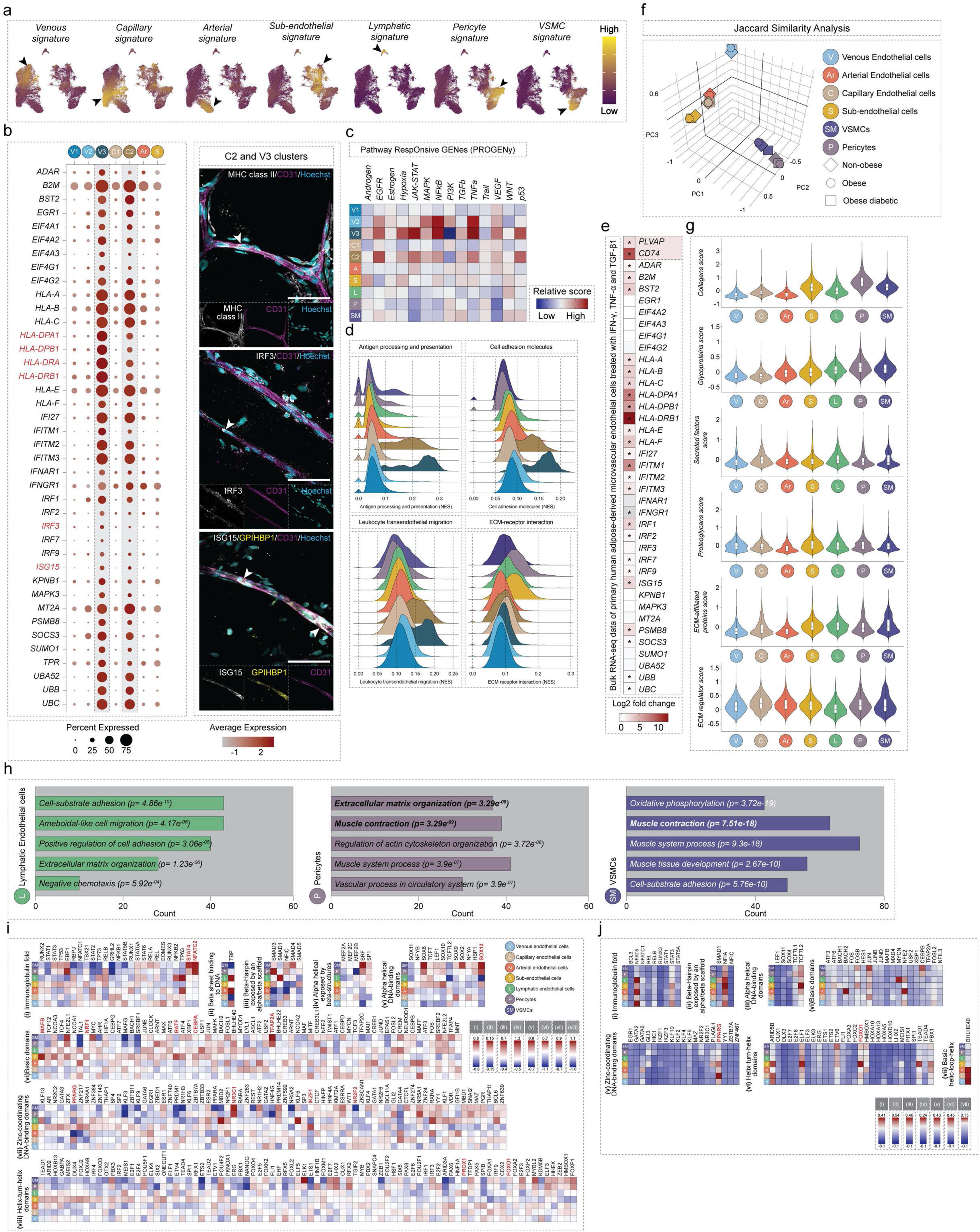
Transcriptomic and functional characterization of human SAT AdEC and mural cell populations. **a**, Feature plots showing gene signatures of distinct vascular cell populations in the human SAT atlas. Arrows highlight cells with enriched expression. Color scale: yellow, high expression; purple, low expression. **b**, *Left:* Dot plot illustrating the enrichment of interferon-related genes in AdECs from the V2 and C3 clusters. Dot size indicates the percentage of cells expressing each gene; color reflects expression level (red: high, grey: low). *Right:* Immunohistochemical staining of human SAT wholemounts showing protein-level expression of selected markers (MHC class II, IRF3, ISG15). White arrows indicate cells with pronounced expression. Scale bars: 50μm **c,** Heatmap showing pathway activity scores inferred using PROGENy (Pathway RespOnsive GENes). Colour scale: red, high activity; blue, low activity. **d,** Ridge plots showing normalized enrichment scores of immune- and ECM-related pathways, computed using scGSVA. **e,** Heatmap depicting the mean log₂ fold changes of interferon-related genes in AdECs treated with IFN-γ, TNF-α, and TGF-β1^15^. Asterisks indicate statistical significance (likelihood ratio test). Color scale: red, high expression; grey, low expression. **f**, Three-dimensional principal component analysis (PCA) based on pairwise Jaccard similarity coefficients of marker genes across vascular cell populations in the human SAT atlas. **g**, Violin plots showing the enrichment of matrisome gene signatures across vascular cell populations. **h**, Bar plots showing five representative Gene Ontology terms among the most significantly enriched pathways in lymphatic endothelial cells, pericytes, and vascular smooth muscle cells (VSMCs). The x-axis indicates the number of associated genes. Bolded GO terms have been highlighted in the text. **i**, Heatmaps showing inferred transcription factor (TF) activity, computed using *DoRothEA*, across endothelial subtypes (venous, capillary, arterial, lymphatic, and sub-AdECs), pericytes, and VSMCs. Color scale: red, high inferred activity; grey, low inferred activity. **j**, Heatmaps showing regulon activity of transcription factors, inferred using *pySCENIC*, across endothelial subtypes (venous, capillary, arterial, lymphatic, and sub-AdECs), pericytes, and VSMCs. Color scale: red, high regulon activity; grey, low regulon activity. TFs highlighted in red correspond to those discussed in the main text.

**Extended Data Fig 3:**
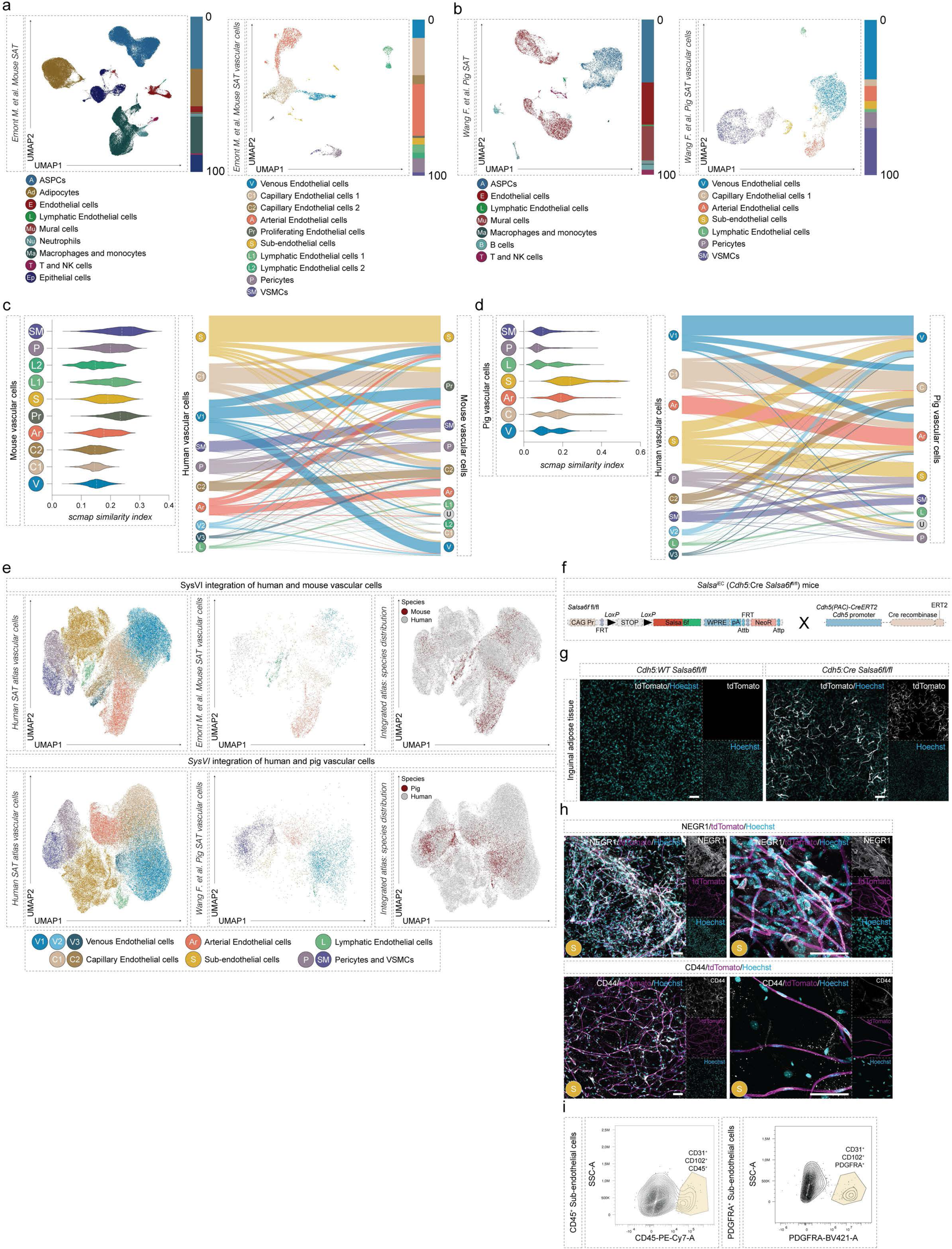
Human, murine, and porcine SAT vascular niches share conserved cell populations. **a**, UMAP projections of single-cell transcriptomics data of clusters formed by murine SAT (inguinal adipose tissue) cells *(left)* and subset vascular cells *(right)* comprising endothelial, lymphatic, and mural cells. Bar graphs show the relative proportion of cell types as a percentage of all cells/nuclei. **b**, UMAP projections of single-cell transcriptomics data of clusters formed by porcine SAT cells *(left)* and subsetted vascular cells *(right)* comprising endothelial, lymphatic, and mural cells. Bar graphs show the relative proportion of cell types as a percentage of all cells/nuclei. **c**, *Left*: Violin plot showing the *scmap* similarity index between human and murine vascular cell populations. *Right*: Riverplot showing the relationship between human and murine vascular cell clusters. **d**, *Left*: Violin plot showing the *scmap* similarity index between human and porcine vascular cell populations. *Right*: Riverplot showing the relationship between human and porcine vascular cell clusters. **e**, UMAP projections of Human SAT vascular cells integrated using *SysVI* with either the murine *(top)* or the porcine vascular cells (*bottom*), split or grouped by species. **f**, Schematic diagram detailing the generation of Salsa^iEC^ mice. **g**, Immunohistochemical validation of reporter mouse strains recombination following tamoxifen administration. **h**, Immunohistochemical validation of wholemount inguinal adipose tissue of reporter mouse strain for proteins encoded by marker genes of sub-AdECs population. **i**, Representative contour plots of flow cytometry experiments showing the occurrence of sub-AdECs expressing CD45 (CD144/VE-Cadherin^+^CD31^+^CD45^+^ cells) and PDGFRA (CD144/VE-Cadherin^+^CD31^+^PDGFRA^+^ cells) within murine inguinal adipose tissue SVF. Scale bars (g, h): 50 μm.

**Extended Data Fig. 4:**
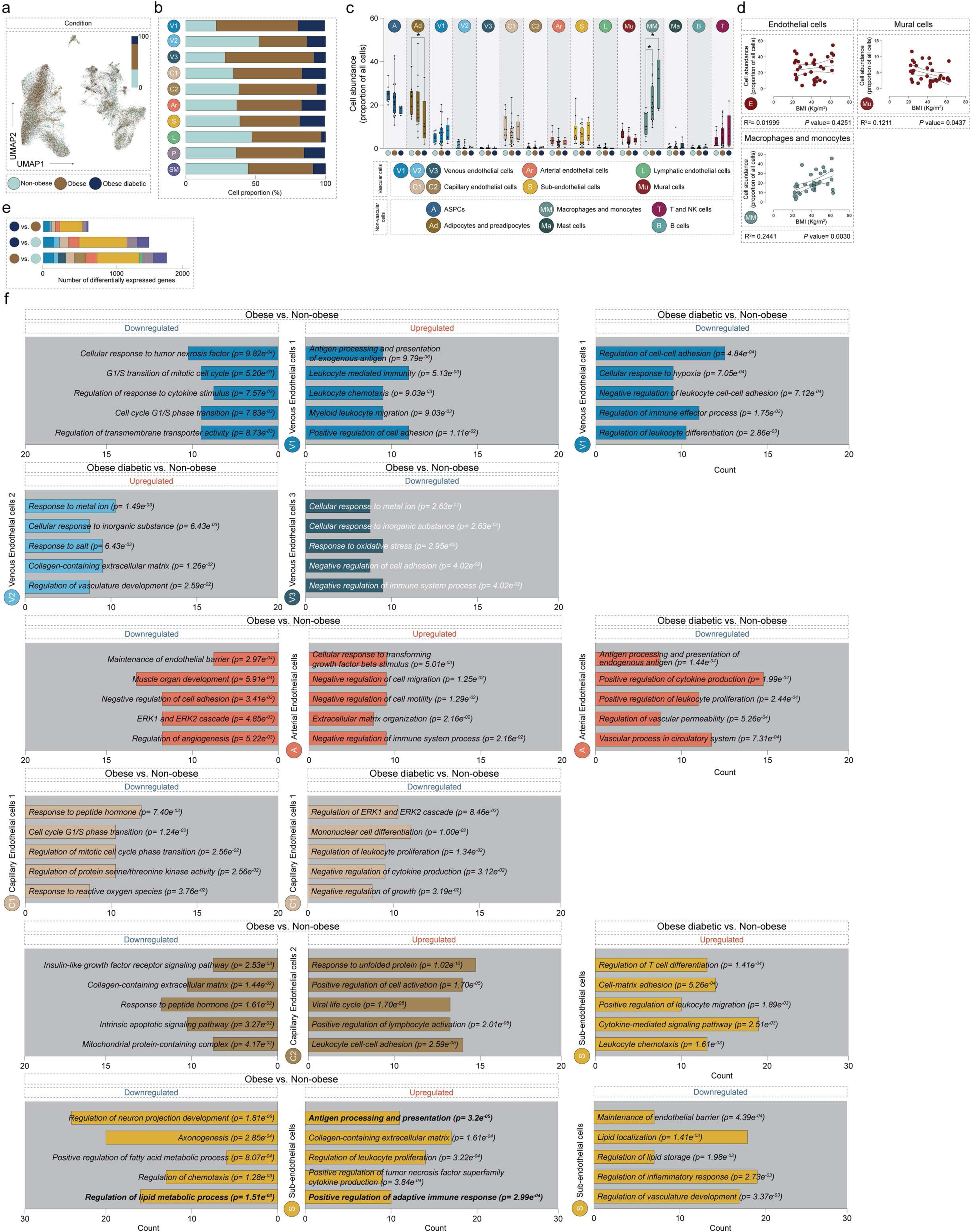
Canonical adipose tissue endothelial cells exhibit transcriptional changes associated with metabolic disease. **a**, UMAP projection of vascular cells from the human SAT atlas grouped by metabolic state. **b**, Proportion of each vascular cell population within each metabolic state. **c**, Bar plots showing the relative abundance of sequenced nuclei per cell type. **d**, Linear regression analysis showing the association between the abundance of sequenced nuclei from selected cell types (endothelial cells, mural cells, macrophages, and monocytes) and body mass index (BMI). **e**, Bar plot indicating the number of significant DEGs in each vascular cell population across three comparisons: Obese vs. Non-obese, Obese diabetic vs. Non-obese, and Obese diabetic vs. Obese. **f**, Bar plots displaying five representative GO terms among the most significantly upregulated and downregulated pathways in AdEC populations across the Obese vs. Non-obese and Obese diabetic vs. Non-obese comparisons. x-axis indicates gene count. Bolded GO terms have been highlighted in the text. Statistics: Statistical significance was assessed using *scCODA* for compositional analysis (c). For regression analysis (d), p-values were calculated using the t-distribution. In all cases, *P*< 0.05 was considered statistically significant.

**Extended Data Fig. 5:**
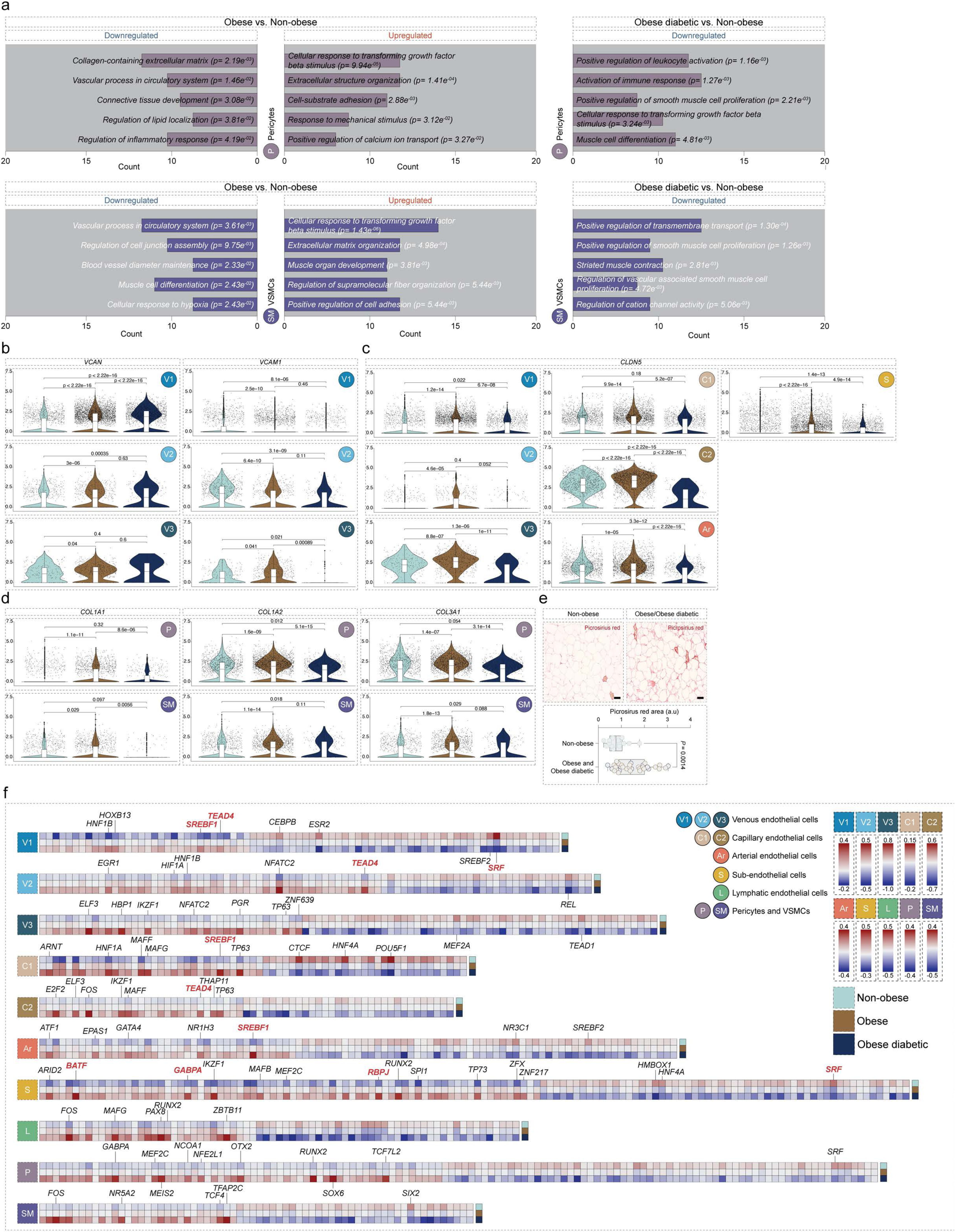
Mural cell populations in adipose tissue exhibit transcriptional signatures associated with metabolic disease. **a**, Bar plots displaying five representative GO terms among the most significantly upregulated and downregulated pathways in mural cell populations across two comparisons: Obese vs. Non-obese and Obese diabetic vs. Non-obese. x-axis indicates the number of associated genes. **b-d**, Violin plots depicting the expression levels of selected differentially expressed genes across metabolic states in selected cell populations (*VCAN*, *VCAM1*, *CLDN5*, *COL1A1*, *COL1A2* and *COL3A1*). **e**, *Top*: Representative images of picrosirius red staining of human subcutaneous adipose tissue sections. *Bottom*: Quantification of the picrosirus red area (Non-obese; n=14, Obese; n=22, and Obese diabetic; n=10). **f,** Heatmaps depicting the inferred enrichment of transcription factor activity in each vascular cell population and in each metabolic state as inferred by *deCoupleR*. Only transcription factors exhibiting progressive trends towards increased or decreased activity in both metabolic disease states are presented. Color scale: red, high expression; blue, low expression. Transcription factors highlighted in red correspond to those discussed in the main text. In violin plots, statistical significance was assessed using Kruskal-Wallis test, followed by Wilcoxon rank-sum tests for pairwise comparisons. In box plots, error bars represent mean ± s.e.m and *P* values were either calculated using unpaired t test. In all cases, *P*< 0.05 was considered statistically significant.

**Extended Data Fig. 6:**
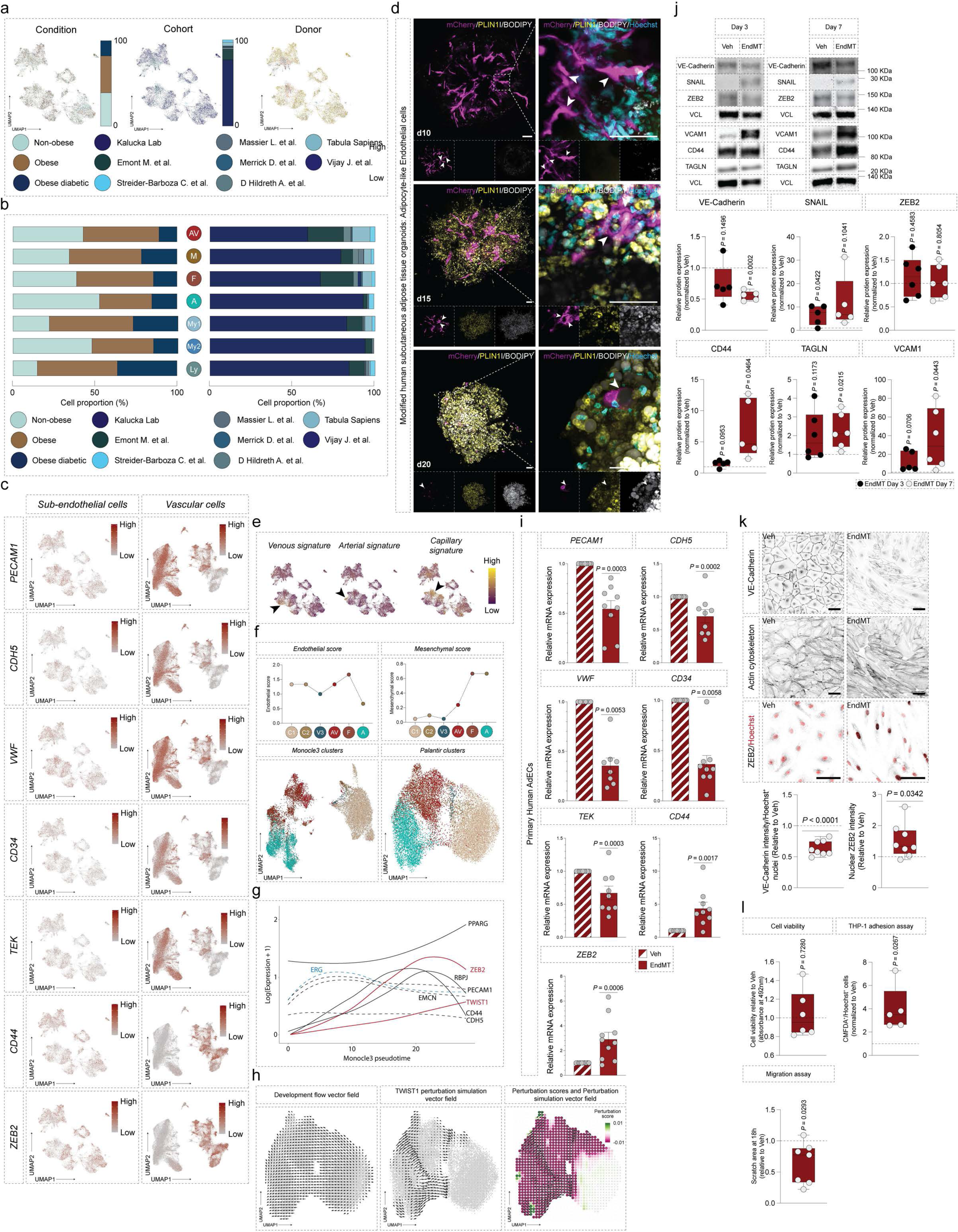
Sub-endothelial cells occur in non-obese donors, donors with obesity, and donors with obesity compounded by T2D and share gene signatures of canonical vascular cells. **a**, UMAP projections of sub-AdECs from the human SAT atlas, grouped by metabolic state (Condition), cohort, and individual donor. Sub-AdECs were detected in donors across all metabolic states, including Non-obese, Obese, and Obese diabetic. Bar graphs show the relative proportion of cells as a percentage of all cells/nuclei**. b,** Bar plots showing the distribution of each sub-AdEC subpopulation across the three metabolic states (Non-obese, Obese, and Obese diabetic, *left*) and eight donor cohorts (*right*). **c,** *Left*: Feature plots showing expression of endothelial markers (*PECAM1*, *CDH5*, *VWF*, *CD34, TEK*) and mesenchymal markers (*CD44*, *ZEB2*) in sub-AdECs. *Right*: Corresponding expression of the same genes across vascular cell populations in the SAT atlas. Color scale indicates gene expression level (red: high, grey: low). **d**, Immunofluorescent staining of 3D-differentiated human adipose tissue organoids for tdTomato (magenta, ECs), PLIN1 (yellow; marker of the adipocyte-like endothelial cell population), BODIPY (white, lipid droplets), and Hoechst (cyan, nuclei). White arrowheads indicate tdTomato⁺ endothelial cells. **e**, Feature plots showing the expression of gene signatures from canonical vascular cell populations within sub-AdECs. Arrows indicate cells with enriched expression. Color scale: yellow, high expression; purple, low expression. **f**, *Top*: Median endothelial and mesenchymal scores for selected endothelial cell populations, ordered by average *Monocle3* pseudotime values. *Bottom*: UMAP projection of *in-house* single-nucleus RNA-seq data showing the cell embeddings used to infer pseudotime trajectories using either *Monocle3* or *Palantir*. **g**, Variation in gene expression (log(Expression + 1)) of selected genes and transcription factors along the *Monocle3*-inferred pseudotime trajectory. **h**, Force-directed graphs illustrating sub-AdEC transition from canonical AdECs. Shown are the original developmental flow vector field, the simulated TWIST1 perturbation vector field, and the associated perturbation scores and resulting flow field. Color scale indicates perturbation scores. **i**, Quantitative RT–PCR analysis of selected endothelial and mesenchymal genes in an *in vitro* model of EndMT (n=8–10). **j**, *Top*: Representative Western blots of VE-Cadherin, the EndMT transcription factors: SNAIL and ZEB2, the mesenchymal proteins CD44 and TAGLN, and the vascular adhesion molecule VCAM1 in vehicle-treated and EndMT-induced primary human AdECs at 3- and 7-days post-induction (n=5–6). *Bottom*: Quantification of VE-Cadherin, SNAIL, ZEB2, VCAM1, CD44, and TAGLN protein levels. **k**, *Top*: Immunofluorescent images of primary human AdECs stained for VE-Cadherin, actin cytoskeleton, and ZEB2 (gray), with Hoechst counterstaining of nuclei, shown under vehicle-treated and EndMT-induced conditions. *Bottom*: Quantification of total VE-Cadherin intensity and nuclear ZEB2 intensity in EndMT-induced cells compared to vehicle-treated controls. **l**, Cell viability of EndMT-induced AdECs at day 7, normalized to vehicle-treated cells (n=6); Quantification of CMFDA-labeled THP-1 cell binding to EndMT-induced AdECs (normalized to Hoechst⁺ nuclei and vehicle-treated controls, n=5); Scratch area quantification at 18 hours post-wounding in EndMT-induced AdECs at day 7 (normalized to vehicle-treated controls, n=7). Scale bars (d,k): 50μm. Statistics: In bar graphs, error bars represent mean ± s.e.m. *P* values were calculated using a one-sample t-test (panels i–l). *P* < 0.05 was considered statistically significant.

**Extended Data Fig. 7:**
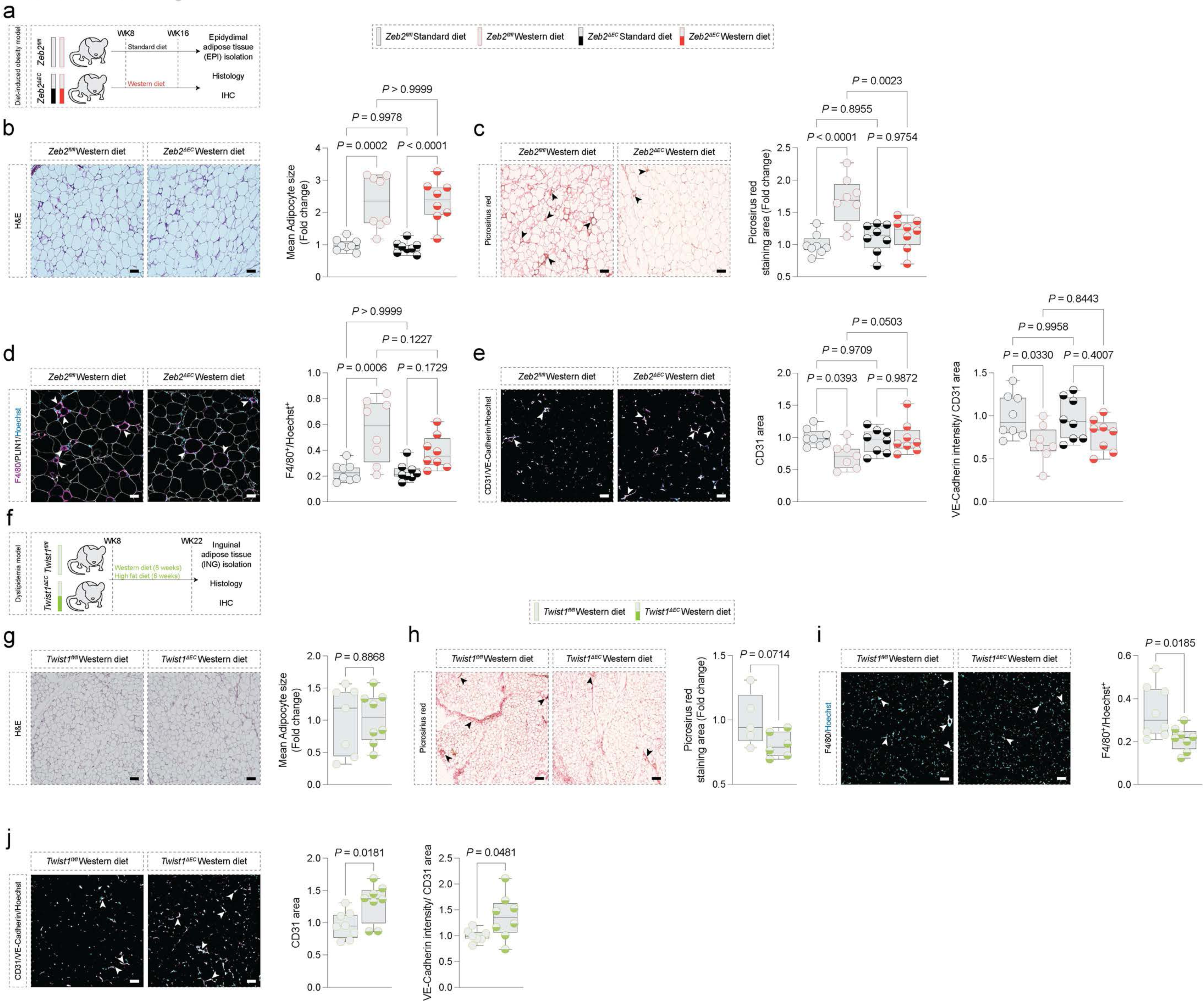
Inhibition of EndMT reduces adipose tissue fibrosis, inflammation, and endothelial junctional remodeling in metabolic disease models. **a,** Schematic illustrating the experimental design for inducing obesity in *Zeb2*^fl/fl^ and *Zeb2*^ΔEC^ mice via Western diet (WD) feeding for 8 weeks. **b,** *Left*: Representative H&E-stained sections; *Right*: Adipocyte size quantification in epididymal adipose tissue from *Zeb2*^fl/fl^ and *Zeb2*^ΔEC^ mice fed chow or WD (n=8). **c,** *Left*: Representative Picrosirius Red-stained sections; *Right*: quantification of collagen deposition in *Zeb2*^fl/fl^ and *Zeb2*^ΔEC^ mice (n=8). Black arrowheads highlight regions with pronounced Picrosirius Red staining. **d,** *Left*: Representative immunohistochemical staining for the macrophage marker F4/80; *Right*: corresponding quantification (normalized to Hoechst⁺ nuclei per image) (n= 8). White arrowheads indicate areas enriched in F4/80⁺ cells. **e,** *Left*: Representative images for the endothelial marker CD31 and the junctional protein VE-Cadherin; *Right:* quantification of total CD31⁺ area per image and VE-Cadherin intensity normalized to CD31⁺ area (n=7–8). White arrowheads indicate CD31⁺ vessels with pronounced VE-Cadherin expression. **f,** Schematic illustrating the experimental design for inducing dyslipidemia in *Twist1*^fl/fl^ and *Twist1*^ΔEC^ mice via Western/high-fat diet feeding for 14 weeks. **g,** *Left:* Representative H&E-stained sections; *Right*: adipocyte size quantification in inguinal adipose tissue of *Twist1*^fl/fl^ and *Twist1*^ΔEC^ mice (n=7–8). **h,** *Left:* Representative Picrosirius Red-stained sections; *Right*: quantification of collagen deposition in *Twist1*^fl/fl^ and *Twist1*^ΔEC^ mice (n = 5–6). Black arrowheads highlight regions with pronounced Picrosirius Red staining. **i,** *Left:* Representative F4/80 staining; *Right:* quantification of macrophage infiltration (normalized to Hoechst⁺ nuclei) (n = 8). White arrowheads indicate areas enriched in F4/80⁺ cells. **j,** *Left:* Representative staining for CD31 and VE-Cadherin; *Right*: quantification of CD31⁺ area and VE-Cadherin intensity (n=7–8). White arrowheads indicate CD31⁺ vessels with pronounced VE-Cadherin expression. Scale bars: 50μm Statistics: In bar graphs, error bars represent mean ± s.e.m. *P* values were calculated using unpaired t-tests or one-way ANOVA. *P* < 0.05 was considered statistically significant.

**Extended Data Fig. 8:**
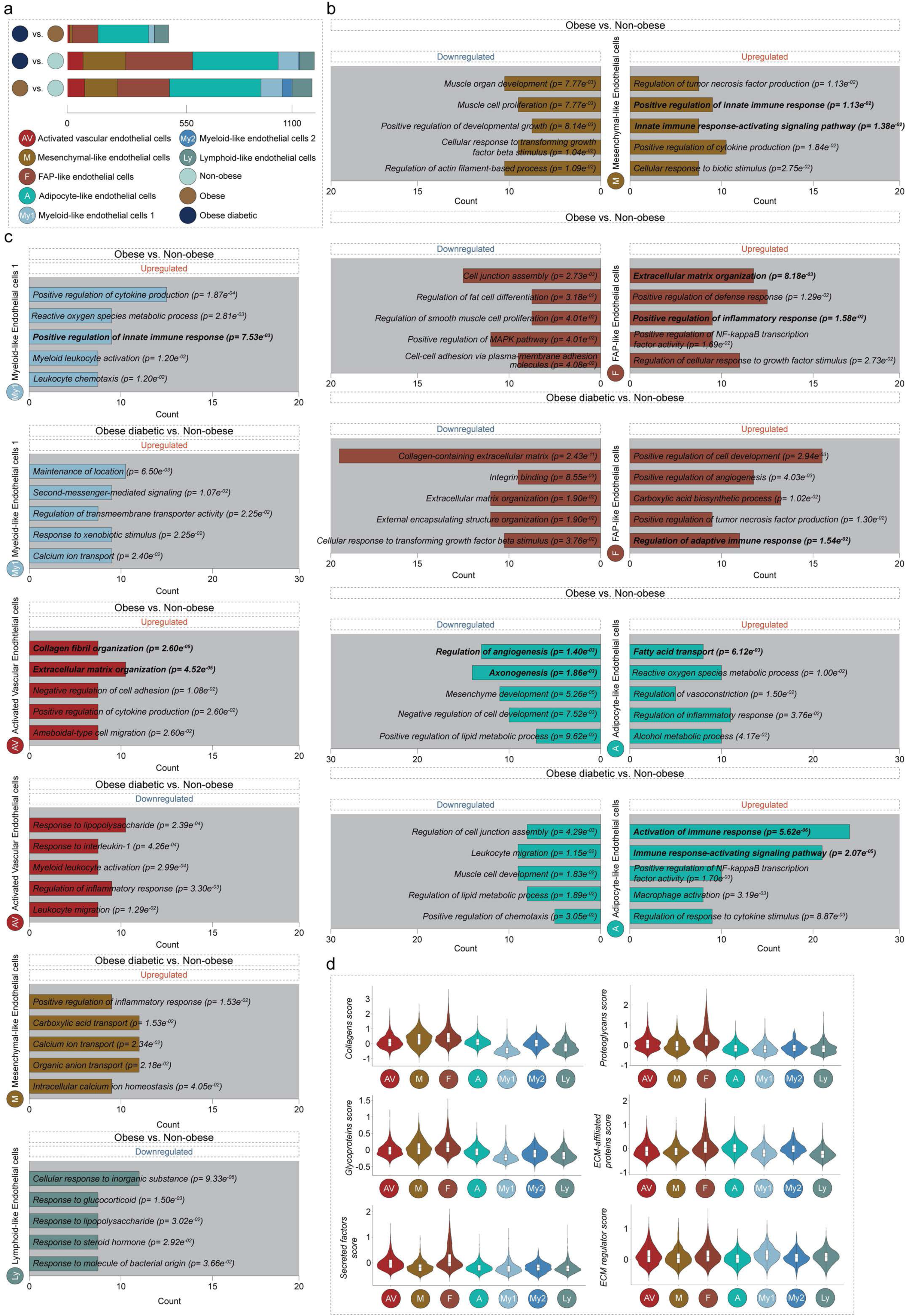
Sub-endothelial cells exhibit transcriptomic features associated with metabolic disease. a, Bar plot showing the number of significant DEGs in each sub-AdEC subpopulation across three comparisons: Obese vs. Non-obese, Obese diabetic vs. Non-obese, and Obese diabetic vs. Obese. **b, c,** Bar plots displaying five representative GO terms among the most significantly upregulated and downregulated pathways in sub-AdEC subpopulations for the Obese vs. Non-obese and Obese diabetic vs. Non-obese comparisons. x-axis indicates gene count. Bolded GO terms have been highlighted in the text. **d,** Violin plots showing the enrichment of matrisome gene signatures across sub-AdEC subpopulations.

**Extended Data Fig. 9:**
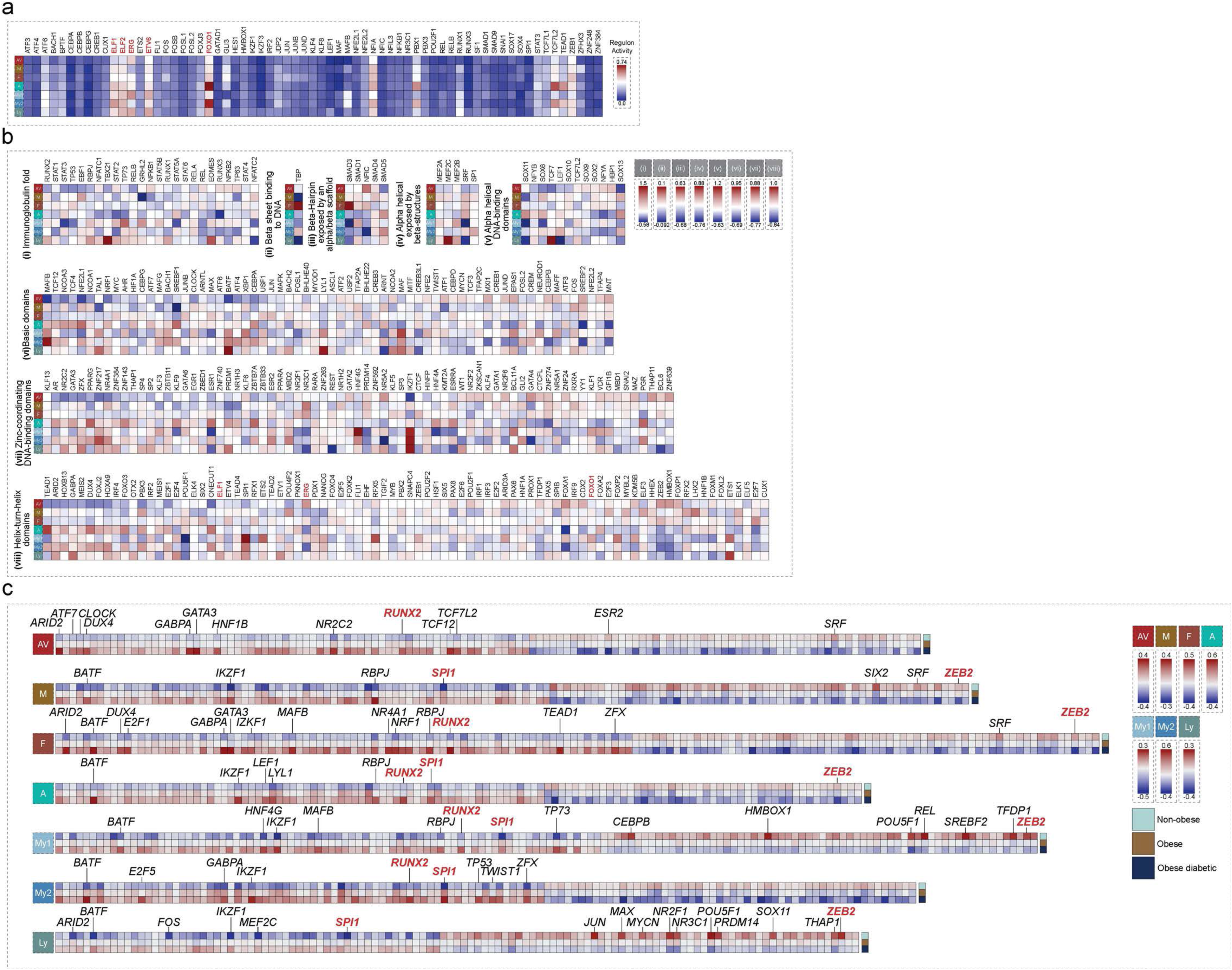
Sub-endothelial cells exhibit metabolic state-associated alteration in transcription factor repertoire activity. **a,b**, Heatmaps showing the inferred enrichment of transcription factor activity in each sub-AdECs subpopulation in the human SAT atlas independent of the metabolic state using *pySCENIC* in a and *deCoupleR* in b. **c**, Heatmap showing the inferred enrichment of transcription factor activity in each sub-AdECs subpopulation in the human SAT atlas across the three metabolic states; non-obese, obese, and obese diabetic. Transcription factors highlighted in red correspond to those discussed in the main text.

**Extended Data Fig. 10:**
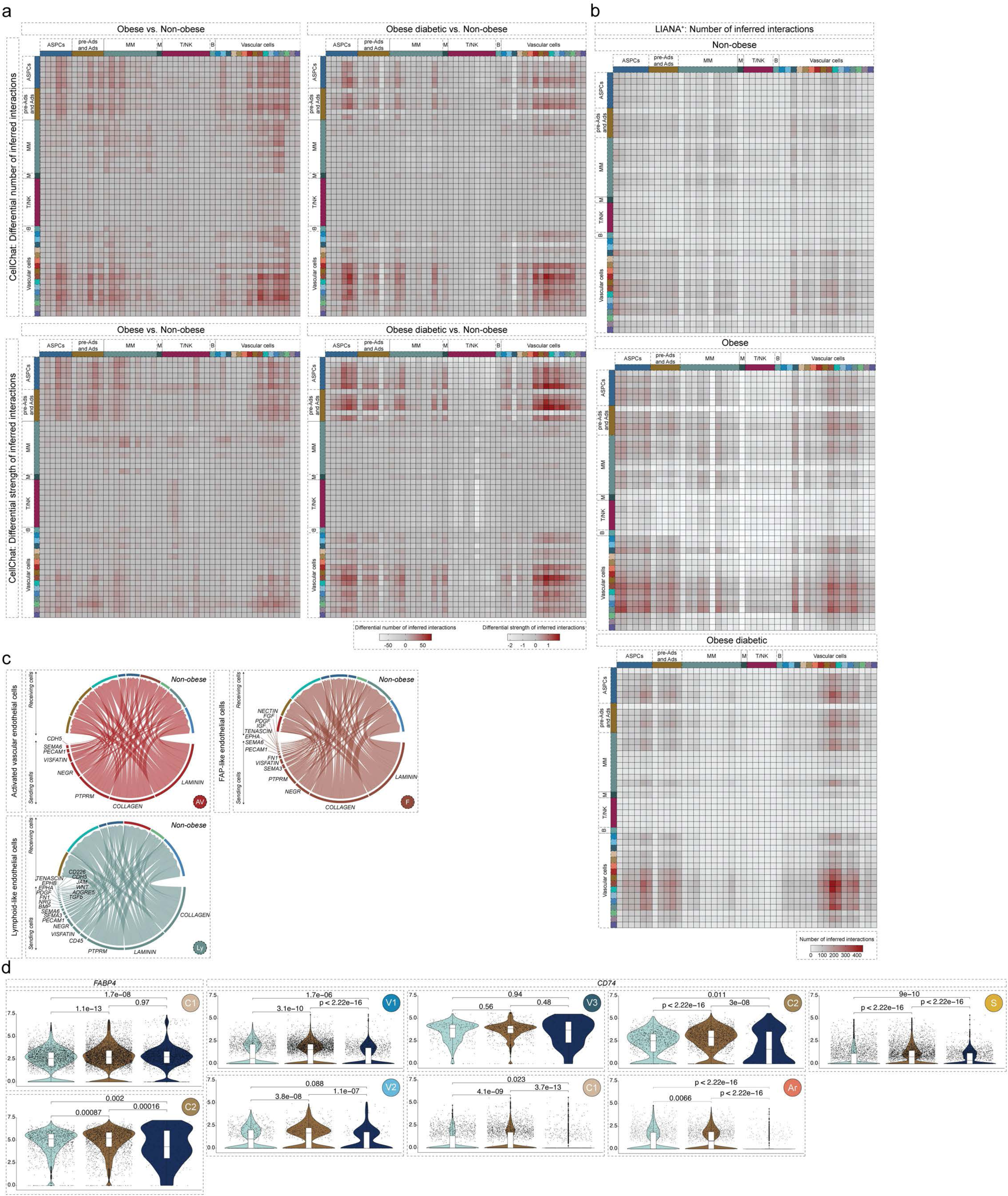
Sub-endothelial cells are implicated in pronounced intercellular communication. **a**, Heatmaps depicting the differential number and strength of interactions in human SAT (in the two comparisons, Obese vs. Non-obese and Obese diabetic vs. Non-obese) with cell types clustered at a high granularity as inferred by *CellChat*. **b**, Heatmaps depicting the number of interactions in human SAT (in the three metabolic states; Non-obese, Obese, and Obese diabetic) with cell types clustered at a high granularity as inferred by *LIANA^+^*. **c**, Chord diagrams depicting the results of *CellChat* inferred ligand–receptor analysis for significant signaling pathways between the three sub-AdECs populations with the highest differential number and strength of interactions (AV, F, and Ly) and their main interacting partners (receiving cells) in the non-obese metabolic state. **d**, Violin plots depicting the expression levels of *FABP4* and *CD74*, inferred genomic protein-coding genes, in selected endothelial cell populations. In violin plots, statistical significance was assessed using Kruskal-Wallis test, followed by Wilcoxon rank-sum tests for pairwise comparisons.

**Supplementary Fig 1.**
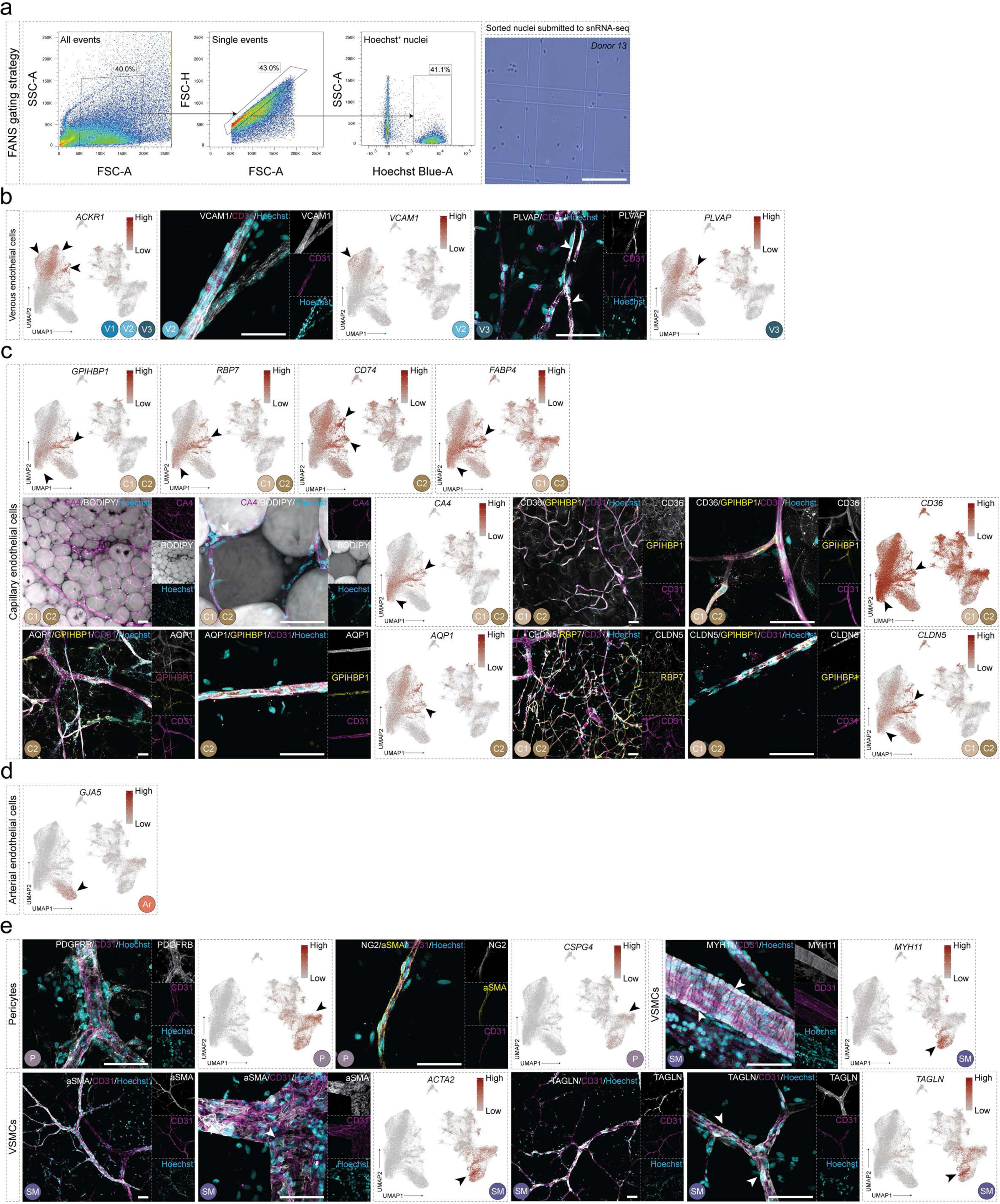
Gating strategies for nuclei sorting and immunohistochemical validation of marker gene expression in human adipose tissue wholemounts. **a**, *Left*: Scatter plots showing the gating strategy for Fluorescence-Activated Nuclei Sorting (FANS) from human subcutaneous adipose tissue (SAT). Nuclei were gated based on size (FSC/SSC), and Hoechst⁺ singlets were isolated for 10x Genomics library preparation. *Right*: Representative image of isolated nuclei stained with trypan blue in a counting chamber, showing preserved morphology and membrane integrity. **b**, *Left*: Feature plots showing gene expression of selected markers, including *ACKR1* (stained in Fig. 1c). Black arrowheads denote populations with increased gene expression. *Right*: Representative high-magnification images of immunohistochemical staining for venous (V2, V3) AdEC markers. White arrowheads mark ECs positive for the target protein. **c**, *Top*: Feature plots of *GPHB5*, *RBP7*, *CD74*, and *FABP4*, corresponding to markers shown in Fig. 1d. *Left*: Representative immunohistochemical images for capillary (C1, C2) AdEC markers. White arrowheads mark ECs positive for the target protein. *Right*: Feature plots of the same genes, with black arrowheads denoting populations with increased gene expression. **d**, Feature plots of *GJA5*, corresponding to immunofluorescence in Fig. 1e. **e**, *Left*: Representative immunohistochemical images for pericyte (P) and vascular smooth muscle (SM) markers. White arrowheads mark cells positive for the target protein. *Right*: Feature plots of the corresponding genes. In all feature plots, color scale indicates gene expression level (red: high, grey: low); Scale bars: 50μm

**Supplementary Fig 2.**
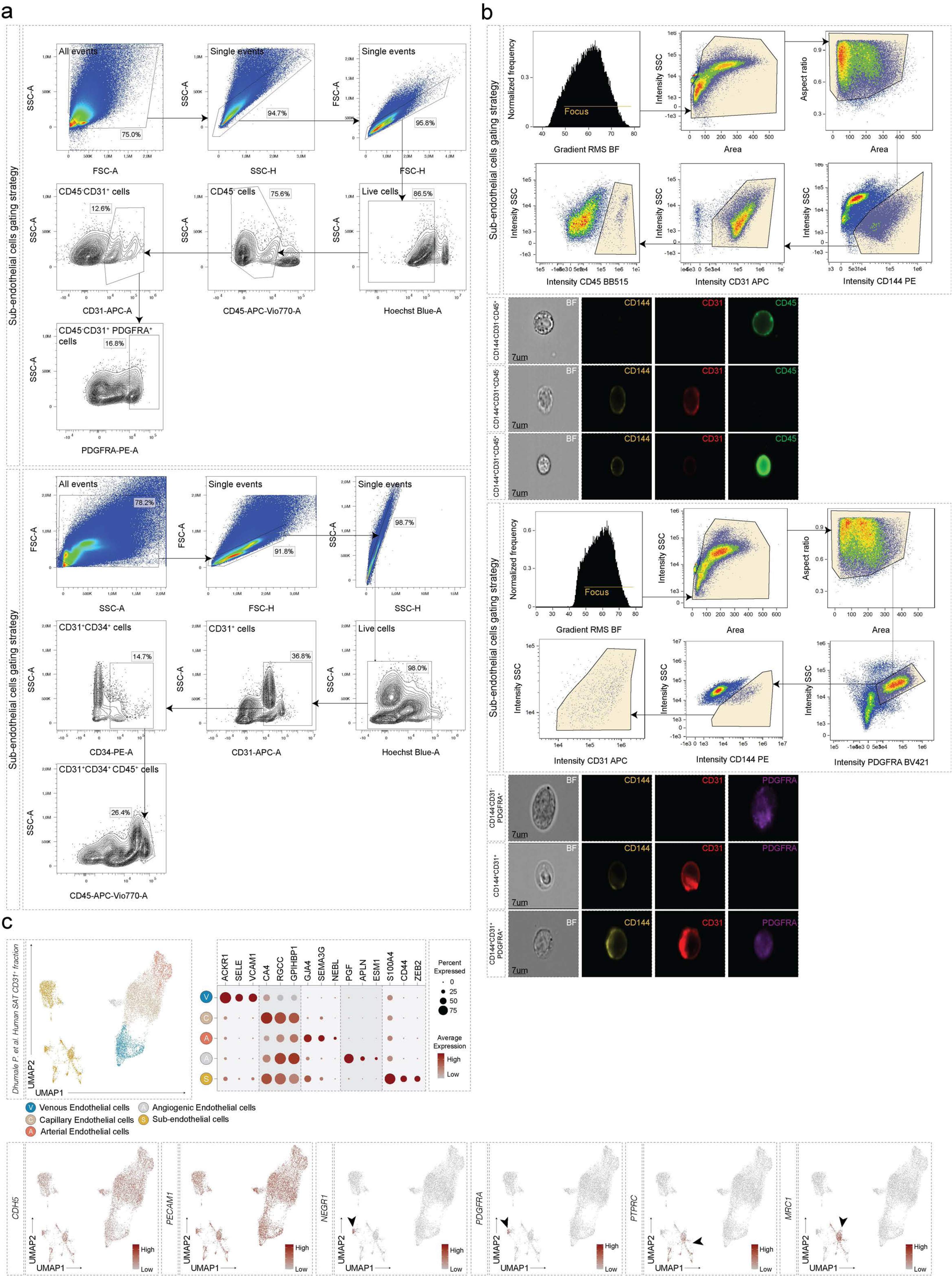
Gating strategies for conventional flow cytometry and ImageStream analyses. **a,** Scatter plots depicting the gating strategy for conventional flow cytometry of human subcutaneous adipose tissue (SAT) stromovascular fraction (SVF) cells. Cells were first gated based on size using forward and side scatter, followed by gating of live singlets. Populations of interest, CD31⁺CD34⁺CD45⁺ (*top*) and CD45⁻CD31⁺PDGFRA⁺ (*bottom*), were subsequently identified. **b**, Scatter plots showing the gating strategy for ImageStream analysis of human SAT SVF cells. In-focus cells were initially gated on side scatter and area. Singlets were selected based on aspect ratio and area, followed by identification of CD144/VE-Cadherin⁺CD31⁺CD45⁺ (*top*) and CD144/VE-Cadherin⁺CD31⁺PDGFRA⁺ (*bottom*) populations. Representative in-focus images show individual cells categorized as immune cells (CD144/VE-Cadherin⁻CD31⁻CD45⁺), fibroblasts (CD144/VE-Cadherin⁻CD31⁻PDGFRA⁺), endothelial cells (CD144/VE-Cadherin⁺CD31⁺), and sub-endothelial populations (CD144/VE-Cadherin⁺CD31⁺CD45⁺ or CD144//VE-Cadherin⁺CD31⁺PDGFRA⁺). **c**, UMAP projection of re-analyzed single-cell RNA sequencing data from human SAT SVF cells enriched for CD31⁺ cells via magnetic-activated cell sorting. Data were integrated using *Harmony* and reveal five distinct cell populations. Feature plots show expression of pan-endothelial marker genes (*CDH5*, *PECAM1*) and sub-endothelial cell markers (*NEGR1*, *PDGFRA*, *PTPRC*, *MRC1*). Color scale: red, high expression; grey, low expression. Black arrowheads denote populations with increased gene expression. Dot size: percentage of cells within each cluster expressing a given gene. Scale bar 7μm.

**Supplementary Fig 3:**
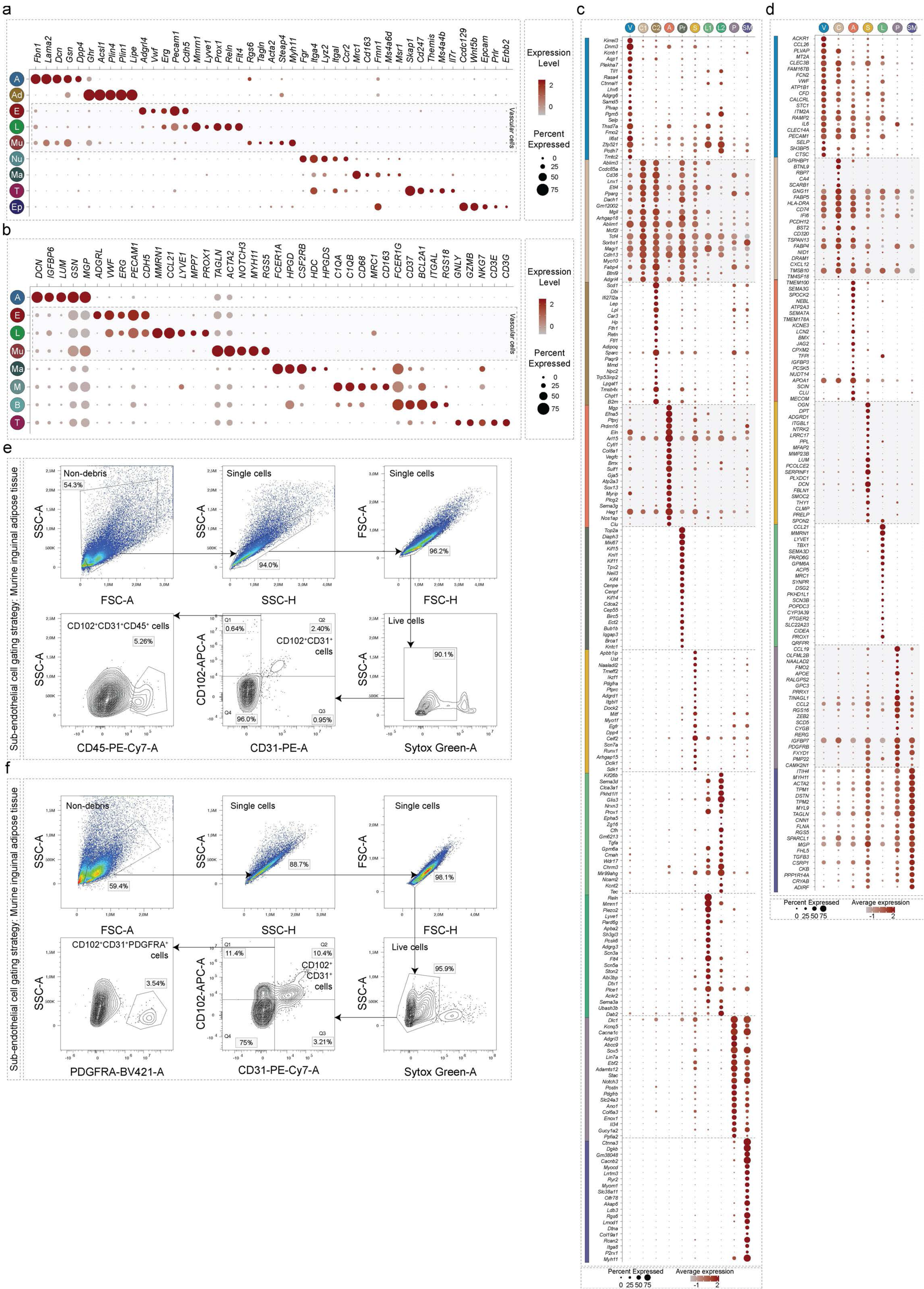
Murine inguinal adipose tissue comprise sub-endothelial cells congruent to that in humans. **a**,**b**, Marker gene expression in the major cell populations of murine inguinal adipose tissue (**a**) and porcine subcutaneous adipose tissue (**b**). **c**,**d**, Marker gene expression in the vascular cell populations of murine inguinal adipose tissue (**c**), porcine subcutaneous adipose tissue (**d**). **e,f**, Scatter plots depicting the gating strategy of murine inguinal adipose tissue SVF cells. Murine inguinal adipose tissue SVF cells were first gated based on size using the forward and side scatters and live singlets were subsequently gated to identify CD31^+^CD102^+^CD45^+^ and CD31^+^CD102^+^PDGFRA^+^ cells.

**Supplementary Fig 4:**
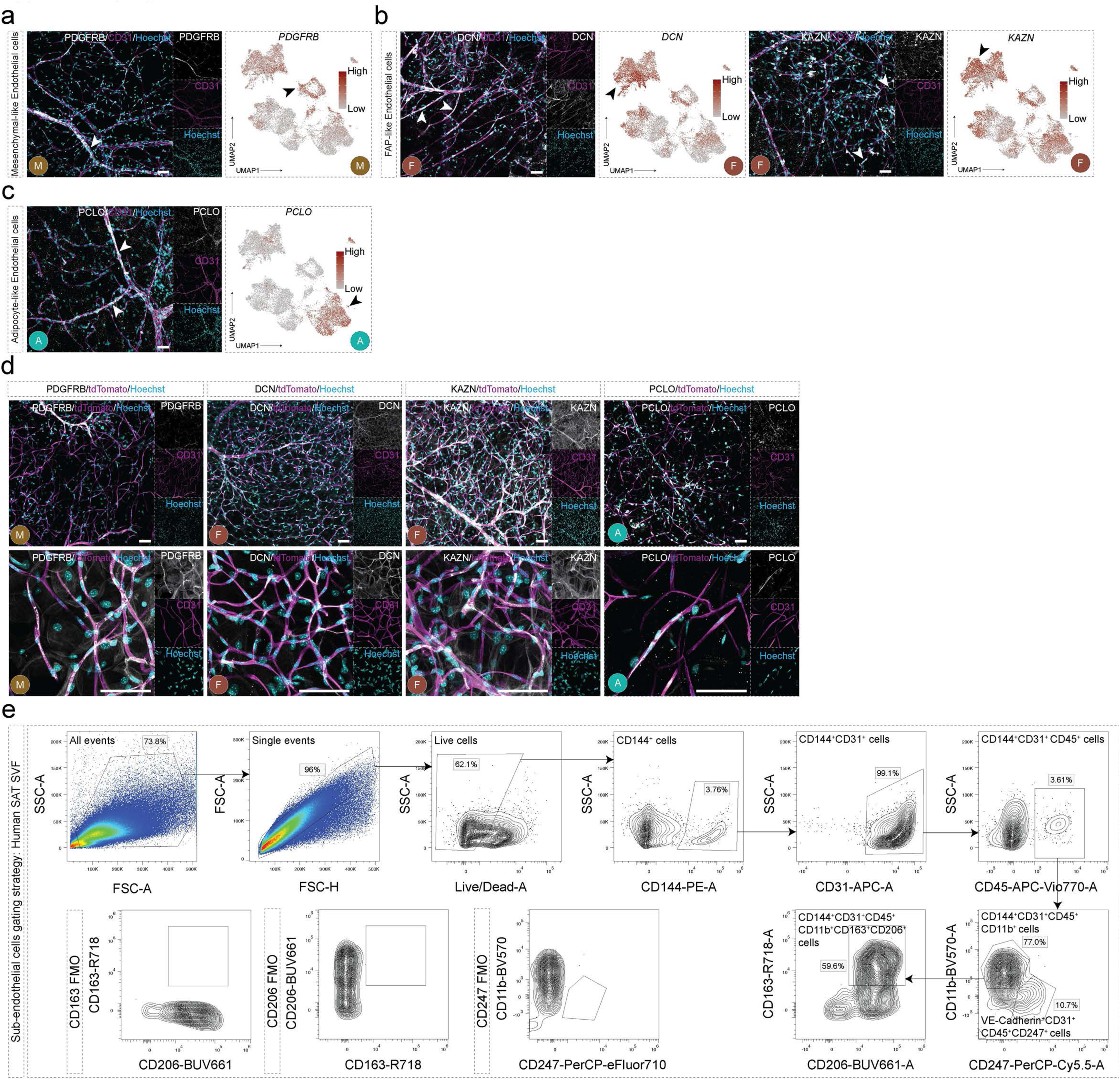
Identification of sub-endothelial cells in human and murine adipose tissue. **a-c**, *Left panels*: Immunohistochemical staining of wholemount human adipose tissue at low magnification for proteins encoded by the marker genes in the sub-AdECs sub-populations (PDGFRB, a; DCN, KAZN, b; PCLO, c). White arrowheads mark ECs positive for the target protein. *Right panels*: Feature plots depicting gene expression. Black arrowheads denote populations with increased gene expression. In all feature plots, color scale indicates gene expression level (red: high, grey: low). **d,** Immunohistochemical staining of wholemount murine adipose tissue at low and high magnification for proteins encoded by the marker genes in the sub-AdECs sub-populations (PDGFRB, DCN, KAZN, and PCLO). **e**, Human SAT stromovascular fraction cells were gated based on size using the forward and side scatters and live singlets were subsequently gated to identify CD144/VE-Cadherin^+^CD31^+^CD45^+^CD247^+^ and CD144/VE-Cadherin^+^CD31^+^CD45^+^CD11b^+^CD163^+^CD206^+^ cells. Fluorescent minus one (FMOs) were used to validate the gating strategy. Scale bars (a-d): 50μm.

**Supplementary Fig 5:**
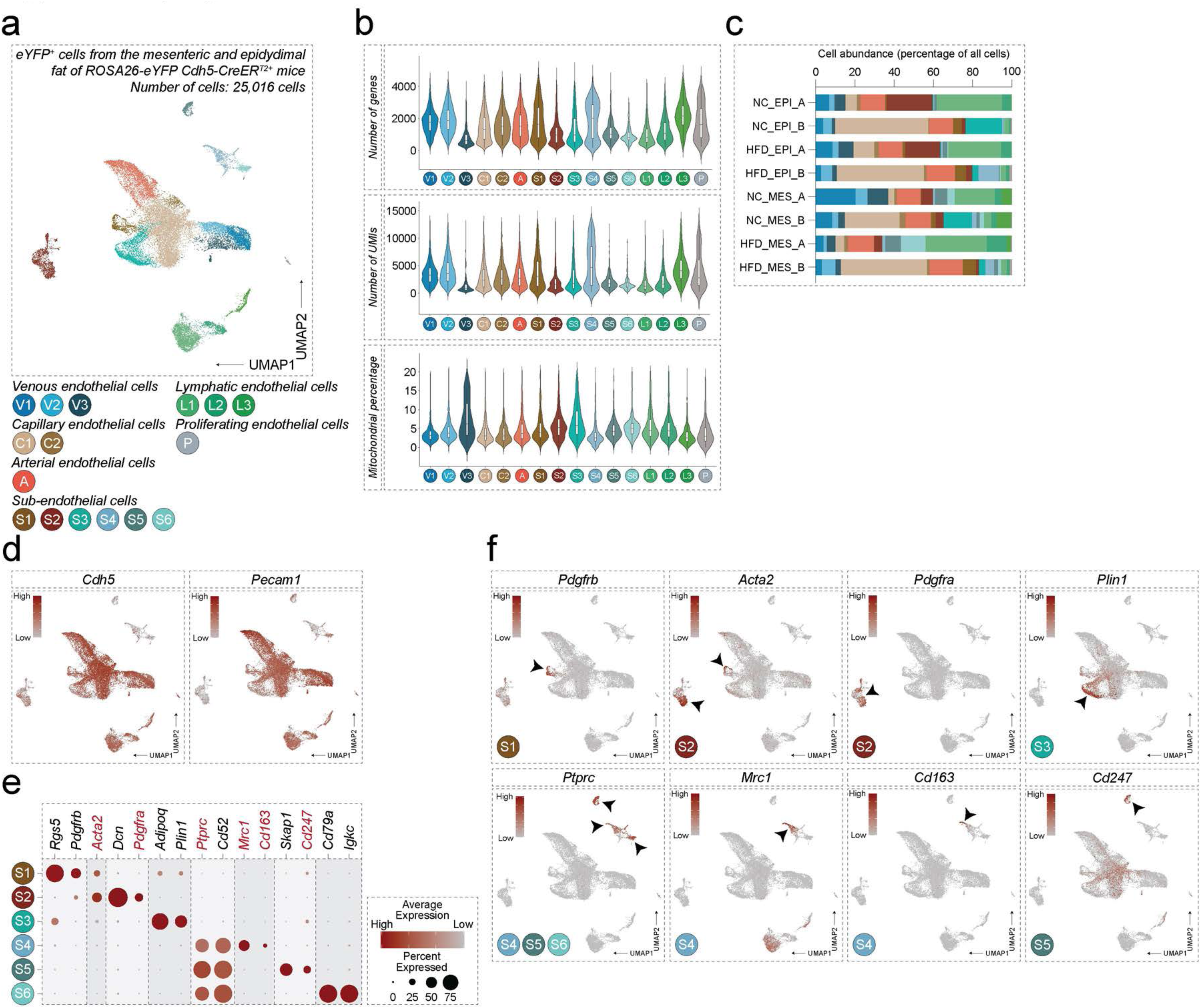
Identification of sub-endothelial cells in endothelial cell reporter mouse strains. **a**, UMAP projection of re-analyzed 25,016 eYFP+ sorted cells from the mesenteric and epidydimal fat depots of ROSA26-eYFP Cdh5:CreERT2+ mice. **b**, Violin plots depicting the quality control metrics of cells presented in (a). Number of genes, Number of UMIs, and Mitochondrial percentage. **c**, Cell abundance as a percentage of all cells sequenced for a single depot depicting the abundance of the different endothelial cell populations. **d**, Feature plots depicting the expression of the endothelial marker genes *Cdh5* and *Pecam1*. Color scale: red, high expression; grey, low expression. **e**, Marker gene expression in sub-endothelial cell populations identified in (a). Color scale: red, high expression; grey, low expression. Dot size: the percentage of cells within a cluster expressing a particular gene. **f**, Feature plots depicting the expression of selected sub-endothelial cell marker genes. Color scale: red, high expression; grey, low expression. Black arrowheads denote populations with increased gene expression.

**Supplementary Fig 6:**
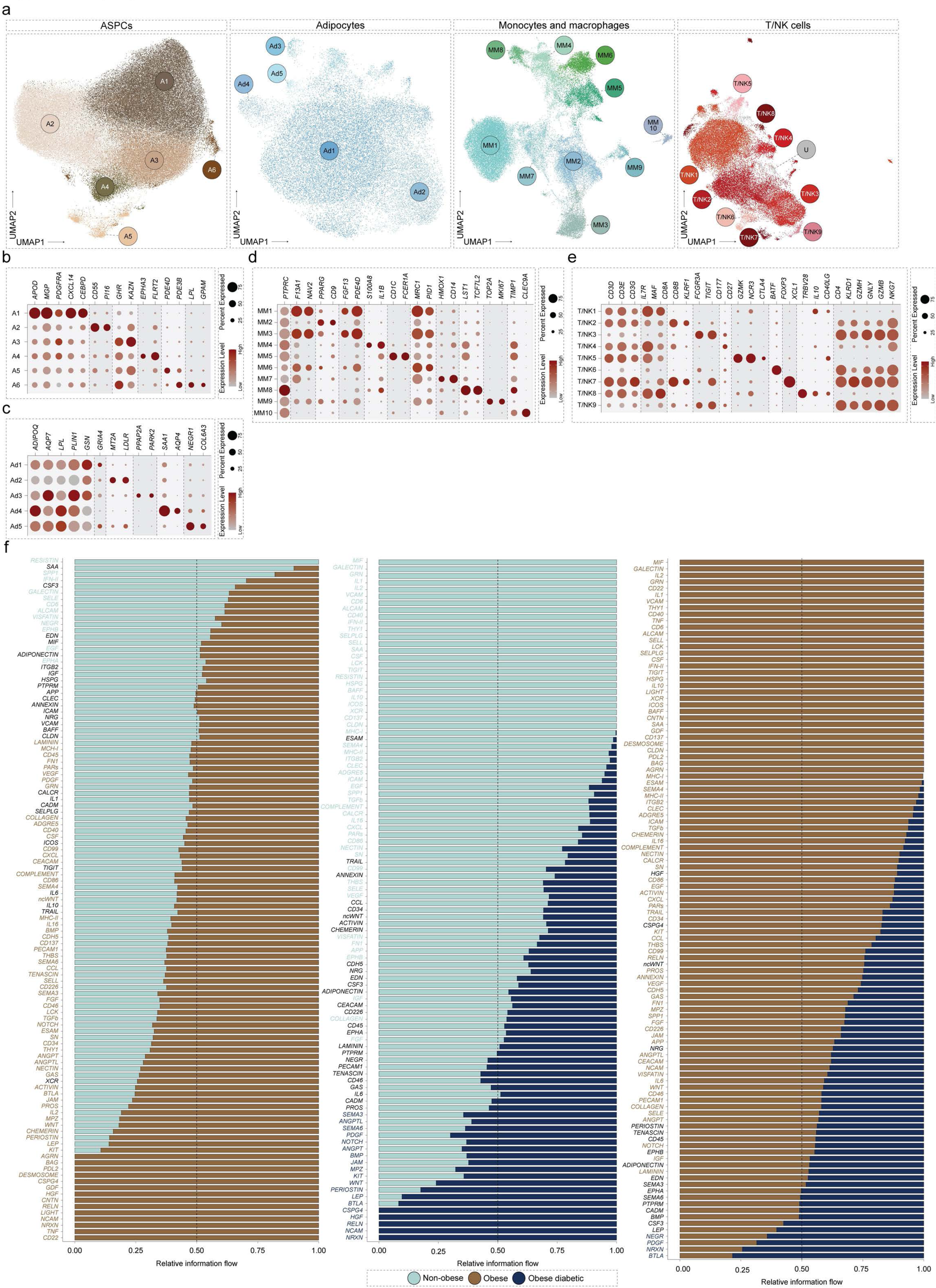
Altered intercellular interactions in human SAT in obesity and obesity compounded by T2D. **a**, UMAP representation of subclusters comprising major cell populations within the human SAT including ASPCs, Adipocytes, Macrophages and Monocytes, and T/NK cells, clustered at high granularity. **b-e**, Marker gene expression in each cell population identified in (**a**). Color scale: red, high expression; grey, low expression. Dot size: the percentage of cells within a cluster expressing a particular gene. **f**, Bar plots showing the relative information flow for significant communication pathways in the three comparisons; Obese vs. Non-obese (*left panel*), Obese diabetic vs. Non-obese (*middle panel*), and Obese diabetic vs. Obese (*right panel*). The y-axis represents the significantly altered pathways. Colored pathways are enriched in a specific condition for a given comparison. The x-axis represents the relative information flow of a given pathway.

**Supplementary Fig 7:**
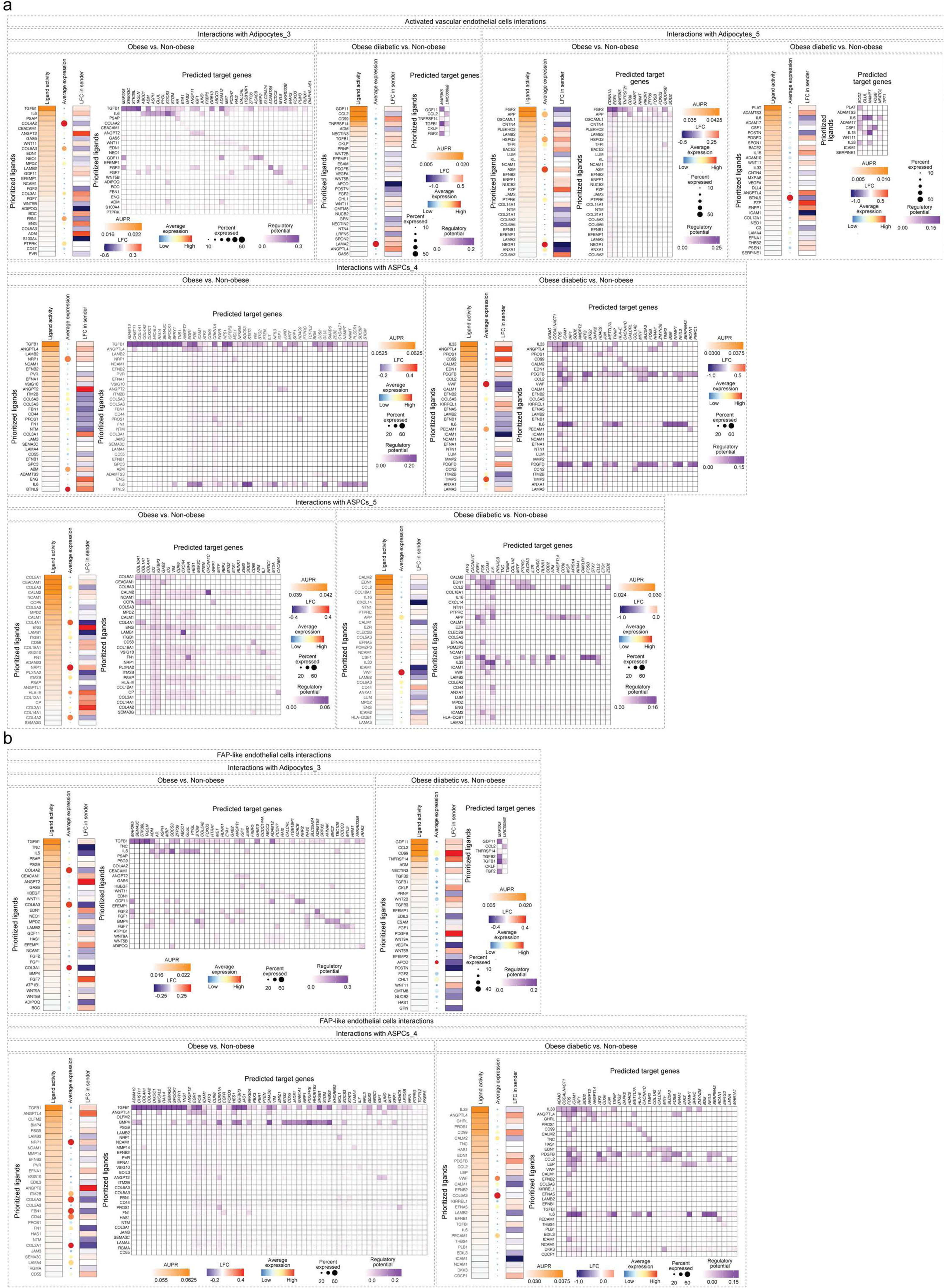
Vascular cell-mediated (Activated vascular endothelial cells and FAP-like endothelial cells) intracellular signaling in receiving cells. **a,b**, Heatmaps depicting prioritized ligands ordered by ligand activity, average expression, and log fold change in sending cells (Activated vascular endothelial cells in **a** and FAP-like endothelial cells in **b**) along with heatmaps depicting the regulatory potential of the prioritized ligands on predicted target genes. AUPR, Area under the precision-recall curve; LFC, log fold change. Average expression color scale: red, high expression; blue, low expression. Dot size: the percentage of cells within a given sending cell cluster expressing a particular gene.

**Supplementary Fig 8:**
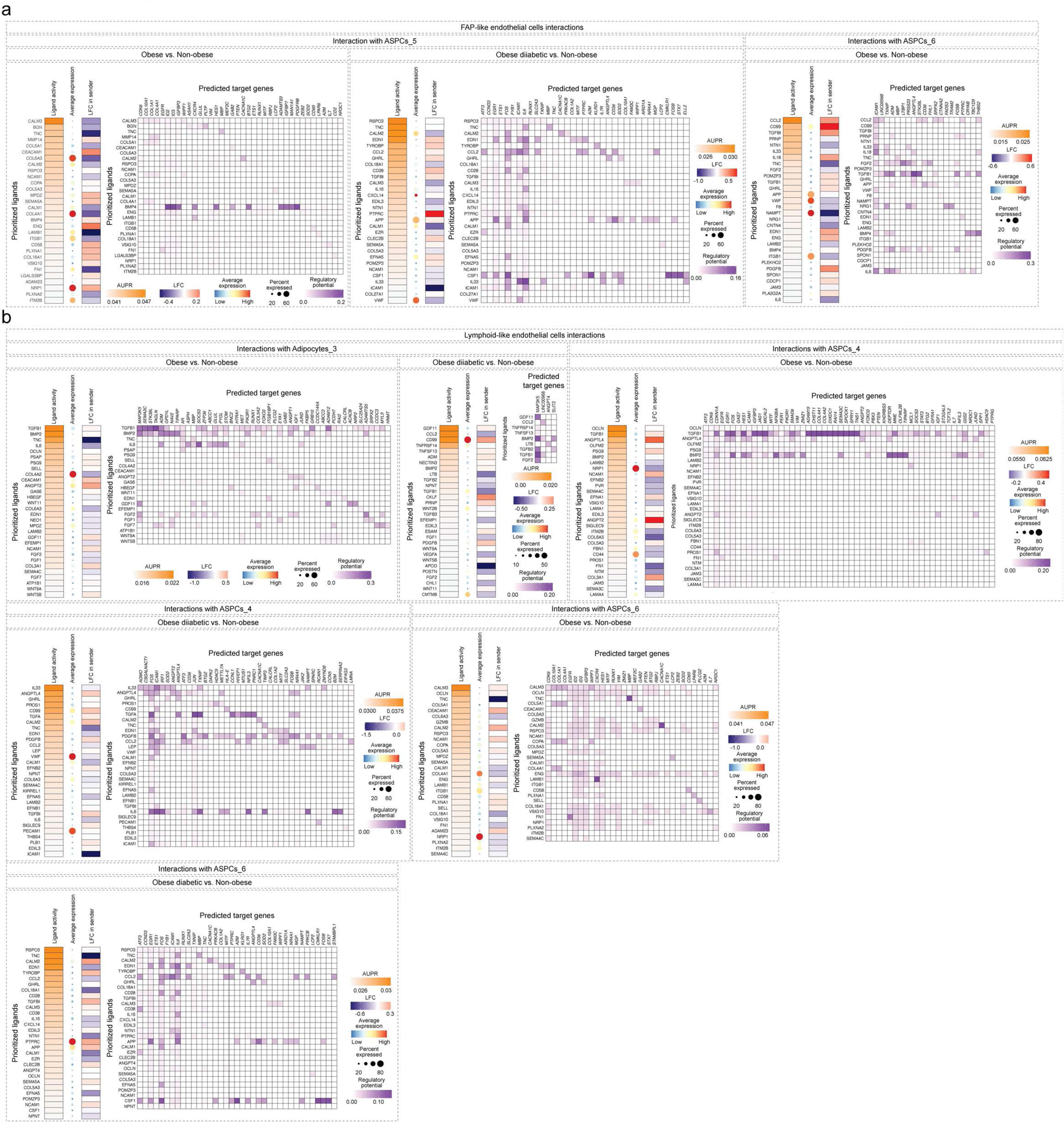
Vascular cell-mediated (FAP-like endothelial cells and lymphoid-like endothelial cells) intracellular signaling in receiving cells. **a,b**, Heatmaps depicting prioritized ligands ordered by ligand activity, average expression, and log fold change in sending cells (FAP-like endothelial cells in **a** and Lymphoid-like endothelial cells in **b**) along with heatmaps depicting the regulatory potential of the prioritized ligands on predicted target genes. AUPR, Area under the precision-recall curve; LFC, log fold change. Average expression color scale: red, high expression; blue, low expression. Dot size: the percentage of cells within a given sending cell cluster expressing a particular gene.

## References

1 AlZaim, I., de Rooij, L. P., Sheikh, B. N., Börgeson, E. & Kalucka, J. The evolving functions of the vasculature in regulating adipose tissue biology in health and obesity. Nature Reviews Endocrinology, 1–17 (2023).

2 Merrick, D. et al. Identification of a mesenchymal progenitor cell hierarchy in adipose tissue. Science 364, eaav2501 (2019).

3 Vijay, J. et al. Single-cell analysis of human adipose tissue identifies depot-and disease-specific cell types. Nature metabolism 2, 97–109 (2020).

4 Hildreth, A. D. et al. Single-cell sequencing of human white adipose tissue identifies new cell states in health and obesity. Nature immunology 22, 639–653 (2021).

5 Consortium*, T. S., et al. The Tabula Sapiens: A multiple-organ, single-cell transcriptomic atlas of humans. Science 376, eabl4896 (2022).

6 Barboza, C. S. et al. Single-nuclei transcriptome of human adipose tissue reveals metabolically distinct depot-specific adipose progenitor subpopulations. bioRxiv, 2022.2006. 2029.496888 (2022).

7 Emont, M. P. et al. A single-cell atlas of human and mouse white adipose tissue. Nature 603, 926–933 (2022).

8 Massier, L. et al. An integrated single cell and spatial transcriptomic map of human white adipose tissue. Nature Communications 14, 1438 (2023).

9 Maniyadath, B., Zhang, Q., Gupta, R. K. & Mandrup, S. Adipose tissue at single-cell resolution. Cell Metabolism 35, 386–413 (2023).

10 Thiriot, A. et al. Differential DARC/ACKR1 expression distinguishes venular from non-venular endothelial cells in murine tissues. BMC biology 15, 1–19 (2017).

11 Kalucka, J. et al. Single-cell transcriptome atlas of murine endothelial cells. Cell 180, 764–779. e720 (2020).

12 Li, J. et al. Neurotensin is an anti-thermogenic peptide produced by lymphatic endothelial cells. Cell metabolism 33, 1449–1465. e1446 (2021).

13 Hasan, S. S. et al. Obesity drives depot-specific vascular remodeling in male white adipose tissue. Nature Communications 16, 5392 (2025).

14 Su, T. et al. Single-cell analysis of early progenitor cells that build coronary arteries. Nature 559, 356–362 (2018).

15 Chavkin, N. W. et al. Obesity accelerates endothelial-to-mesenchymal transition in adipose tissues of mice and humans. Frontiers in Cardiovascular Medicine 10, 1264479 (2023).

16 Dhumale, P. et al. CD31 defines a subpopulation of human adipose-derived regenerative cells with potent angiogenic effects. Scientific Reports 13, 14401 (2023).

17 Bäckdahl, J. et al. Spatial mapping reveals human adipocyte subpopulations with distinct sensitivities to insulin. Cell metabolism 33, 1869–1882. e1866 (2021).

18 Shao, X. et al. MatrisomeDB 2.0: 2023 updates to the ECM-protein knowledge database. Nucleic Acids Research 51, D1519–D1530 (2023).

19 van Splunder, H., Villacampa, P., Martínez-Romero, A. & Graupera, M. Pericytes in the disease spotlight. Trends in Cell Biology (2023).

20 Wilhelm, K. et al. FOXO1 couples metabolic activity and growth state in the vascular endothelium. Nature 529, 216–220 (2016). 10.1038/nature16498

21 Trimm, E. & Red-Horse, K. Vascular endothelial cell development and diversity. Nature Reviews Cardiology 20, 197–210 (2023).

22 Jeong, H.-W. et al. Transcriptional regulation of endothelial cell behavior during sprouting angiogenesis. Nature communications 8, 726 (2017).

23 Kakogiannos, N. et al. JAM-A Acts via C/EBP-α to Promote Claudin-5 Expression and Enhance Endothelial Barrier Function. Circ Res 127, 1056–1073 (2020). 10.1161/circresaha.120.316742

24 Reyes-Palomares, A. et al. Remodeling of active endothelial enhancers is associated with aberrant gene-regulatory networks in pulmonary arterial hypertension. Nature communications 11, 1673 (2020).

25 Hoshino, A. et al. Gain-of-function IKZF1 variants in humans cause immune dysregulation associated with abnormal T/B cell late differentiation. Sci Immunol 7, eabi7160 (2022). 10.1126/sciimmunol.abi7160

26 Goh, W. et al. IKAROS and AIOLOS directly regulate AP-1 transcriptional complexes and are essential for NK cell development. Nature Immunology, 1–16 (2024).

27 Boutboul, D. et al. Dominant-negative IKZF1 mutations cause a T, B, and myeloid cell combined immunodeficiency. The Journal of clinical investigation 128, 3071–3087 (2018).

28 Lv, L., Meng, Q., Ye, M., Wang, P. & Xue, G. STAT4 deficiency protects against neointima formation following arterial injury in mice. Journal of Molecular and Cellular Cardiology 74, 284–294 (2014).

29 Mognol, G. P., Carneiro, F. R. G., Robbs, B. K., Faget, D. V. & Viola, J. P. d. B. Cell cycle and apoptosis regulation by NFAT transcription factors: new roles for an old player. Cell death & disease 7, e2199–e2199 (2016).

30 Wang, F. et al. Endothelial cell heterogeneity and microglia regulons revealed by a pig cell landscape at single-cell level. Nature communications 13, 3620 (2022).

31 Bondareva, O. et al. Single-cell profiling of vascular endothelial cells reveals progressive organ-specific vulnerabilities during obesity. Nature Metabolism 4, 1591–1610 (2022).

32 Naba, A. Mechanisms of assembly and remodelling of the extracellular matrix. Nature Reviews Molecular Cell Biology, 1–21 (2024).

33 Arbee, S. et al. Versican Maintains the Homeostasis of Adipose Tissues and Regulates Energy Metabolism. Biochemical and Biophysical Research Communications, 150309 (2024).

34 Han, C. Y. et al. Adipocyte-derived versican and macrophage-derived biglycan control adipose tissue inflammation in obesity. Cell reports 31 (2020).

35 AlZaim, I., de Rooij, L. P., Sheikh, B. N., Börgeson, E. & Kalucka, J. The evolving functions of the vasculature in regulating adipose tissue biology in health and obesity. Nature Reviews Endocrinology 19, 691–707 (2023).

36 Armulik, A., Genové, G. & Betsholtz, C. Pericytes: developmental, physiological, and pathological perspectives, problems, and promises. Developmental cell 21, 193–215 (2011).

37 Holm, A., Heumann, T. & Augustin, H. G. Microvascular mural cell organotypic heterogeneity and functional plasticity. Trends in cell biology 28, 302–316 (2018).

38 Ahmad, R., Rah, B., Bastola, D., Dhawan, P. & Singh, A. B. Obesity-induces Organ and Tissue Specific Tight Junction Restructuring and Barrier Deregulation by Claudin Switching. Sci Rep 7, 5125 (2017). 10.1038/s41598-017-04989-8

39 Feng, Z., Fang, C., Ma, Y. & Chang, J. Obesity-induced blood-brain barrier dysfunction: phenotypes and mechanisms. J Neuroinflammation 21, 110 (2024). 10.1186/s12974-024-03104-9

40 Basatemur, G. L., Jørgensen, H. F., Clarke, M. C., Bennett, M. R. & Mallat, Z. Vascular smooth muscle cells in atherosclerosis. Nature reviews cardiology 16, 727–744 (2019).

41 Liu, Y. et al. Shear stress activation of SREBP1 in endothelial cells is mediated by integrins. Arteriosclerosis, thrombosis, and vascular biology 22, 76–81 (2002).

42 Shimano, H. et al. Sterol regulatory element-binding protein-1 as a key transcription factor for nutritional induction of lipogenic enzyme genes. Journal of Biological Chemistry 274, 35832–35839 (1999).

43 Ong, Y. T. et al. A YAP/TAZ-TEAD signalling module links endothelial nutrient acquisition to angiogenic growth. Nature metabolism 4, 672–682 (2022).

44 Cai, X., Wang, K. C. & Meng, Z. Mechanoregulation of YAP and TAZ in Cellular Homeostasis and Disease Progression. Front Cell Dev Biol 9, 673599 (2021). 10.3389/fcell.2021.673599

45 Taylor, J. et al. Endothelial Notch1 signaling in white adipose tissue promotes cancer cachexia. Nature Cancer 4, 1544–1560 (2023).

46 Franco, C. A. et al. Serum response factor is required for sprouting angiogenesis and vascular integrity. Developmental cell 15, 448–461 (2008).

47 Weinl, C. et al. Endothelial SRF/MRTF ablation causes vascular disease phenotypes in murine retinae. The Journal of clinical investigation 123, 2193–2206 (2013).

48 Andueza, A. et al. Endothelial reprogramming by disturbed flow revealed by single-cell RNA and chromatin accessibility study. Cell reports 33 (2020).

49 Loft, A. et al. Towards a consensus atlas of human and mouse adipose tissue at single-cell resolution. Nat Metab 7, 875–894 (2025). 10.1038/s42255-025-01296-9

50 Tang, W. et al. White fat progenitor cells reside in the adipose vasculature. Science 322, 583–586 (2008).

51 Tran, K.-V. et al. The vascular endothelium of the adipose tissue gives rise to both white and brown fat cells. Cell metabolism 15, 222–229 (2012).

52 Cattaneo, P. et al. Parallel lineage-tracing studies establish fibroblasts as the prevailing in vivo adipocyte progenitor. Cell reports 30, 571–582. e572 (2020).

53 Ioannidou, A. et al. Hypertrophied human adipocyte spheroids as in vitro model of weight gain and adipose tissue dysfunction. The Journal of physiology 600, 869–883 (2022).

54 Amersfoort, J., Eelen, G. & Carmeliet, P. Immunomodulation by endothelial cells—partnering up with the immune system? Nature Reviews Immunology 22, 576–588 (2022).

55 Yang Loureiro, Z., et al. Wnt signaling preserves progenitor cell multipotency during adipose tissue development. Nature Metabolism, 1–15 (2023).

56 Palani, N. P. et al. Adipogenic and SWAT cells separate from a common progenitor in human brown and white adipose depots. Nature metabolism, 1–18 (2023).

57 Xiong, J. et al. A metabolic basis for endothelial-to-mesenchymal transition. Molecular cell 69, 689–698. e687 (2018).

58 Xu, Y. & Kovacic, J. C. Endothelial to mesenchymal transition in health and disease. Annual Review of Physiology 85, 245–267 (2023).

59 Dunaway, L. S. et al. Obesogenic diet disrupts tissue-specific mitochondrial gene signatures in the artery and capillary endothelium. Physiological Genomics 56, 113–127 (2024).

60 Dheedene, W. et al. Loss of endothelial ZEB2 in mice attenuates steatosis early during metabolic dysfunction-associated steatotic liver disease. Scientific Reports 15, 23434 (2025).

61 Tardajos-Ayllon, B. et al. TWIST1 drives endothelial-to-mesenchymal-transition to stabilize atherosclerotic plaques. bioRxiv, 2025.2005.2019.654847 (2025). 10.1101/2025.05.19.654847

62 Ren, J. et al. Single-cell sequencing reveals that endothelial cells, EndMT cells and mural cells contribute to the pathogenesis of cavernous malformations. Experimental & Molecular Medicine 55, 628–642 (2023).

63 Pierce, A. D. et al. Glucose-activated RUNX2 phosphorylation promotes endothelial cell proliferation and an angiogenic phenotype. Journal of cellular biochemistry 113, 282–292 (2012).

64 Mochin, M. T. et al. Hyperglycemia and redox status regulate RUNX2 DNA-binding and an angiogenic phenotype in endothelial cells. Microvascular Research 97, 55–64 (2015).

65 Monelli, E. et al. Angiocrine polyamine production regulates adiposity. Nat Metab 4, 327–343 (2022). 10.1038/s42255-022-00544-6

66 Festa, J., AlZaim, I. & Kalucka, J. Adipose tissue endothelial cells: insights into their heterogeneity and functional diversity. Current Opinion in Genetics & Development 81, 102055 (2023).

67 Jin, S. et al. Inference and analysis of cell-cell communication using CellChat. Nature communications 12, 1088 (2021).

68 Dimitrov, D. et al. LIANA+ provides an all-in-one framework for cell–cell communication inference. Nature Cell Biology, 1–10 (2024).

69 Herold, J. & Kalucka, J. Angiogenesis in adipose tissue: the interplay between adipose and endothelial cells. Frontiers in physiology 11, 624903 (2021).

70 Cao, Y. Angiogenesis modulates adipogenesis and obesity. The Journal of clinical investigation 117, 2362–2368 (2007).

71 Browaeys, R., Saelens, W. & Saeys, Y. NicheNet: modeling intercellular communication by linking ligands to target genes. Nature methods 17, 159–162 (2020).

72. Inouye, K. E., et al. Endothelial-derived FABP4 constitutes the majority of basal circulating hormone and regulates lipolysis-driven insulin secretion. JCI insight 8 (2023).

73 Ke, K. et al. Increased expression of CD74 in atherosclerosis associated with inflammatory responses of endothelial cells and macrophages. Biochemical Genetics, 1–17 (2023).

74 Rodor, J. et al. Single-cell RNA sequencing profiling of mouse endothelial cells in response to pulmonary arterial hypertension. Cardiovascular Research 118, 2519–2534 (2022).

75 Goveia, J. et al. An integrated gene expression landscape profiling approach to identify lung tumor endothelial cell heterogeneity and angiogenic candidates. Cancer cell 37, 21–36. e13 (2020).

76 Wälchli, T. et al. Single-cell atlas of the human brain vasculature across development, adulthood and disease. Nature, 1–11 (2024).

77. Zhang, L., et al. Single-cell transcriptomic profiling of lung endothelial cells identifies dynamic inflammatory and regenerative subpopulations. JCI insight 7 (2022).

78 Suur, B. E. et al. Therapeutic potential of the Proprotein Convertase Subtilisin/Kexin family in vascular disease. Frontiers in Pharmacology 13, 988561 (2022).

79 Kanemaru, K. et al. Spatially resolved multiomics of human cardiac niches. Nature 619, 801–810 (2023).

80 Sakaue, S. & Okada, Y. GREP: genome for REPositioning drugs. Bioinformatics 35, 3821–3823 (2019).

81 Suttorp, N., Weber, U., Welsch, T. & Schudt, C. Role of phosphodiesterases in the regulation of endothelial permeability in vitro. The Journal of clinical investigation 91, 1421–1428 (1993).

82 MacKeil, J. L. et al. Phosphodiesterase 3B (PDE3B) antagonizes the anti-angiogenic actions of PKA in human and murine endothelial cells. Cellular Signalling 62, 109342 (2019).

83 Zhu, Z., Chambers, S. & Bhatia, M. Substance P Promotes Leukocyte Infiltration in the Liver and Lungs of Mice with Sepsis: A Key Role for Adhesion Molecules on Vascular Endothelial Cells. International Journal of Molecular Sciences 25, 6500 (2024).

84 Bhatia, M. et al. Role of substance P and the neurokinin 1 receptor in acute pancreatitis and pancreatitis-associated lung injury. Proceedings of the National Academy of Sciences 95, 4760–4765 (1998).

85 Li, M. et al. Potential therapeutic effect of NK1R antagonist in diabetic non-healing wound and depression. Frontiers in Endocrinology 13, 1077514 (2023).

86 Zhang, M. J. et al. Polygenic enrichment distinguishes disease associations of individual cells in single-cell RNA-seq data. Nature genetics 54, 1572–1580 (2022).

87. Whytock, K. L., et al. Single cell full-length transcriptome of human subcutaneous adipose tissue reveals unique and heterogeneous cell populations. Iscience 25 (2022).

88 Zhang, Y., Kang, Z., Liu, M., Wang, L. & Liu, F. Single-cell omics identifies inflammatory signaling as a trans-differentiation trigger in mouse embryos. Developmental Cell (2024).

89. Nasim, S., et al. CD45 Is Sufficient to Initiate Endothelial-to-Mesenchymal Transition in Human Endothelial Cells—Brief Report. Arteriosclerosis, Thrombosis, and Vascular Biology 43, e124-e131 (2023).

90 Yamashiro, Y. et al. Partial endothelial-to-mesenchymal transition mediated by HIF-induced CD45 in neointima formation upon carotid artery ligation. Cardiovascular Research 119, 1606–1618 (2023).

91 Min, S. Y. et al. Human’brite/beige’adipocytes develop from capillary networks, and their implantation improves metabolic homeostasis in mice. Nature medicine 22, 312–318 (2016).

92 Herold, J. & Kalucka, J. Angiogenesis in Adipose Tissue: The Interplay Between Adipose and Endothelial Cells. Front Physiol 11, 624903 (2020). 10.3389/fphys.2020.624903

93 Herold, J. & Kalucka, J. in Angiogenesis 235–250 (Springer, 2022).

94 Frankish, A. et al. GENCODE reference annotation for the human and mouse genomes. Nucleic acids research 47, D766–D773 (2019).

95 Fleming, S. J. et al. Unsupervised removal of systematic background noise from droplet-based single-cell experiments using CellBender. Nature methods 20, 1323–1335 (2023).

96 McGinnis, C. S., Murrow, L. M. & Gartner, Z. J. DoubletFinder: doublet detection in single-cell RNA sequencing data using artificial nearest neighbors. Cell systems 8, 329–337. e324 (2019).

97 Korsunsky, I. et al. Fast, sensitive and accurate integration of single-cell data with Harmony. Nature methods 16, 1289–1296 (2019).

98 Lopez, R., Regier, J., Cole, M. B., Jordan, M. I. & Yosef, N. Deep generative modeling for single-cell transcriptomics. Nature methods 15, 1053–1058 (2018).

99 Polański, K. et al. BBKNN: fast batch alignment of single cell transcriptomes. Bioinformatics 36, 964–965 (2020).

100 Zappia, L. & Oshlack, A. Clustering trees: a visualization for evaluating clusterings at multiple resolutions. Gigascience 7, giy083 (2018).

101 Büttner, M., Miao, Z., Wolf, F. A., Teichmann, S. A. & Theis, F. J. A test metric for assessing single-cell RNA-seq batch correction. Nature methods 16, 43–49 (2019).

102 Azzalini, A. & Menardi, G. Clustering via nonparametric density estimation: The R package pdfCluster. *arXiv preprint arXiv:1301*.6559 (2013).

103 Luecken, M. D. et al. Benchmarking atlas-level data integration in single-cell genomics. Nature methods 19, 41–50 (2022).

104 Buettner, M., Ostner, J., Mueller, C. L., Theis, F. J. & Schubert, B. scCODA is a Bayesian model for compositional single-cell data analysis. Nature communications 12, 6876 (2021).

105 Hänzelmann, S., Castelo, R. & Guinney, J. GSVA: gene set variation analysis for microarray and RNA-seq data. BMC bioinformatics 14, 1–15 (2013).

106 Andreatta, M. & Carmona, S. J. UCell: Robust and scalable single-cell gene signature scoring. Computational and structural biotechnology journal 19, 3796–3798 (2021).

107 Schubert, M. et al. Perturbation-response genes reveal signaling footprints in cancer gene expression. Nature communications 9, 20 (2018).

108 Wu, T. et al. clusterProfiler 4.0: A universal enrichment tool for interpreting omics data. The innovation 2 (2021).

109 Aibar, S. et al. SCENIC: single-cell regulatory network inference and clustering. Nature methods 14, 1083–1086 (2017).

110 Badia-i-Mompel, P. et al. decoupleR: ensemble of computational methods to infer biological activities from omics data. Bioinformatics Advances 2, vbac016 (2022).

111 Wingender, E., Schoeps, T., Haubrock, M. & Dönitz, J. TFClass: a classification of human transcription factors and their rodent orthologs. Nucleic acids research 43, D97–D102 (2015).

112 Van de Sande, B. et al. A scalable SCENIC workflow for single-cell gene regulatory network analysis. Nature protocols 15, 2247–2276 (2020).

113 Moerman, T. et al. GRNBoost2 and Arboreto: efficient and scalable inference of gene regulatory networks. Bioinformatics 35, 2159–2161 (2019).

114 Kiselev, V. Y., Yiu, A. & Hemberg, M. scmap: projection of single-cell RNA-seq data across data sets. Nature methods 15, 359–362 (2018).

115 Durinck, S., Spellman, P. T., Birney, E. & Huber, W. Mapping identifiers for the integration of genomic datasets with the R/Bioconductor package biomaRt. Nature protocols 4, 1184–1191 (2009).

116 Hrovatin, K. et al. Integrating single-cell RNA-seq datasets with substantial batch effects. bioRxiv, 2023.2011. 2003.565463 (2024).

117 Malagoli Tagliazucchi, G., Wiecek, A. J., Withnell, E. & Secrier, M. Genomic and microenvironmental heterogeneity shaping epithelial-to-mesenchymal trajectories in cancer. Nature communications 14, 789 (2023).

118 Cao, J. et al. The single-cell transcriptional landscape of mammalian organogenesis. Nature 566, 496–502 (2019).

119 Kang, M. et al. Mapping single-cell developmental potential in health and disease with interpretable deep learning. bioRxiv, 2024.2003. 2019.585637 (2024).

120 Guo, M., Bao, E. L., Wagner, M., Whitsett, J. A. & Xu, Y. SLICE: determining cell differentiation and lineage based on single cell entropy. Nucleic acids research 45, e54–e54 (2017).

121 Wolf, F. A. et al. PAGA: graph abstraction reconciles clustering with trajectory inference through a topology preserving map of single cells. Genome biology 20, 1–9 (2019).

122 Wolf, F. A., Angerer, P. & Theis, F. J. SCANPY: large-scale single-cell gene expression data analysis. Genome biology 19, 1–5 (2018).

123 Setty, M. et al. Characterization of cell fate probabilities in single-cell data with Palantir. Nature biotechnology 37, 451–460 (2019).

124 Street, K. et al. Slingshot: cell lineage and pseudotime inference for single-cell transcriptomics. BMC genomics 19, 1–16 (2018).

125 Lange, M. et al. CellRank for directed single-cell fate mapping. Nature methods 19, 159–170 (2022).

126 La Manno, G. et al. RNA velocity of single cells. Nature 560, 494–498 (2018).

127 Bergen, V., Lange, M., Peidli, S., Wolf, F. A. & Theis, F. J. Generalizing RNA velocity to transient cell states through dynamical modeling. Nature biotechnology 38, 1408–1414 (2020).

128 Kamimoto, K. et al. Dissecting cell identity via network inference and in silico gene perturbation. Nature 614, 742–751 (2023).

129 Finan, C. et al. The druggable genome and support for target identification and validation in drug development. Science translational medicine 9, eaag1166 (2017).

130 Kleshchevnikov, V. et al. Cell2location maps fine-grained cell types in spatial transcriptomics. Nature biotechnology 40, 661–671 (2022).

131 Bak, R. O., Dever, D. P. & Porteus, M. H. CRISPR/Cas9 genome editing in human hematopoietic stem cells. Nature protocols 13, 358–376 (2018).

132 Galarraga, M. et al. Adiposoft: automated software for the analysis of white adipose tissue cellularity in histological sections. Journal of lipid research 53, 2791–2796 (2012).

133 Kristensen, M. W. et al. Vol. 99 265–268 (Wiley Online Library, 2021).

134 Emmaneel, A. et al. PeacoQC: Peak-based selection of high quality cytometry data. Cytometry Part A 101, 325–338 (2022).

